# Mapping the genomic and transcriptomic features associated with telomere maintenance mechanisms in complex karyotype sarcomas

**DOI:** 10.64898/2026.05.17.725728

**Authors:** Nicola Biondi, Maria Luisa Ratto, Rajesh Pal, Tobias Rausch, Stefanie Stirl, Laura Villacorta, Armin Hadzic, Caroline Knotz, Lina Sieverling, Miguel Gonzalez Woge, Katrin Pfütze, Christina Georg, Cihan Erkut, Umut Toprak, Sabine Stainczyk, Maria-Veronica Teleanu, Simon Kreutzfeldt, Peter Horak, Christoph Heining, Daniel Hübschmann, Bernd Kasper, Peter Hohenberger, Klaus Schulze-Osthoff, Ulrich Keilholz, Damian Rieke Lang, Alisa Lörsch, Nicole Pfarr, Thomas Kindler, Christian H. Brandts, Melanie Boerres, Patrick Metzger, Frederik Klauschen, Sebastian Bauer, Hanno Glimm, Stefan Fröhling, Claudia Scholl, Frank Westermann, Karsten Rippe, Vladimir Benes, Isidro Cortes-Ciriano, Jan O. Korbel, Benedikt Brors, Lars Feuerbach, Priya Chudasama

**Affiliations:** Precision Sarcoma Research Group, German Cancer Research Center (DKFZ), Heidelberg, Germany; Genomics Core Facility, European Molecular Biology Laboratory (EMBL), 69117 Heidelberg, Germany; Chromatin Networks, German Cancer Research Center (DKFZ) and BioQuant, Heidelberg, Germany; Division of Applied Bioinformatics, German Cancer Research Center, German Cancer Consortium, Heidelberg, Germany; Faculty of Biosciences, Heidelberg University, 69120, Heidelberg, Germany; Department of General Surgery ABC Medical Center, Mexico City, Mexico; Division of Graduate Studies, Faculty of Medicine, Universisad Nacional Autonoma de Mexico, Mexico City, Mexico; NCT Sample Processing Laboratory, German Cancer Research Center (DKFZ) and National Center for Tumor Diseases (NCT), NCT Heidelberg, A Partnership Between DKFZ, The University Hospital Heidelberg (UKHD), The Heidelberg Medical Faculty of the Heidelberg University, and The Thorax Clinic Heidelberg, Heidelberg, Germany; Division of Applied Functional Genomics, German Cancer Research Center (DKFZ), Heidelberg, Germany; National Center for Tumor Diseases (NCT), NCT Heidelberg, a partnership between DKFZ and Heidelberg University Hospital, Heidelberg, Germany; German Cancer Consortium (DKTK), Core Center Heidelberg, Heidelberg, Germany; Translational Neuroblastoma Research, German Cancer Research Center (DKFZ), Heidelberg, Germany; Hopp Children’s Cancer Center (KiTZ), Heidelberg, Germany; Division of Translational Medical Oncology, German Cancer Research Center (DKFZ), Heidelberg, Germany; Department for Translational Medical Oncology, National Center for Tumor Diseases (NCT/UCC) Dresden, a partnership between DKFZ, Faculty of Medicine and University Hospital Carl Gustav Carus, TUD Dresden University of Technology, and Helmholtz-Zentrum Dresden-Rossendorf (HZDR), Germany; Translational Medical Oncology, Faculty of Medicine and University Hospital Carl Gustav Carus, TUD Dresden University of Technology, Dresden, Germany; German Cancer Consortium (DKTK), partner site Dresden; Computational Oncology Group, Molecular Precision Oncology Program, NCT Heidelberg and DKFZ, Heidelberg, Germany; Innovation and Service Unit for Bioinformatics and Precision Medicine, DKFZ, Heidelberg, Germany; Pattern Recognition and Digital Medicine Group, Heidelberg Institute for Stem Cell Technology and Experimental Medicine, Heidelberg, Germany; Sarcoma Unit, Mannheim Cancer Center (MCC), University of Heidelberg / Mannheim University Medical Center, Mannheim, Germany; Division of Surgical Oncology, Medical Faculty Mannheim, University of Heidelberg, 68167 Mannheim, Germany; Deparment of Molecular Medicine, University of Tübingen; Department of Hematology, Oncology and Cancer Immunology, Charité-Universitätsmedizin Berlin, Corporate Member of Freie Universität Berlin, Humboldt-Universität zu Berlin, and Berlin Institute of Health, Berlin, Germany; Comprehensive Cancer Center, Charité-Universitätsmedizin Berlin, Corporate Member of Freie Universität Berlin, Humboldt-Universität zu Berlin, and Berlin Institute of Health, Berlin, Germany; German Cancer Consortium (DKTK) partner site Berlin and German Cancer Research Center (DKFZ), Heidelberg, Germany; National Center for Tumor Diseases (NCT), NCT Berlin, a partnership between DKFZ, Charité Universitätsmedizin, BIH and MDC, Berlin, Germany; Center for Personalized Medicine, TUM School of Medicine and Health, TUM University Hospital, Munich, Germany; Clinical Department of Internal Medicine III, TUM School of Medicine and Health, TUM University Hospital, Munich, Germany; Bavarian Cancer Research Center (BZKF), Munich, Germany; Institute of Pathology, Technical University Munich, TUM School of Medicine and Health, Munich, Germany; University Cancer Center, University Medical Center, Johannes Gutenberg University Mainz, Langenbeckstraße 1, 55131 Mainz, Germany; 3rd Medical Department, University Medical Center, Johannes Gutenberg University Mainz, Langenbeckstraße 1, 55131 Mainz, Germany; TRON-Translational Oncology, University Medical Center, Johannes Gutenberg University Mainz, Freiligrathstraße 12, 55131 Mainz, Germany; German Cancer Consortium (DKTK), partner site Frankfurt/Mainz, a partnership between DKFZ and University Medical Center Mainz, Germany; Department of Medicine, Hematology/Oncology, University Hospital, Goethe University, Frankfurt, Germany; University Cancer Center Frankfurt (UCT), University Hospital, Goethe University, Frankfurt, Germany; German Cancer Consortium, Partner Site Frankfurt/Mainz and German Cancer Research Center, Heidelberg, Germany; Institute of Medical Bioinformatics and Systems Medicine, Medical Center - University of Freiburg, Faculty of Medicine, University of Freiburg, Freiburg, Germany; German Cancer Consortium (DKTK), Partner site Freiburg, a partnership between DKFZ and Medical Center - University of Freiburg; Institute of Pathology, LMU Munich; Dept. of Internal Medicine and Sarcoma Center, University Hospital Essen, University Duisburg-Essen; DKTK, Partner Site Essen, Essen, Germany; Institute of Human Genetics, Heidelberg University, Heidelberg, Germany; Department of Pediatric Hematology and Oncology, University Hospital, Im Neuenheimer Feld 430, 69120, Heidelberg, Germany; European Molecular Biology Laboratory, European Bioinformatics Institute, Hinxton CB10 1SA, UK; Somatic Genomics Programme, Wellcome Sanger Institute, Hinxton, UK; European Molecular Biology Laboratory (EMBL), Genome Biology Unit, Heidelberg, Germany; Bridging Research Division on Mechanisms of Genomic Variation and Data Science, German Cancer Research Center (DKFZ), Heidelberg, Germany

**Author notes:** Corresponding author: Priya Chudasama, Ph.D.

## Abstract

Complex karyotype sarcomas (CKS) are heterogeneous mesenchymal malignancies that typically lack recurrent actionable oncogenic drivers and remain therapeutically challenging. Loss of ATRX is a recurrent feature of CKS and defines a particularly high-risk subgroup. ATRX loss is also associated with activation of the alternative lengthening of telomeres (ALT) pathway, and ALT-positive sarcomas have been linked to poor clinical outcomes. However, the molecular underpinnings underlying ALT-status-dependent differences in CKS, as well as the therapeutic vulnerabilities associated with ALT, remain poorly defined. By integrating C-circle-based ALT detection across 776 sarcoma samples with multi-modal sequencing of five CKS subtypes, we find that ALT activity is associated with enriched hallmarks of genomic instability. ALT-positive transcriptomes are dominated by a coordinated DNA damage response and mitotic program, in contrast to oncogenic signaling pathways that drive TERT activation in ALT-negative tumors. Long-read sequencing reveals telomere repeat clusters and telomere-mediated healing at structural breakpoints in ALT-positive tumors. These events also occur on extrachromosomal DNA (ecDNA), linking ALT activity to ecDNA biology. Together, our findings position ALT status as an important stratifying feature of CKS and identify ALT-associated transcriptional programs as potential therapeutic targets.

## Introduction

Sarcomas comprise more than 150 mesenchymal malignancies that, despite their rarity, account for a large share of cancer-related morbidity in young adults and remain among the most challenging cancers to treat across all age groups^1,2^. From a genomic perspective, sarcomas separate into two broad groups^3^. The first group is defined by simple, near-diploid karyotypes and a single recurrent driver, typically a pathognomonic gene fusion^3^ (e.g., Ewing sarcoma, synovial sarcoma, alveolar soft part sarcoma) or a single oncogenic mutation^4^ (*KIT* in GIST), which serves both as a diagnostic marker and an actionable therapeutic target. The second comprises complex karyotype sarcomas (CKS), that lack a recurrent oncogenic drivers and are characterized by complex, highly rearranged genomes with widespread copy-number alterations, frequent whole-genome doubling, bi-allelic *TP53* and *RB1* loss, and chromothripsis^5,6^. As a result, CKS present high-levels of inter- and intratumor heterogeneity, which drives tumor resistance.

Loss of the chromatin remodeling factor *ATRX* is a frequent event in CKS and defines a clinically high-risk subset of these tumors^7,8^. *ATRX* loss has been mechanistically linked to the alternative lengthening of telomeres (ALT) pathway, a recombination-based, telomerase-independent route to replicative immortality. ALT activity can be measured experimentally with the C-circle assay, which detects partially single-stranded telomeric DNA circles specific to ALT-positive (ALT+) cells^9,10^, or in situ assays such as ALT-FISH or detection of ALT-associated promyelocytic leukemia (PML) body^11,12^. In parallel, computational approaches applied to whole-genome sequencing data have been developed to infer ALT status at scale, including *ATRX/DAXX* alteration status, classifiers built on telomere content and variant repeat features in the PCAWG pan-cancer cohort^13^ and detection of outward telomere fusions that arise specifically from ALT activity^14^. Direct comparisons have shown that genomic classifiers and assay-based readouts do not always agree^15^, reinforcing the value of the C-circle assay as the standard for ALT calling. Using these tools, ALT has emerged as a clinically consequential feature in several cancers. ALT positivity is consistently associated with poorer overall survival, higher metastatic potential, and worse therapeutic response^16,17^. In high-risk neuroblastoma it defines a distinct subgroup with poor outcome and stratifies patients independently of MYCN amplification and TERT rearrangements^18,19^. In sarcomas, however, systematic ALT profiling has been limited to a few subtypes, most notably leiomyosarcoma^5^ and osteosarcoma^6^, leaving the prevalence and biology of ALT across the broader CKS landscape largely uncharacterized.

By profiling ALT frequency across a large and diverse pan-sarcoma cohort and integrating multi-omics data from four chromosomally unstable subtypes, we show that the telomere maintenance mechanism used by CKS shapes their genomic architecture and complexity, as well as the transcriptional programs. Using long-read sequencing, we identify ALT-associated telomere healing as a recurrent mechanism in CKS that links telomere biology to the genomic instability of these tumors. Together, our findings establish TMM status as a biologically and clinically meaningful feature of CKS biology, and provide rationale for therapeutics development in this aggressive disease.

### ALT is a frequent feature of complex karyotype sarcoma

To systematically investigate frequency of ALT across human sarcomas, we performed C-circle assay^10^ in a cohort of 776 primary sarcoma samples representing 57 distinct sarcoma subtypes, profiled within the DKFZ/NCT/DKTK-MASTER registry trial^20^. A comprehensive list of sarcoma subtypes analyzed in this study is provided in **Supplementary Table 1**. The cohort comprised 54% male and 46% female patients, with most showing >50% purity (**Figure 1a, upper panel, Supplementary Figure 1**). We detected C-circles in 22 out of 57 subtypes, with higher ALT frequency (20-66%) in CKS, including uterine leiomyosarcoma (ULMS, 66%), soft-tissue LMS (referred to as LMS in this study, 64%), pleomorphic liposarcoma (PLS, 50%), undifferentiated soft tissue sarcoma (USTS, 42%), myxofibrosarcoma (MFS, 33%), osteosarcoma (OS, 33%), and dedifferentiated liposarcoma (DDLS, 24%) (**Figure 1a, lower panel**). Simple karyotype sarcomas showed low frequency (synovial sarcoma, 3%) to absence of ALT (Ewing sarcoma, alveolar soft part sarcoma). Given the high ALT frequency and sample size in our cohort, we focused our comparative analysis of ALT+ and ALT-negative (ALT–) tumors on CKS subtypes DDLS, LMS, ULMS, OS, and USTS, integrating whole-exome sequencing (WES, n=179), whole-genome sequencing (WGS, n=146) and RNA sequencing (RNASeq, n=260) data of tumor specimens.

**Figure 1.**
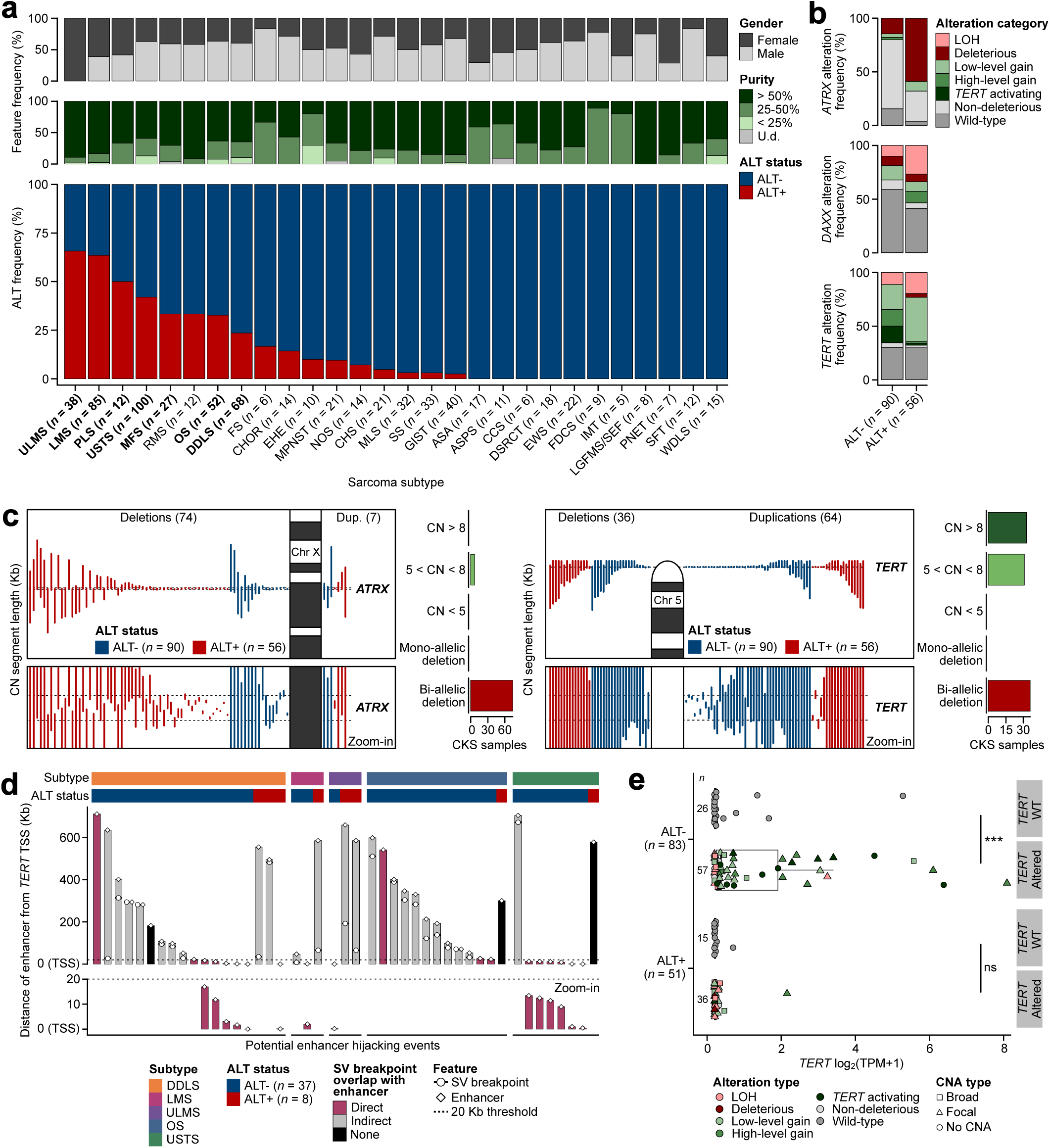
ALT is frequent in human complex karyotype sarcomas. (**a**) Stratification of human bone and soft-tissue sarcoma samples (*n* = 778) based on gender, purity, and ALT frequency determined using the C-circle assay. Cases with C-circle levels ≥ 0.1 arbitrary units (AU) were considered ALT+. CKS subtypes with an ALT frequency ≥ 20% are highlighted in bold. Subtypes with less than five samples were excluded. A comprehensive list of sarcoma subtype abbreviations is provided in **Supplementary Table 1**. (**b**) Frequency of somatic SNVs and indels, broad and focal CNAs, and SVs in *ATRX*, *DAXX*, and *TERT* across the CKS cohort (*n* = 146, WGS only), including DDLS, LMS, ULMS, OS, and USTS, grouped as ALT- and ALT+. Somatic alterations in these genes were integrated with RNA-Seq data relative to gene-specific wild-type log2(TPM+1) thresholds (reduced expression: ≤5^th^ percentile, increased expression: ≥75^th^ percentile). The following categories were defined: LOH (locus-level loss of heterozygosity), deleterious (loss-of-function variants, variants with CADD PHRED ≥ 30, missense variants with REVEL ≥ 0.5, homozygous deletions, and/or other loss events supported by reduced expression), low-level gain (duplications without increased expression), high-level gain (duplication/amplification events or duplications with increased expression), *TERT* activating (promoter mutations and/or enhancer hijacking events), non-deleterious, or wild-type. (**c**) GenomeTornadoPlot of the *ATRX* and *TERT* loci displaying focal copy number (CN) segments ranked by size and alteration type (deletions vs. duplications) and grouped by ALT status (left panel). The total number of affected CKS samples stratified by CN status is shown in the right panel. (**d**) Distribution of SVs involving dbSUPER enhancer regions within 0.7 Mb upstream of the *TERT* transcription start site (TSS). SV breakpoint overlaps are color-coded. Tumors with potential enhancer hijacking events were identified by applying a threshold of 20 Kb from the *TERT* TSS for the location of both the SV breakpoint and the rearranged enhancer site (represented by distinct symbols). CKS samples were arranged by subtype and ALT status. (**e**) Comparison of *TERT* expression levels between ALT- and ALT+ CKS samples with WGS and RNA-Seq data (*n* = 134), stratified by *TERT* alteration status (two-sided Wilcoxon rank-sum tests with Benjamini-Hochberg correction; **** *p* < 0.0001, ns: not significant). The horizontal line within each boxplot represents the median TPM value. The sample size (*n*) for each group is indicated. CKS: complex karyotype sarcoma, U.d.: undetermined, CNA: copy number alteration, SV: structural variant, TPM: transcripts per million, WT: wild-type, DDLS: dedifferentiated liposarcoma, LMS: leiomyosarcoma, ULMS: uterine leiomyosarcoma, OS: osteosarcoma, USTS: undifferentiated soft-tissue sarcoma.

### Genomic alterations at *ATRX*, *DAXX*, and *TERT* are unreliable surrogate markers for ALT

The process of ALT activation has been linked to the loss of the ATRX/DAXX chromatin remodeling complex that disrupts telomeric heterochromatin^13,21,22^, and alterations in *ATRX/DAXX* have frequently been used as surrogate markers for ALT positivity^13,23,24^. However, *ATRX*/*DAXX* loss may be predictive of ALT activity in specific tumor types and particular genomic or epigenetic contexts^15,25–27^.

To clarify the relationship in CKS, we analyzed *ATRX*, *DAXX*, and *TERT* in each sample with both WGS and RNA-seq data available, classifying each by genomic event, namely mutations, somatic copy number alterations (CNA) and other structural variants (SVs), and integrating with expression consequence (**Figure1b, Supplementary Figure 2a–b, Supplementary Table 1**). Deleterious ATRX aberrations involving truncating mutations, homozygous deletions, or SVs were present in only 59% of ALT+ CKS. The remaining ALT+ cases carried non-deleterious *ATRX* events (29%), low-level copy number gains (9%), or wild-type *ATRX* (4%). Deleterious *ATRX* alterations were also detected in 14% of ALT-negative tumors. Deleterious *DAXX* events were rare and comparable between groups (7–9%), but ALT+ tumors showed enriched *DAXX* loss of heterozygosity (27%) and high-level gains (11%), yielding a higher overall *DAXX* alteration frequency than in ALT– cases (54% vs. 32%). Deleterious *ATRX* or *DAXX* events were associated with reduced expression, as expected (p = 8.09 × 10⁻⁵ and p = 2.57 × 10⁻⁶, Wilcoxon rank-sum test; **Supplementary Figure 2c**). At the *TERT* locus, copy number gains or activating events were detected in 54% of ALT– tumors. LOH was more common in ALT+ (20%) than ALT– (11%) cases. Low-level *TERT* gains in ALT+ tumors (41%) were not associated with increased expression or evidence of *TERT* activation (**Supplementary Figure 2d**).

Collectively, these findings demonstrate that ALT activation in CKS can occur independently of the loss of the *ATRX/DAXX* chromatin remodeling complex, not all observed genomic alterations in *ATRX* are consequential, CKS may exhibit non-canonical alterations driving ALT, and that detection of genomic alterations alone in *ATRX* and *DAXX* is not a reliable marker for ALT activity.

### Focal CNAs and enhancer hijacking represent recurrent mechanisms of ATRX loss and TERT activation in CKS

We next sought to determine which specific mutational mechanisms targeting key TMM effector genes are operative in CKS. To identify which alteration types at TMM effector loci have consequences on expression, we examined CNAs, point mutations, and structural variants alongside RNA-seq data using *GenomeTornadoPlot*^28^. ALT+ tumors carried deep *ATRX* deletions at high focality (focality score 81.74; **Figure 1c, left, Supplementary Figure 2d**), and a smaller number of broader deletions spanning *ATRX*. *ATRX* mutations ALT+ tumors were nonsense, frameshift, and splice-site variants, predicted to be deleterious, whereas, ALT– tumors were missense and other non-synonymous variants (**Supplementary Figure 2d**). *ATRX* SVs comprised single-breakend events, inversions, and deletions, with inactivating SVs restricted to ALT+ tumors.

ALT– tumors carried focal *TERT* duplications and amplifications (copy number 5 to >8; focality score 3837.22; **Figure 1c, right**) and structural rearrangements that placed *TERT* next to super-enhancers from the *dbSUPER* repository^29^ (**Figure 1d**). SVs at the *TERT* locus were present in 31% of CKS (45 of 146), most of them in ALT– tumors. Of these, 14 SVs fell within 20 kb upstream of the TERT TSS with direct enhancer-breakpoint overlap, indicating enhancer hijacking as an activation mechanism^30^; the remainder mapped 20-700 kb from the TSS to predicted super-enhancer regions. *TERT* promoter SVs were more frequent in DDLS than in LMS or ULMS, and 6 of 8 USTS SV events directly overlapped enhancer regions (**Supplementary Figure 2f**). *TERT* coding mutations were present in both ALT+ and ALT– tumors. *TERT* promoter point mutations were rare and detected in only one ALT– USTS case (USTS83; **Supplementary Figure 2d–e**).

*TERT* expression was markedly higher in ALT– tumors with focal duplications or activating alterations than in ALT– tumors wild-type *TERT* (p = 5.58 × 10⁻⁴, Wilcoxon rank-sum test; **Figure 1e**). *TERT* expression above 0.5 log₂(TPM+1) was associated with focal events. Some tumors without *TERT* duplications also showed elevated *TERT* expression, pointing to alternative regulatory mechanisms such as methylation-mediated control^31^. Several tumors carrying activating *TERT* alterations showed no detectable *TERT* expression, likely reflecting known limitations of bulk RNA-seq for *TERT* quantification^32,33^. One ALT+ tumor (DDLS66) carried a *TERT* enhancer hijacking event and showed detectable *TERT* expression, indicating that ALT and telomerase activity can co-occur within the same tumor.

The alterations most consequential for expression in both subgroups were deleterious focal CNAs at *ATRX* and amplifying focal CNAs at *TERT*. Both event types were more frequent in CKS with whole-genome doubling (**Supplementary Figure 2d**).

### ALT activation reshapes the genomic landscape of CKS through polyploidy, chromothripsis, and HRD-like scarring

To assess potential differences in the global genomic landscape, we compared ploidy and somatic alteration burden between ALT− and ALT+ tumors across matched WES (n = 176) and WGS (n = 145) datasets. ALT+ tumors consistently showed higher ploidy (WES median: 3.2 vs. 2.3, p = 0.017; WGS median: 2.92 vs. 2.18; Wilcoxon rank-sum test; **Figure 2a, Supplementary Figure 3a**), higher broad and focal CNA burden (WES: p = 9.94 × 10⁻⁵ and p = 4.15 × 10⁻⁸; WGS: p = 0.021 and p = 2.25 × 10⁻⁵; **Figures 2b-c, Supplementary Figures 3b-c**), elevated SV counts (p = 0.001; **Figure 2d**), and higher somatic mutation burden (WES: p = 0.012; WGS: p = 0.031; all Wilcoxon rank-sum test, **Figure 2e, Supplementary Figure 3d**). Next, we asked if specific SV classes associated with these differences. Indeed, ALT+ OS showed enrichment of duplications, insertions, and H2H and T2T inversions (p < 0.05), while in USTS the increased burden was driven by duplications, H2H and T2T inversions, and translocations (p < 0.001; **Supplementary Figure 3e**). No major TMM-associated differences were observed in DDLS, LMS, or ULMS. Together, these data indicate that ALT activation in CKS is associated with a broadly remodeled genome marked by polyploidy, elevated CNA and SV burden, and subtype-specific structural rearrangements.

**Figure 2.**
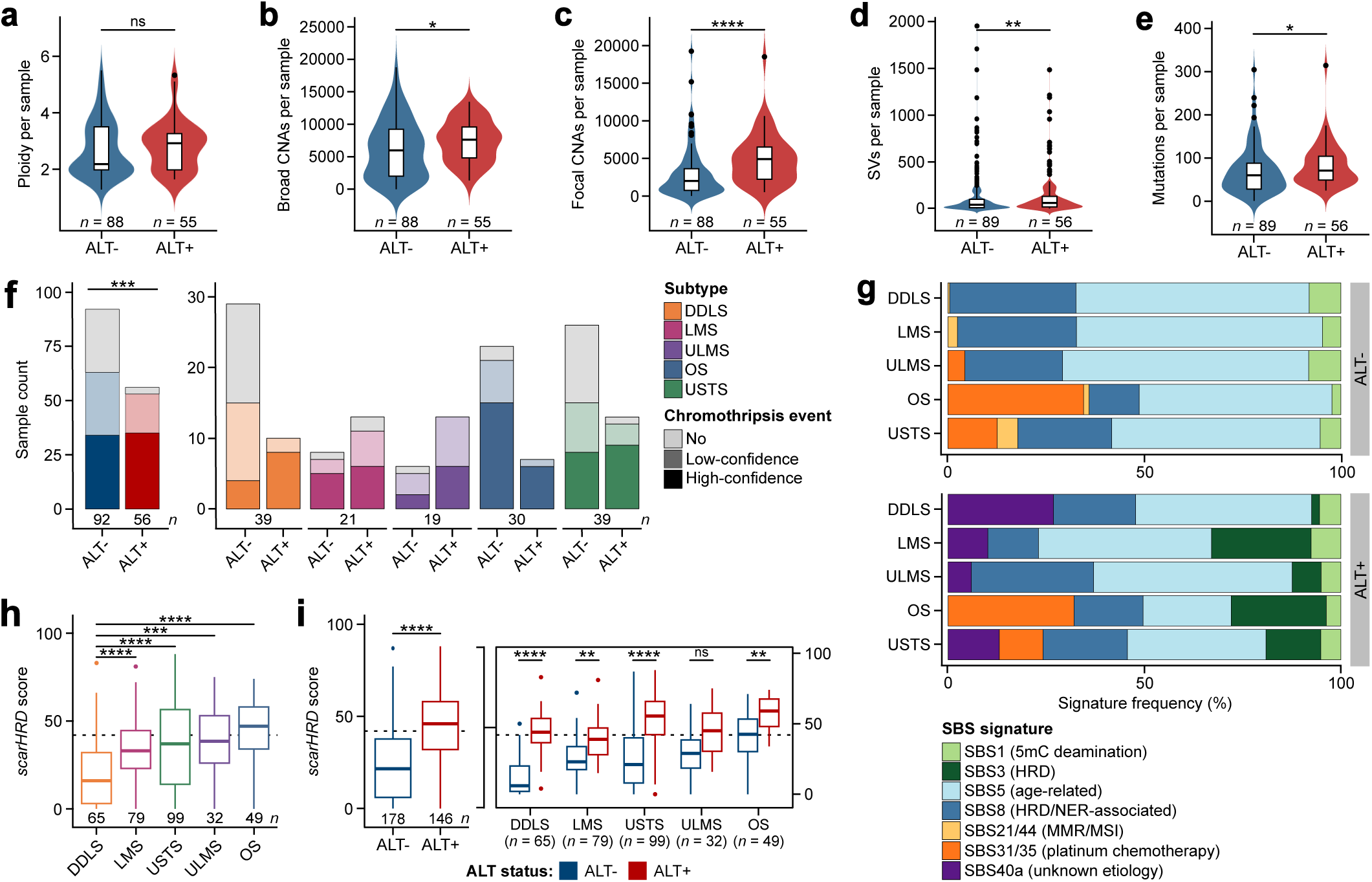
ALT activation is associated with polyploidy, chromothripsis and HRD-like genomic scarring in complex karyotype sarcomas. (**a**) Analysis of ploidy levels between ALT- and ALT+ CKS samples using WGS data (*n* = 145). Samples with extreme ploidy values (>7) were excluded. (**b-e**) Load of somatic broad CNAs (b), focal CNAs (c), SVs (d), and mutations (e) in ALT- and ALT+ CKS tumors based on WGS data. CNA and mutation burden were restricted to protein-coding genes, whereas SV burden was assessed genome-wide to capture both coding and non-coding effects. The width of the violin plots reflects the density of the data distribution. The center horizontal line of boxplots indicates the median value (two-sided Wilcoxon rank-sum tests with Benjamini-Hochberg correction). (**f**) Absolute counts of ALT- and ALT+ CKS samples with or without chromothripsis detected using *ShatterSeek* on WGS data (*n* = 148), shown for the merged cohort (left) and after stratification by CKS subtype (right). Low- and high-confidence calls were defined by 3-6 and ≥7 adjacent CN segments oscillating between two states, respectively. For statistical testing, low- and high-confidence calls were grouped as chromothripsis+ and compared with chromothripsis- samples by ALT status using two-sided Fisher’s exact tests with Benjamini-Hochberg correction. (**g**) Frequency of individual COSMIC SBS signature activities in the CKS cohort grouped by ALT status and individual subtypes (WGS only). Outlier samples with high alteration counts or with signatures linked to sequencing artifacts were excluded from this analysis. (**h**) Quantification of *scarHRD* scores in the CKS cohort (*n* = 324, combined WES and WGS datasets). CKS subtypes are ranked based on increasing median *scarHRD* scores. (**i**) Comparison of *scarHRD* scores between ALT- (*n* = 178) and ALT+ (*n* = 146) CKS samples (left) and further stratification by entity (right). The dotted line represents the *scarHRD* cut-off value set at 42. The center horizontal line of boxplots indicates the median value. Pairwise comparisons were conducted using two-sided Wilcoxon rank-sum tests with the Bonferroni correction. The sample size (*n*) for each group is indicated. CNA: copy number alteration, SV: structural variant, SBS: single-base substitution, HRD: homologous recombination deficiency, NER: nucleotide excision repair, MMR: mismatch repair, MSI: microsatellite instability, DDLS: dedifferentiated liposarcoma, LMS: leiomyosarcoma, ULMS: uterine leiomyosarcoma, OS: osteosarcoma, USTS: undifferentiated soft-tissue sarcoma, ns: not significant, * *p* < 0.05, ** *p* < 0.01, *** *p* < 0.001, **** *p* < 0.0001.

CNAs and SVs in cancer can arise through replication stress, mitotic errors, and breakage–fusion–bridge (BFB) cycles. These processes that also underlie chromothripsis, a catastrophic genomic event in which localized chromosomal shattering and reassembly drives oncogenic rearrangements and poor prognosis^34–36^. Given the elevated CNA and SV burden in ALT+ CKS, we asked whether chromothripsis prevalence differed between ALT− and ALT+ tumors across CKS subtypes. To this end, we analyzed WGS data using *Shatterseek*^35^. Hallmark of chromothripsis events are oscillating CN states, as well as inter or intrachromosomal SVs, indicative of random stitching of breaks via non-homologous end joining (NHEJ) repair^35,37^. As per *Shatterseek* recommendations, we classified chromothripsis calls with 7 contiguous segments oscillating between 2 CN states and at least 3 interleaved intrachromosomal SVs and 4 or more interchromosomal SVs as “high confidence”. The calls with 4, 5 or 6 adjacent segments oscillating between 2 CN states and at least 6 interleaved intrachromosomal SVs were termed as “low confidence”. Strikingly, chromothripsis was detected in 53 of 56 ALT+ CKS samples (95%) compared with 63 of 92 ALT- samples (69%), indicating a significant enrichment of chromothripsis+ cases in the ALT+ subset (p = 1.42 × 10⁻⁴, odds ratio = 8.04, Fisher’s exact test; **Figure 2f, left panel**). At the subtype level, after combining low- and high-confidence calls, chromothripsis was frequent across all CKS entities, affecting 64% of DDLS, 86% of LMS, 95% of ULMS, 93% of OS and 69% of USTS cases, consistent with previous reports^6,38^ (**Figure 2g, right panel**).

In line with its localized nature^35^, chromothripsis showed an overall non-uniform chromosomal distribution, with chromosomes 1, 11, and X most frequently affected (42, 36, and 29 events), and chromosomes 21 and 22 rarely involved (**Supplementary figure 3f**). Subtype-specific patterns included enrichment on chromosome 8 in USTS, chromosomes 9 and 16 in LMS and ULMS, and chromosome 12 in DDLS, while frequent involvement of chromosomes 1 (mainly ALT−), 3, 4, and 11 was observed in OS. TMM-specific patterns were stronger in ALT+ tumors, which related to chromosome-specific enrichment (frequency >50%) with chromothripsis frequencies exceeding 50% across multiple loci (e.g., chromosomes 7, 9, 10, 12, 16–19). Examining the chromothriptic architecture of representative events across subtypes and TMM status using *ReConPlots*^39^(**Supplementary Figure 4a**) revealed near-identical chromothriptic profiles localized to a 50–100 Mb window of chromosome 12 with high CN amplitudes (≥8) and densely interleaved deletions, duplications, and inversions, consistent with the canonical 12q13-15 amplicon in DDLS. For the OS, LMS and USTS, ALT+ chromothriptic events showed broader spatial spread of clustered breakpoints, and greater SV class diversity than their ALT− counterparts.

To identify mutational processes underlying the enriched hallmarks of genomic instability in ALT+ CKS and TMM and subtype-specific patterns of structural rearrangements, we decomposed mutation catalogues from CKS WGS samples (n = 142) using *SigProfilerExtractor* against COSMIC reference signatures^40^ (**Figure 2g**). SBS5 (clock-like, age-related) and SBS8^41^ (HRD- and NER-associated, dominated by C>A and T>A mutations) predominated across all subtypes regardless of ALT status, reflecting the shared age-related and replication-stress mutagenic background of CKS. Platinum chemotherapy-associated SBS31 and SBS35 were detected in OS (32-35%) and USTS (11-13%) cases independently of TMM, consistent with prior treatment exposure.

Importantly, SBS3, the canonical signature of HRD frequently coinciding with biallelic *BRCA1/2* loss^41^, was selectively enriched in ALT+ LMS (25%), OS (24%), and USTS (14%). In addition, SBS40a, a signature of unknown etiology resembling SBS31/35, additionally distinguished a subset of ALT+ tumors. To further interrogate this finding, we computed *scarHRD* scores^42^ and *HRProfiler* probabilities^43^ on merged WES and WGS data (n = 324). Following the threshold of scarHRD ≥ 42 that detects *BRCA1/2*-deficient tumors with 95% sensitivity^44^, and the analogous *HRProfiler* probability ≥ 0.42^43^, we classified samples above HR-deficient. ALT+ tumors displayed significantly higher *scarHRD* scores than ALT− cases (p = 2.40 × 10⁻¹⁸, Wilcoxon rank-sum test; **Figure 2h-i**). OS showed the highest baseline *scarHRD* scores across subtypes (median 47) and DDLS the lowest (median 16), with ALT+ samples shifted upward within each subtype (**Figure 2i**). Stratification by ploidy revealed that *scarHRD* scores rose with ploidy in both ALT subsets, with ALT+ tumors scoring significantly above ALT− counterparts in the <2n and 2n–4n bins, but comparable at >4n (**Supplementary Figure 5b**). *HRProfiler* analysis recapitulated these findings (**Supplementary Figure 5c-e**). Of note, alterations in *BRCA1* & *BRCA2* were rare in our cohort (n=14 in ALT+ CKS, n=8 in ALT− CKS, **Supplementary Table 1**). Combined with the SBS3 enrichment observed in the same tumors, these data indicate that ALT activation in CKS leaves genomic scars normally diagnostic of HR pathway loss, which is paradoxical finding as ALT itself depends on functional HR for RAD51/RAD52-mediated, recombination-driven telomere extension.

Together, these results demonstrate that the choice of telomere maintenance mechanism in CKS is associated with substantial alterations in genome organization, and ALT+ tumors display increased hallmarks of genomic instability compared to ALT− tumors.

### ALT+ and ALT– CKS display distinct chromosome-level CNA landscapes

As CNA burden emerged as the genomic feature with the strongest discrimination between ALT+ and ALT− tumors (**Figures 2b–c; Supplementary Figures 3b–c**), we next interrogated their genome-wide CNA landscapes at chromosome level to identify potential TMM-specific differences. Comparison of CN profiles revealed that ALT− tumors displayed genome-wide low magnitude gains and shallow losses, whereas ALT+ tumors showed comparably frequent and high-magnitude gains and losses across nearly all autosomes (**Figure 3a**).

**Figure 3.**
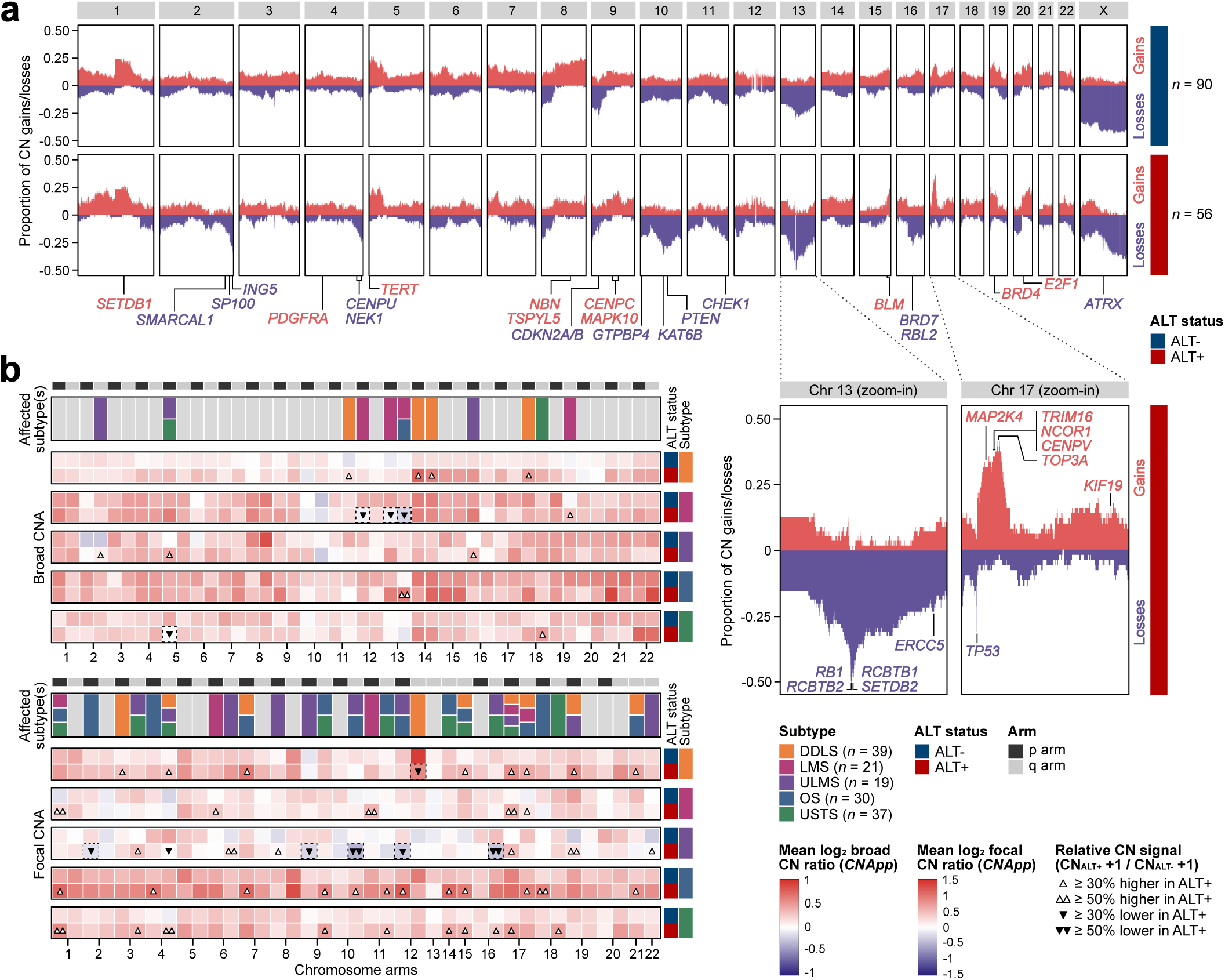
The landscape of broad and focal copy number changes in ALT- and ALT+ complex karyotype sarcomas. (**a**) Proportion of CN gains (red) and losses (purple) across chromosomes for ALT- (upper panel) and ALT+ (lower panel) CKS samples (*n* = 146, WGS only). Zoomed-in sections of chromosomes 13 and 17 were included to emphasize ALT-specific gene-level CN changes. Frequently altered genes associated with tumorigenesis, DNA repair, cell cycle, chromatin remodelling, and telomere maintenance are highlighted along the chromosomes and colored based on CN status in ALT+ cases. (**b**) Arm-level analysis of broad (upper panel) and focal (lower panel) CN changes grouped by CKS subtype and ALT status (WGS only). Heatmaps show the mean *CNApp*-derived log2 CN ratio for each chromosomal arm, calculated separately for ALT-and ALT+ samples within each entity. Single and double arrows denote ≥30% and ≥50% relative differences, respectively, in pseudocount-adjusted mean CN ratios between ALT-and ALT+ CKS samples calculated as (CNALT++1)/(CNALT-+1) within each subtype. Upward and downward arrows indicate higher and lower CN signal in ALT+ samples, respectively. The upper annotation bar indicates the subtype(s) most affected by CNAs for each chromosomal arm, based on relative differences in mean CN ratio. The sample size (*n*) for each group is indicated. CN: copy number, CNA: copy number alteration, DDLS: dedifferentiated liposarcoma, LMS: leiomyosarcoma, ULMS: uterine leiomyosarcoma, OS: osteosarcoma, USTS: undifferentiated soft-tissue sarcoma.

ALT+ tumors carried frequent arm-level losses on 2q, 4q, and 16q. These regions contain *SMARCAL1*, which resolves replication stress at ALT telomeres and whose loss promotes ALT and tumorigenesis^45,46^; *SP100*, an ALT suppressor that sequesters the MRN complex away from ALT-associated PML bodies^47^; *ING5*^48^; the cell-cycle regulators *NEK1* and *CENPU* (4q33–q35.1); and the heterochromatin remodelers *BRD7* and *RBL2* (16q12.1– q12.2; **Figure 3a**). Whole-chromosome losses on 10, 13, and X were also enriched in ALT+ tumors (p = 0.0002, OR = 3.75, Fisher’s exact test; Figure 3a), affecting *PTEN* (10q23.31), *KAT6B* (10q22.2), a 13q14.2 tumor suppressor cluster (*RB1*, *RCBTB1*, *RCBTB2*, *SETDB2*), and *ERCC5* (13q33.1). A focal *ATRX* deletion peak at Xq21.1 was present only in ALT+ tumors (**Figures 1b–c, Supplementary Figure 2d**), whereas segmental X-chromosome losses in ALT– tumors reflected the higher proportion of male patients in that group (58.9% vs. 35.7%).

Further different gains in ALT+ tumors were detected at 1q21.3 (*SETDB1*), 4q12 (*PDGFRA*), 5p15.33 (*TERT*), 9q21.11–q21.13 (*CENPC*, *MAPK10*), and 15q26.1 (*BLM*). An ALT+-enriched high-magnitude amplicon at 17q11.2–q23.3 (*MAP2K4*, *TRIM16*, *NCOR1*, *CENPV*, *TOP3A*, extending to *KIF19*; p = 0.002, OR = 2.98, Fisher’s exact test) which was further resolved by *GenomeTornadoPlot*, that showed multiple focal segments encompassing two or more of these genes (**Supplementary Figure 5b**), identifying 17q11.2–q23.3 as a candidate ALT-specific dependency. In ALT– tumors, the distinguishing feature beyond focal *TERT* gains was broad gain of chromosome 8q, which contains *NBN* (8q21.3) and *RECQL4* (8q24.3) — both involved in telomere replication and resolution of telomeric secondary structures^49,50^.

To resolve subtype- and TMM-associated CN changes, we quantified broad and focal CNAs across CKS subtypes using *CNApp*^51^ and computed pseudocount-adjusted ALT+/ALT– ratios for each subtype (**Figure 3b**). Broad CN ratios were similar between ALT+ and ALT– tumors, with modest differences limited to 5p losses in ALT+ USTS, biarmal chromosome 13 deletions in ALT+ LMS, and 11q, 14q, and 18p gains in DDLS. Focal CNAs separated subtypes more clearly. ALT+ DDLS, OS, and USTS showed higher focal amplifications, including recurrent 1p gains. ALT+ ULMS instead carried focal losses at 2p, 4q, 9p, 10q, 12p, and 16q. The diagnostic 12q13–15 amplicon harboring *MDM2, CDK4*, and *HMGA2* in DDLS^52,53^ showed lower focal gains in ALT+ than ALT– cases. Across all subtypes except OS, ALT+ tumors carried higher focal gains at 17q, matching the amplified gene cluster described above.

These findings identify a qualitatively distinct pattern of copy number alterations in ALT+ CKS, focused on focal rather than arm-level events, and recurrent at loci involved in DNA repair, cell cycle, chromatin remodeling, and telomere maintenance. Subtype-specific focal CNA patterns indicate that ALT status is associated with genome architecture together with lineage-specific biology^54–56^.

### TMM status partitions the CKS transcriptomes into DDR–mitotic and telomerase-associated oncogenic signaling programs

To assess transcriptional consequences of TMM activation, we analyzed RNA-seq data from 260 CKS samples spanning DDLS (n = 57), LMS (n = 65), ULMS (n = 34), OS (n = 48), and USTS (n = 56) (**Supplementary Figure 6a**), classified as ALT− (n = 142) or ALT+ (n = 118) by C-circle assay (**Supplementary Figure 6b**). Partial least squares discriminant analysis revealed clear TMM-driven separation within every subtype (**Figure 4a**), indicating that choice of TMM imposes a consistent transcriptional signature across CKS lineages. Differential expression analysis using *limma-voom*^57^ on 16,743 protein-coding genes identified 1,677 genes significantly altered between ALT subsets (|log₂FC| ≥ 0.6, BH-adjusted p ≤ 0.01), with 888 upregulated and 789 downregulated in ALT+ tumors (**Figure 4b, Supplementary Table 1**).

**Figure 4.**
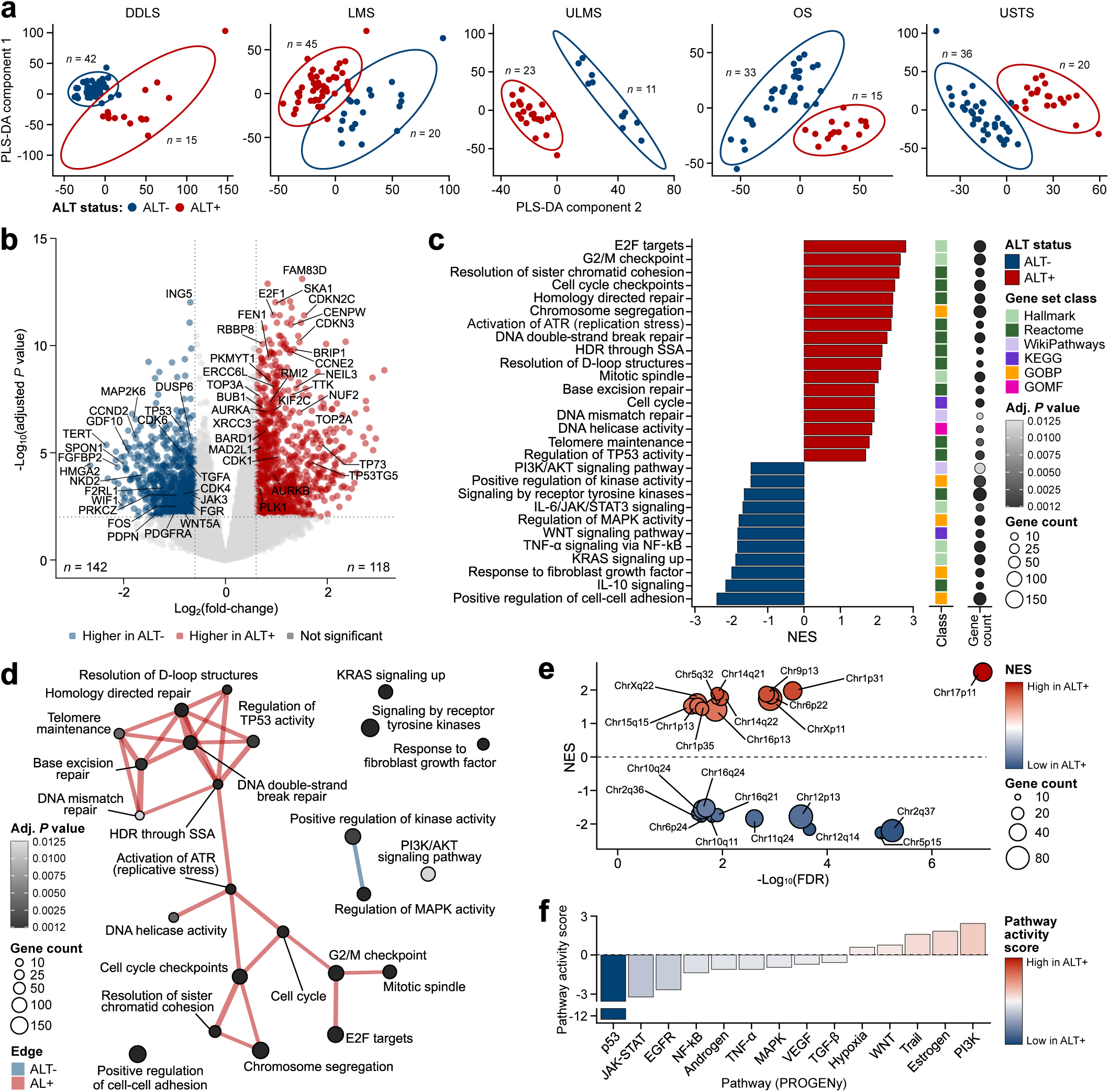
TMM status is associated with distinct transcriptome programs in complex karyotype sarcomas. (**a**) Partial least squares discriminant analysis (PLS-DA) of CKS transcriptomes based on ALT status. The ellipses denote the 95% confidence interval for each group. (**b**) Differential gene expression analysis between ALT- and ALT+ CKS samples (*n* = 260) using the *limma-voom* workflow. Differentially expressed genes (DEG) were defined using thresholds of log₂(fold-change) ≥ 0.6 or ≤ -0.6 and *P* value ≤ 0.01 (Benjamini-Hochberg correction). The top deregulated genes are highlighted. (**c**) Gene set enrichment analysis (GSEA) of CKS transcriptomes stratified by ALT status using the *genekitr* package. Gene sets enriched in ALT- and ALT+ cases are ranked by normalized enrichment score (NES) and classified into six categories: Hallmark, Reactome, WikiPathways, KEGG, GOPB, and GOMF. Dot size and color reflect the total gene count and *P* value (Benjamini-Hochberg correction) in each set, respectively. (**d**) Network of enriched gene sets derived from GSEA. Nodes represent the enriched terms, while edges indicate overlapping gene sets. The thickness of the edges corresponds to the number of overlapping genes. Dot size and color represent the total gene count and *P* value (Benjamini-Hochberg correction) in each set, respectively. (**e**) Cytobands enriched with DEGs, where red and blue indicate higher or lower enrichment in ALT+ cases, respectively. NES values of selected cytobands were plotted against their - log10(FDR) values, with dot size representing the total gene count for each cytoband. (**f**) Activity inference of curated PROGENy pathways in the CKS cohort using *decoupleR*. The pathway activity score is color-coded to show whether it is higher or lower in ALT+ tumors. The sample size (*n*) for each group is indicated. KEGG: Kyoto encyclopedia of genes and genomes, GOBP: gene ontology biological process, GOMF: gene ontology molecular function, SSA: single-strand annealing, HDR: homology-directed repair, FDR: false discovery rate, DDLS: differentiated liposarcoma, LMS: leiomyosarcoma, ULMS: uterine leiomyosarcoma, OS: osteosarcoma, USTS: undifferentiated soft-tissue sarcoma.

Gene set enrichment analysis (**Figure 4c, Supplementary Figure 6c**) identified two transcriptional programs upregulated in ALT+ CKS. The first comprised DNA repair and replication stress responses: homology-directed repair (NES = 2.44, p = 4.16 × 10⁻¹⁰; *RMI2*, *BRIP1*, *RBBP8*, *XRCC3*, *BARD1*), ATR activation under replication stress (NES = 2.39, p = 1.74 × 10⁻⁶; *RAD1*, *CDK2*, *CHEK1*, *RPA1*), telomere maintenance (NES = 1.79, p = 0.004; *H2AZ1*, *POLD3*, *BLM*, *POT1*), base excision repair (NES = 1.93, p = 0.001), and DNA mismatch repair (NES = 1.93, p = 0.01), consistent with the role of ATR–CHEK1 signaling in recombination-based telomere elongation^58^. The second program comprised cell cycle and mitotic regulation: chromosome segregation (NES = 2.42, p = 6.46 × 10⁻¹⁹), G2/M and cell cycle checkpoints (NES = 2.49, p = 6.05 × 10⁻¹⁹; *CDK1*, *PLK1*, *AURKB*, *TOP2A*, *TOP3A*, *BUB1*), and E2F target genes (NES = 2.79, p = 2.99 × 10⁻²⁷; *AURKA*, *E2F8*, *KIF2C*, *MAD2L1*, *PLK4*). These two transcriptional programs strongly align with process of telomere synthesis in ALT: a homology-directed repair step in G2 cell cycle that takes place inside APBs and depends on ATR–CHEK1 signaling along with BLM, POLD3, and RAD52, followed by the actual telomere extension step at the G2-to-mitosis transition, carried out by mitotic DNA synthesis (MiDAS)^55^. Upregulation of both DDR and mitotic programs at the same time is points to high demand and activity of for these genes, highlighting potential for DDR- and mitosis-directed agents in these tumors.

ALT– tumors showed upregulation of oncogenic and inflammatory signaling, including KRAS (NES = −1.88, p = 1.87 × 10⁻⁶; *DUSP6*, *MYCN*, *MAP3K1*), TNF-α/NF-κB (NES = −1.83, p = 8.35 × 10⁻⁶), WNT (NES = −1.82, p = 5 × 10⁻⁴), MAPK (NES = −1.78, p = 1 × 10⁻⁴), IL-6/JAK/STAT3 (NES = −1.68, p = 0.003), and PI3K/AKT (NES = −1.45, p = 0.01; *CDK4*, *CDK6*, *CCND2*, *PDGFRA*). Each of these pathways induces *TERT* expression or activity: WNT/β-catenin and KRAS–MAPK activate *TERT* transcription through c-MYC and AP-1^59,60^, IL-6/STAT3 and TNF-α/NF-κB act at the *TERT* promoter^61^, and PI3K/AKT phosphorylates TERT to promote nuclear localization and enzymatic activity^62^. Semantic similarity-based network analysis of enriched gene sets confirmed this dichotomy (**Figure 4d**): ALT+ programs formed two tightly interconnected modules spanning DDR/TMM (p = 0.0006 to 4.15 × 10⁻¹²) and cell cycle (p = 3.36 × 10⁻⁶ to 5.98 × 10⁻²⁹), whereas ALT− programs were dispersed across heterogeneous oncogenic signaling nodes. Collectively, our data establish oncogenic signaling pathway activation closely linked to different mechanisms of TERT activation as dominant programs of ALT–CKS, distinct from DDR-mitosis programs of ALT+CKS.

Notably, ALT+ transcriptional changes mapped onto the CN landscape (**Figures 3a**, **4e**). Upregulated programs were concentrated at recurrent gains on 17p11–q23.3 (*GID4*, *NCOR1*, *CENPV*, *TOP3A*), 1p13–p35 (*RBBP4*, *DCLRE1B*, *GADD45A*), 14q21–q22 (*CDKL1*, *CDKN3*, *FANCM*), 15q15 (*BUB1B*, *KNL1*), and Xp11 (*CDK16*, *HDAC6*, *KDM6A*). Downregulated programs mapped to recurrent deletions on 2q36–q37 (*HDAC4*, *SP100*, *ING5*), 10q11–q24, and 16q21–q24 (*FANCA*). The ALT+ transcriptome therefore reflects gene dosage at recurrent CNAs affecting telomere maintenance, chromatin remodeling, and DNA repairs, in turn indicating that CNA patterns in ALT+CKS are biologically meaningful.

Pathway activity inference using *decoupleR*^63^ with *PROGENy*^64^ recapitulated and extended these findings (**Figure 4f, Supplementary Figure 6d**). Strikingly, p53 pathway activity was consistently suppressed in ALT+ CKS (activity score = −12.9), consistent with the high frequency of inactivating *TP53* alterations in this group and with the GSEA-derived enrichment of *TP53* regulation (NES = 1.69, p = 0.001). Beyond genetic loss, transcriptional uncoupling of *p53* was evident: positive regulators normally restrained by p53 were elevated, including FANCD2^65^ (replication fork stability), POLQ^66^ (mediates survival in TP53-mutant cancers), and AURKA^67^ (a p53 antagonist), while p53 effectors TP53I3^68^ (ROS-mediated apoptosis), CDKN1A^69^ (cell cycle arrest), and PLK3^70^ (p53 stabilization) were repressed (**Supplementary Figure 7e**). Together, these findings suggest DDR upregulation, enhanced mitotic control while transcriptionally dismantling p53-mediated genome surveillance in CKS lacking genomic inactivation as key characteristics of ALT+ CKS.

### Elevated telomere content and telomere variant repeat enrichment characterize ALT+ CKS

ALT+ tumor cells carry telomere variant repeats (TVRs) in addition to canonical TTAGGG hexamers^71^. TVRs spread from proximal telomeric and subtelomeric regions through recombination^72^ and create binding sites for sequence-specific factors, including the orphan nuclear receptors COUP-TFII and TR4^72,73^ and the zinc finger proteins ZBTB10 and ZNF524^74,75^. Binding of these factors remodels telomeric chromatin and supports recombination-based telomere extension in ALT cells.

To characterize the telomeric landscape of CKS, we estimated telomere content and variant repeat composition using *TelomereHunter*^76^ on WES (n = 169) and WGS (n = 146) data. Analysis of telomere content across CKS (**Figure 5a**) and other sarcoma entities (**Supplementary Figure 7a**) revealed highest median telomere content (expressed as log2 ratio of telomere content between tumor and control samples) in ULMS (0.83), followed by LMS (0.54), OS (0), USTS (-0.08), and DDLS (-0.51). ALT+ CKS samples had higher telomere content than ALT– samples in both WES and WGS cohorts (adjusted p < 0.0001 in both; Wilcoxon rank-sum test; **Figure 5b**).

**Figure 5.**
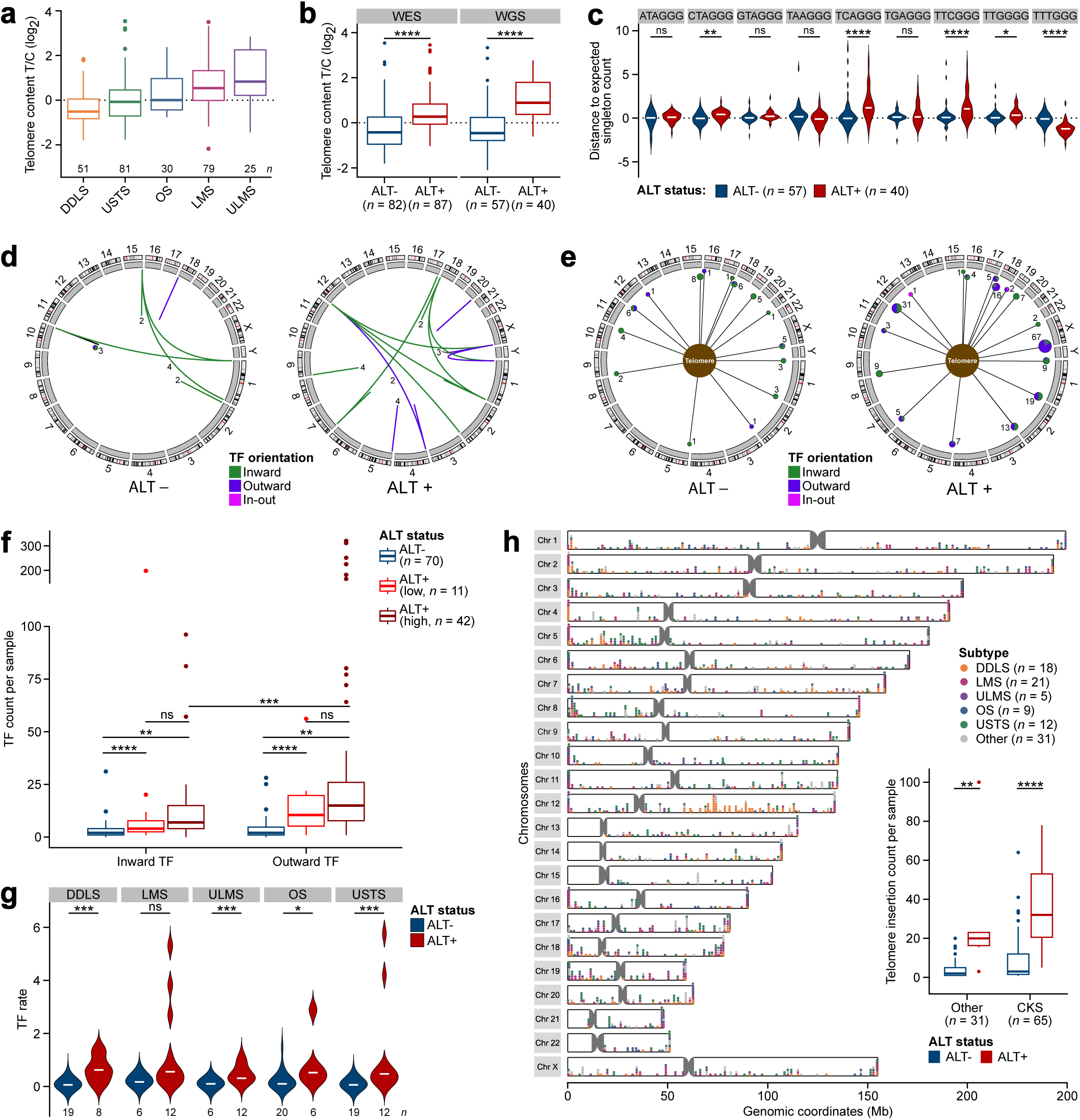
Telomere variant repeats, outward telomere fusions, and telomere insertions are enriched in ALT+ complex karyotype sarcomas and favor recombination processes. (**a**) Telomere content analysis across the CKS cohort (*n* = 266, combined WES and WGS datasets) calculated as log2 ratio of telomere content between tumor and control samples. (**b**) Comparison of telomere content distribution between WES (*n* = 169) and WGS (*n* = 97) datasets from CKS samples grouped based on ALT status. (**c**) Distribution of telomere variant repeat (TVR) content between ALT- and ALT+ CKS tumors (WGS only) displayed as distance to expected singleton counts. (**d-e**) Circos plots showing the genome-wide distribution and frequency of telomere fusion (TF) events with at least one anchored subtelomeric mate (MQ ≥ 10). For events with two anchored subtelomeric mates (d), both breakpoint positions could be assigned to specific chromosome ends. Each arc represents a fusion between two subtelomeric regions. The arc color indicates fusion orientation, while the numbers next to selected arcs denote the corresponding TF count. For events with one anchored subtelomeric mate and one telomeric mate (e), the subtelomeric breakpoint is connected to a central telomeric hub. Pie chart size and adjacent labels indicate event counts, whereas colors denote fusion orientation. (**f**) Comparison of inward and outward TF counts between ALT- and ALT+ CKS tumors. ALT+ cases were further stratified based on C-circle levels (ALTlow: 0.1 - 0.2, ALThigh ≥ 0.2). (**g**) Comparative analysis of TF rate between ALT- and ALT+ samples stratified by CKS entity. The TF rate is defined as the number of TFs normalized to 1X genome coverage and corrected for sequencing depth, read length, and tumor purity. (**h**) Genome-wide chromosomal location of telomere insertions (left panel) and analysis of telomere insertion counts (right panel) between ALT status groups in the pan-sarcoma cohort (*n* = 96, WGS only). CKS cases are color-coded, while other sarcoma entities are shown in grey. The width of the violin plots reflects the density of data distribution. The center horizontal line of boxplots indicates the median value. All pairwise comparisons were conducted using two-sided Wilcoxon rank-sum tests with Benjamini-Hochberg correction (ns: not significant, * *p* < 0.05, ** *p* < 0.01, *** *p* < 0.001, **** *p* < 0.0001). The sample size (*n*) for each group is indicated. T/C: tumor/control, CKS: complex karyotype sarcoma, Mb: megabase, DDLS: dedifferentiated liposarcoma, LMS: leiomyosarcoma, ULMS: uterine leiomyosarcoma, OS: osteosarcoma, USTS: undifferentiated soft-tissue sarcoma, MQ: mapping quality.

We then quantified singletons, i.e., TVRs embedded within canonical telomeric repeats, as markers of ALT activity^13,18^. Consistent with prior pan-cancer observations in tumors with truncating *ATRX*/*DAXX* alterations^13^, ALT+ CKS samples were enriched for CTAGGG (adjusted p = 1.14 × 10⁻³), TCAGGG (adjusted p = 4.54 × 10⁻⁵), TTCGGG (adjusted p = 9.3 × 10⁻⁵), and TTGGGG (adjusted p = 1.27 × 10⁻²) variants, and depleted for TTTGGG repeats (adjusted p = 3.1 × 10⁻⁹; Wilcoxon rank-sum test; **Figure 5c**). C-type repeats (TCAGGG) have reduced TRF2 binding affinity^77^, and their enrichment in ALT+ CKS is consistent with shelterin destabilization, nuclear receptor recruitment, and a DNA damage response that supports HR-mediated telomere elongation^77^.

### Outward telomere fusions are a hallmark of ALT+ CKS

Telomere fusions (TFs) generate dicentric chromosomes and anaphase bridges, driving genomic instability through chromothripsis and BFB cycles^38,78^. Among TFs, outward fusions (CCCTAA→TTAGGG) are a recently described marker of ALT, distinct from inward TFs (TTAGGG→CCCTAA) that arise from classical end-to-end joining^14^. To test whether this ALT signature is present in CKS and to identify subtype-specific patterns, we applied *TelFusDetector*^14^ to WGS data from 148 CKS samples.

ALT+ CKS carried more TFs than ALT– tumors across both orientations, stratified by C-circle level (ALT-low 0.1–0.2 and ALT-high >0.2; inward p = 7.5 × 10⁻³ and p = 1.69 × 10⁻⁷; outward p = 1.03 × 10⁻³ and p = 6.7 × 10⁻¹²; Wilcoxon rank-sum test; **Figure 5f**). ALT-high tumors carried a clear excess of outward over inward TFs (adjusted p = 3.79 × 10⁻⁴), with some samples exceeding 200 outward events. Outward TFs in ALT+ tumors formed large clusters at chromosomes 12, 18, and X (**Figure 5d–e**); in ALT– tumors, TFs were sparse and scattered. Across subtypes, ALT+ CKS had higher TF rates than ALT– (0.662–4.579 vs. 0.138–0.686; **Figure 5g**), with significant differences in DDLS (adjusted p = 6.95 × 10⁻⁴), ULMS (adjusted p = 5.37 × 10⁻⁴), OS (adjusted p = 1.64 × 10⁻²), and USTS (adjusted p = 5.37 × 10⁻⁴). The TF rate difference was less pronounced in LMS (adjusted p = 0.102), despite a higher median in ALT+ LMS (0.554 vs. 0.171). The correlation between telomere content and TF rate was modest in both ALT– (Pearson R = 0.35, p = 0.02) and ALT+ tumors (Pearson R = 0.31, p = 0.06; **Supplementary Figure 7b**), consistent with the coexistence of critically short and aberrantly elongated telomeres in ALT+ tumors that uncouples bulk telomere content from fusion frequency. Together, these findings establish outward TFs as a hallmark of ALT+ CKS.

In addition to telomere and sub-telomeric regions, telomeric repeats can be added de novo to intrachromosomal double-strand breaks by telomerase-mediated chromosome healing^79^ or acquired from another chromosomal position through recombination-mediated telomere capture^80^. To assess the prevalence of potential intra chromosomal telomere insertions across CKS subtypes stratified by ALT status, we applied *TelomereHunter*^76^ to WES and WGS cohorts of CKS and other non-CKS sarcoma. The telomeric insertion burden was significantly higher in ALT+ than ALT− tumors (adjusted p = 1.24 × 10⁻³), an effect even more pronounced in CKS samples alone (adjusted p = 3.9 × 10⁻⁷; Wilcoxon rank-sum test; **Figure 5g**). Interestingly, these insertions were enriched in rearrangement-prone regions including 12q in DDLS and chromosomes 17 and 19 in LMS, suggesting the relevance of healing and capture mechanisms also in ALT+ CKS.

### Telomere repeat clusters in ALT+ CKS occur at structural breakpoints with distinct methylation and motif features at long-read resolution

To characterize the genomic context and structural origin of intrachromosomal telomeric insertions identified by short-read sequencing (**Figure 5h**), we performed Oxford Nanopore long-read sequencing on high-molecular-weight gDNA from eight CKS patient samples. We applied *Lorax*^81^ to determine the landscape of canonical and non-canonical telomeric repeats at SV breakpoints, enabling a single-nucleotide resolution comparison of telomere-associated structural events between ALT− and ALT+ tumors.

We identified frequent telomere insertions, also termed as telomere repeat clusters (TRCs) at SV breakpoints across CKS tumor genomes. In ALT+ tumors, chromosomes carrying TRCs had higher SV counts than chromosomes without TRCs (adjusted p < 0.001, Wilcoxon rank-sum test; **Supplementary Figure 8a**), and TRC count correlated with C-circle levels (Spearman ρ = 0.76, p = 0.03; **Supplementary Figure 8c**). TRC enrichment in ALT+ DDLS mapped to chromosome 12 (**Supplementary Figure 8b**), consistent with the short-read insertion pattern described above (**Figure 5h**). ALT– tumors showed no association between TRCs and SV burden (adjusted p > 0.05; **Supplementary Figure 8a**). TRC length distributions were narrower in ALT– than ALT+ tumors (**Supplementary Figure 8d**). *Lorax* classification identified intra-chromosomal insertions and neo-telomeres in ALT+ tumors across all five subtypes (**Supplementary Figure 8e**). In ALT– tumors, neo-telomeres were detected in a single sample (ULMS28). This pattern is consistent with ATR kinase suppression of telomerase-mediated de novo telomere addition at DSBs^82^, the dominant TMM in ALT− CINS tumors (**Figure 1e**).

Methylation and sequence features at breakpoints may carry information on mechanisms underlying their formation^83,84^. We therefore examined CpG methylation and motif composition within ±1 kb of TRC breakpoints. Both intra-chromosomal insertions and neo-telomeric events were located in regions of high CpG methylation (**Supplementary Figure 8f**). This signal pointed to a heterochromatic environment at TRC sites, consistent with DNA methylation’s established role in stabilizing chromosome ends and limiting aberrant recombination at telomeric and subtelomeric regions^83^. Motif analysis supported this interpretation: NRF1 binding sites were enriched at both intra-chromosomal insertions and neo-telomeres, with NR2F6 motifs additionally enriched upstream of neo-telomeres (**Supplementary Figure 8g**). NR2F6 has been reported to recruit TRIM28, which drives H3K9me3 deposition^85^, and DNA methylation and H3K9me3 are functionally coupled at heterochromatin^86^. Together, the methylation and motif features at TRC sites are consistent with formation in a heterochromatic context, narrowing the candidate mechanisms of TRC genesis to those operating in repressive chromatin.

Next, we characterized the telomeric repeat signal along each consensus sequence to accurately define the position and length of telomeric repeat tracts. **Figure 6a** (middle panels) highlights representative examples of telomeric repeat-containing consensus sequences supporting both single and multiple intra-chromosomal insertions, as well as one-sided and two-sided neo-telomere formation (**Figure 6a, upper panels**). A telomeric repeat signal value of 1 indicates a region composed entirely of canonical telomeric repeats, with the length of this signal corresponding to the size of the repeat region. IGV tracks confirmed these events, typically revealing multiple telomere-containing reads at SV breakpoints accompanied by coverage-based CN shifts (**Figure 6a, lower panels**). We also observed heterogeneity in repeat architecture among reads at the same locus, suggesting sub-clonal variability at TRC sites.

**Figure 6.**
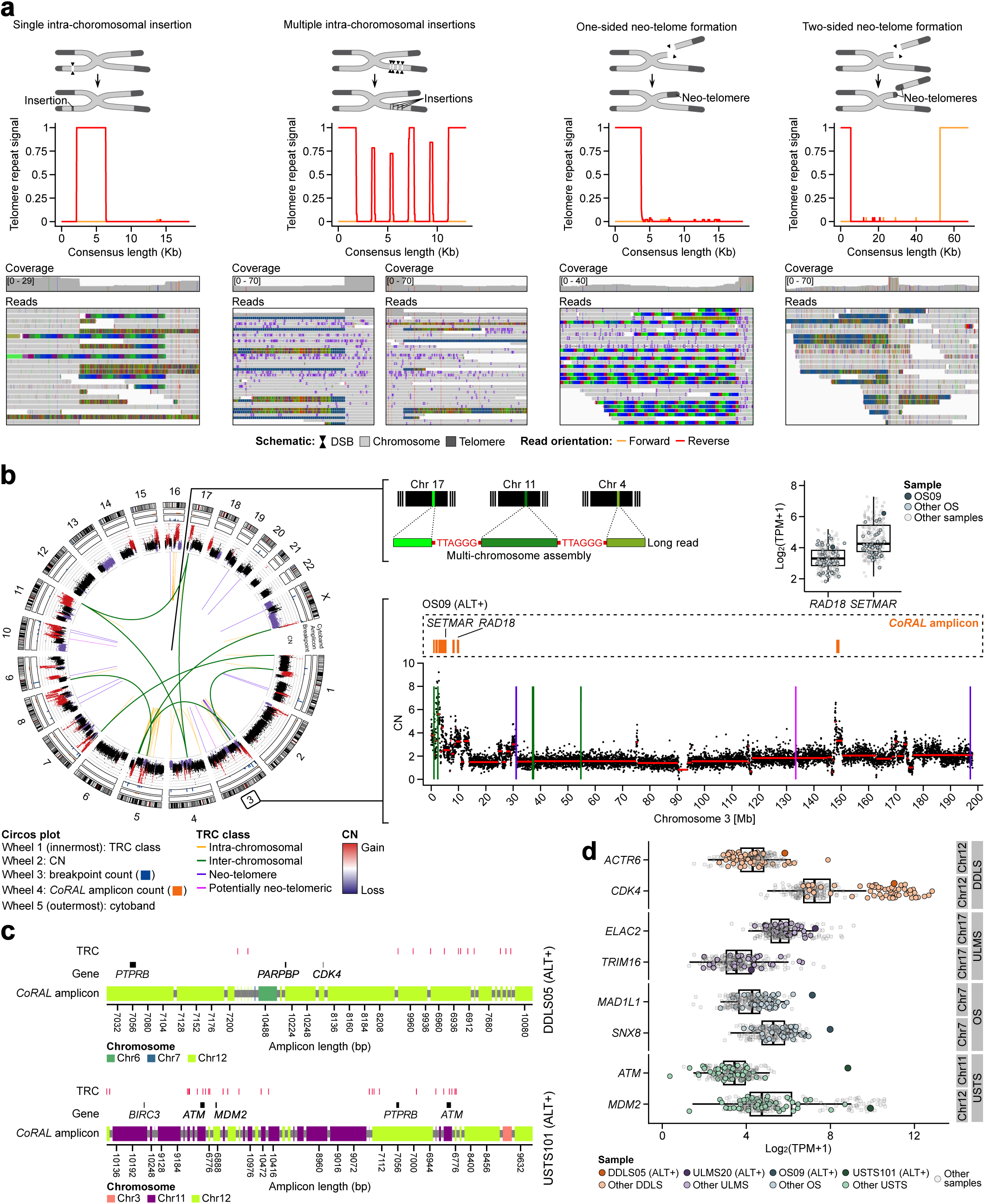
Telomere-mediated healing of DNA double-strand breaks occurs at higher rates in ALT+ complex karyotype sarcomas and converges with focal amplifications in highly rearranged genomic regions. (**a**) Representative telomeric repeat-containing consensus sequences detected by Nanopore sequencing across the eight CKS samples, supporting single and multiple intra-chromosomal insertions, as well as one-sided and two-sided neo-telomere formation. Upper panels show schematics of the inferred structural rearrangements. Middle panels depict the telomeric repeat signal along each consensus sequence, where a value of 1 denotes a sequence composed entirely of canonical telomeric repeats and the width reflects the length of the repeat tract. Lower panels show coverage profiles and read alignments that directly support the corresponding event. (**b**) Circos plot showing telomeric repeat clusters (TRCs) distributed across the genome of the ALT+ OS09 sample. From outermost to innermost, tracks show cytobands, CN profile, SV breakpoint count, and *CoRAL*-predicted amplicon count. Internal links denote intra- or inter-chromosomal, neo-telomeric, and potentially neo-telomeric TRCs. The right panel shows the CN profile of chromosome 3, including a chromothriptic region with co-occurring telomeric repeat-containing rearrangements and focal amplifications reconstructed as *CoRAL* amplicon. A long read supporting a multi-chromosomal assembly of several TRCs mapping to chromosomes 17, 11, and 4 was included to illustrate the complexity of these events. Boxplots indicate *RAD18* and *SETMAR* log2(TPM+1) expression in OS09 relative to other OS and CKS samples in our RNA-seq dataset (*n* = 260). (**c**) Representative *CoRAL* linear amplicons from the ALT+ DDLS05 and USTS01 samples, with colored segments indicating the chromosomal origin of individual amplicon fragments. Red marks denote TRCs. Selected cancer-associated loci are annotated. (**d**) Expression of selected genes located within recurrently amplified chromosomal regions reconstructed as linear amplicons by *CoRAL*. Each point represents a CKS sample. Darker colors indicate the ALT+ cases DDLS05, ULMS20, OS09, and USTS01, with each CKS subtype represented by a distinct color. Kb: kilobase, Mb: megabase, DSB: double-strand break, CN: copy number, TPM: transcripts per million, DDLS: dedifferentiated liposarcoma, ULMS: uterine leiomyosarcoma, OS: osteosarcoma, USTS: undifferentiated soft-tissue sarcoma.

### Telomere-mediated DSB healing is associated with complex interchromosomal rearrangements

Our consensus sequencing strategy distinguished simple single-breakpoint events with mappable flanking regions from complex rearrangements with clustered SV breakpoints around the TRCs, the latter generating prefixes and suffixes that mapped to multiple positions within the same chromosome and, strikingly, across different chromosomes (**Supplementary Figure 8h, Figure 6b**, **green line and insert**). As shown for OS in **Figure 6b** and USTS, DDLS and LMS samples in **Supplementary Figure 9** (**green line**) these multichromosome mappings associated with TRCs were observed almost exclusively in ALT+ tumors (**green line**, **Supplementary Figure 9**), implicating telomere-mediated repair in complex inter- and intra-chromosomal recombination events. Additionally, TRCs frequently co-occurred in the regions displaying high-level structural rearrangement and chromothripsis (**Figure 6b, Supplementary Figure 9, circos plot wheel 1-3**) consistent with our short-read TelomereHunter findings (Figure 5h), underscoring emergence of TRCs as ALT-associated telomere healing events stabilizing DNA breaks. Together, these observations indicate that telomere-mediated DSB healing in ALT+ CKS extends beyond simple chromosome-end repair to involve complex interchromosomal rearragenements. By stabilizing broken chromosomes, these ALT-associated healing events may protect ALT+ tumor cells from instability-induced crisis while simultaneously contributing to a higher-order layer of genomic complexity.

### Telomere repeat clusters are embedded in focal amplifications and ecDNA in ALT+ CKS, linking ALT to amplicon biology and oncogene overexpression

The structural instability of the genomes in cancer enables a further layer of genomic remodeling: the formation of focally amplified DNA elements that drive oncogene dosage independently of canonical chromosomal regulation. Such amplifications can adopt distinct configurations, including circular extrachromosomal DNA (ecDNA) and linear chromosomal amplicons^87–89^. To examine whether the structural instability observed in our CKS cohort is accompanied by such focally amplified architectures, we next characterized amplicon prevalence and configuration in our Oxford Nanopore sequencing cohort (n = 8) using *CoRAL*^90^.

*CoRAL*-resolved amplicons were more abundant in ALT+ ULMS, OS, and USTS samples relative to their ALT− counterparts, whereas DDLS showed the opposite pattern, with higher amplicon counts in ALT− cases (**Supplementary Figure 10, bottom left panel**). Aggregating predicted amplicon length per chromosome revealed subtype-specific patterns (**Supplementary Figure 10, right panel**). DDLS amplifications concentrated on chromosome 12 in both ALT subsets, with greater cumulative length in ALT+ cases (19 Mb vs. 11 Mb; **Supplementary Figure 10, top left panel**). ULMS amplicon burden peaked on chromosome 17 (up to 5 Mb), OS distributed broadly across chromosomes 7, 8, 9, and 17 (each exceeding 10 Mb), and USTS clustered on chromosomes 12 and 17 (also exceeding 10 Mb).

Next, we examined the prevalence of TRCs, genomic scars and gene loci prevalent in these amplicons. In ALT+ OS09 (**Figure 6b**), *CoRAL* detected multiple genome-wide amplicons (wheel 4), including a chromothriptic region on chromosome 3p with focal amplifications spanning *SETMAR*, an H3K36 methyltransferase implicated in NHEJ-mediated DSB repair^91^ and *RAD18*, an E3 ubiquitin ligase coordinating post-replication repair and DSB repair through homologous recombination^92^. Both genes overlapped with intra-chromosomal TRCs and neo-telomeres. *SETMAR* and *RAD18* expression in OS09 was elevated relative to cohort medians (n = 260; *SETMAR* log₂(TPM+1): 6.16 vs. 4.25; *RAD18*: 3.99 vs. 3.31), consistent with sample-specific focal amplification of these genes.

Two additional representative amplicons in ALT+ DDLS05 and USTS01 (**Figure 6c**) displayed combination of both, multi-chromosomal mapping joined by TRCs and overlapping *CoRAL* amplicons. Both contained segments from multiple chromosomes (3, 6, 7, 11, and 12, with chromosome 12 shared between cases) and included *ATM*, *PTPRB*, *CDK4*, and *MDM2*. Multiple TRCs were embedded within these amplicons. Across the cohort, expression of genes recurrently incorporated into amplicons, including *MAD1L1*, *SNX8* (chromosome 7), *ATM* (11), *ACTR6*, *CDK4*, *MDM2* (12), and *ELAC2*, *TRIM16* (17), was elevated in ALT+ tumors carrying the corresponding amplicons, exceeding both cohort medians and the expression of these genes in other tumors of the same subtype (**Figure 6d**). These data link focal amplification to increased expression of the amplified genes in ALT+ CKS.

### Amplicon architecture in CKS reflects subtype and TMM context

To resolve amplicon architecture in the larger WGS cohort, we applied *AmpliconArchitect*^93^ across DDLS, LMS, ULMS, OS, and USTS (**Supplementary Figure 11a**). In DDLS, ecDNA amplicons were enriched in ALT– tumors, while ALT+ DDLS amplicons were predominantly linear and complex-non-cyclic. This pattern is consistent with the established association between telomerase reactivation, driven in DDLS by recurrent *TRIO::TERT* rearrangements at 5p15^94^, and focal circular amplification of the 12q13–15 *MDM2*/*CDK4* core^88^. In OS and USTS, amplifications distributed broadly across linear and complex-non-cyclic classes in both ALT subsets, consistent with the chromothripsis-prone genomes of these subtypes^34^. ULMS showed the opposite pattern to DDLS, with complex-non-cyclic amplicons enriched in ALT+ tumors. BFB amplicons were rare across subtypes regardless of ALT status.

We then mapped the chromosomal distribution of protein-coding genes carried on ecDNA, stratified by ALT status and CKS subtype (**Supplementary Figure 11b**). Most ecDNA gene cargo came from DDLS and OS, the subtypes with the highest proportion of ecDNA-positive tumors, and clustered on chromosomes 1, 6, and 12. DDLS ecDNA-associated genes mapped to chromosome 12 and were recurrent in more than 30 tumors; genes identified in other subtypes occurred in only one or two cases. *TERT*, *CDK4*, *JUN*, *MDM2*, *HMGA2*, and *ROS1* appeared on ecDNA in both ALT– and ALT+ tumors. Other genes showed subtype- or ALT-restricted distributions: *MDM4*, *SETDB1*, *JAK2*, and *SMARCA4* in DDLS; *METTL3*, *SALL2*, and *NKX2-1* in LMS; *LATS2*, *SKA3*, *ITCH*, and *SNTA1* in ULMS; *HMGA1*, *KDM2A*, *NCOR1*, *TOP3A*, and *PATZ1* in OS; and *CCNE1*, *HDAC1*, *CDC5L*, *ATM*, and *TRIM8* in USTS. Taken together, we observe that ecDNA formation in CKS varies by subtype rather than by ALT status, and the gene content of ecDNA differs between ALT+ and ALT– tumors within each subtype.

## Discussion

We performed a pan-CKS analysis to characterize how TMM status associates with the genomic and transcriptomic state of chromosomally unstable sarcomas. By comparative interrogation of ALT+ and ALT– tumors of four CKS subtypes, we found that ALT activity in CKS is associated with enriched hallmarks of genomic instability, and a transcriptional program centered on DNA damage response and mitotic control, and uncovered TRCs and multi-chromosome mappings on chromosomal and extra-chromosomal DNA as footprints of ALT-associated “telomere healing” process.

All CKS subtypes investigated shared TMM-associated distinction in genomic and transcriptomic landscape, but also displayed subtype-specific differences in specific genomic features, such as focal CNA patterns or ecDNA type and cargo. For example, *SMARCAL1* loss on 2q was observed in ALT+ DDLS, LMS, ULMS, and USTS, but not in ALT+ OS (**Figure 3a, Supplementary Figure 6a**). This is consistent with the recently established requirement for SMARCAL1 in ALT+ OS to mitigate BLM-driven telomeric replication stress^95^. This subtype-specific dichotomy suggests that ALT+ CKS tumors may follow two evolutionary trajectories: one in which *SMARCAL1* loss facilitates ALT activity^45,46^, and another in which *SMARCAL1* retention acts as an essential brake on ALT-associated replication stress, potentially representing a therapeutically exploitable vulnerability in ALT+ OS. Considering broader differences between subtypes, ALT+ ULMS stood apart from the closely related LMS and other CKS subtypes as the most structurally rearranged ALT+ subset, combining a deletion-dominant CNA profile, the highest chromothripsis frequency in CKS (95%), and complex-non-cyclic amplicon enrichment. These differences indicate that anatomical site and lineage context, potentially determined by accessible chromatin, may play a role how ALT shapes CKS genomes.

ALT+ CKS were enriched for two HRD readouts: elevated scarHRD scores (genomic scarring) and elevated SBS3 burden (a mutational signature). This was unexpected, since ALT requires functional homologous recombination for RAD51/RAD52-mediated telomere extension^96,97^ and deleterious *BRCA1/2* alterations were rare in our cohort, in line with previous studies, ruling out canonical HR loss as the source. A likely explanation could be chronic telomeric recombination in ALT+ tumors that may generate HRD-like genomic scars despite retained HR machinery. Under sustained replication stress and frequent TP53 loss (**Figure 4f**), these scars may accumulate rather than triggering checkpoint arrest. Thus, BRCA1/2-driven HRD and ALT-associated scarring may produce similar genomic patterns through distinct biological mechanisms. This, however, does not explain SBS3 burden in ALT+ CKS. Whether HRD readouts in ALT+ CKS predict PARP inhibitor response, or reflect past recombination without a targetable vulnerability will require functional testing in ALT+ sarcoma models. The investigations around ALT status as a biomarker for response to PARP inhibition are clinically relevant and urgent, given recently reported moderate responses to PARP inhibition in unstratified sarcoma^98,99^.

Beyond PARP inhibitors, our transcriptomic data point to broader dependencies in ALT+ CKS. The coordinated upregulation of DNA damage response and mitotic regulatory programs identifies ATR–CHK1 signaling and mitotic kinases (AURKA, AURKB, PLK1) as candidate vulnerabilities, supported by existing preclinical and clinical-stage activity of ATR inhibitors in ALT-positive cancers^58,100^. Because this transcriptional state was subtype-agnostic in our data, it offers a strategy not constrained by the lineage-specific oncogene patterns that currently limit targeted therapy in sarcoma. Our data point towards DDR-targeting agents and mitotic kinase inhibitors as a promising combination worth exploring preclinically in ALT+ sarcoma models.

Long-read sequencing identified intra-chromosomal telomeric repeat clusters and neo-telomere formation at SV breakpoints across rearrangement-prone regions of ALT+ CKS, with multichromosome assemblies stabilized by telomeric repeats representing a novel mode of structural complexity. These findings extend recent observations of telomere insertions as TMM footprints^13^, neotelomere formation in ALT-positive neuroblastoma^18^, and ATR-mediated suppression of telomerase-driven neotelomere formation at DSBs^82^. Together they suggest that ALT-associated recombination provides an additional DNA repair-like capacity allowing broken chromosomes to be stabilized rather than triggering cell death through unresolved DSBs, BFB cycles, or end-to-end fusion catastrophe. By stabilizing massively rearranged genomes, telomere-mediated healing may provide a substrate for new oncogenic configurations through ecDNA formation, multichromosome amplicon assembly, and reshuffling of tumor suppressor and oncogene loci, which may result in genome diversification leading to treatment resistance, which could contribute to poor prognosis of ALT+ sarcoma^16^. The convergence of TRCs with chromothripsis-derived ecDNA and complex linear amplicons in ALT+ CKS, together with their oncogene-rich gene cargo, supports this view and identifies telomere-mediated healing as a previously underappreciated driver of structural complexity in cancer. Many established sarcoma therapies, including doxorubicin, ifosfamide, etoposide, trabectedin, and radiotherapy, exert their cytotoxic effects through DSB induction. In light of our findings, in an ALT-proficient cell, if sublethal DSB induction selects for cells that have successfully invoked telomere-mediated healing, treatment may inadvertently contribute to tumor aggressiveness, accelerated clonal evolution, and emergence of resistant clones. This concept warrants experimental testing in ALT+ sarcoma models exposed to standard cytotoxic agents, as well as in-depth analysis of longitudinal tumor samples from patients undergone treatment with these agents.

Several limitations of this study warrant consideration. First, the C-circle assay, although considered the “gold standard” for ALT detection^10^, provides a single-timepoint readout of ALT activity and may not fully capture intratumoral TMM heterogeneity, particularly in tumors where telomerase- and ALT-dependent populations coexist. Second, our long-read sequencing cohort of eight paired ALT−/ALT+ samples across four CKS subtypes was designed to resolve the nucleotide-level architecture of complex telomere-healing events, including multichromosome assemblies and neo-telomere formation. However, population-level frequencies of these events across the broader CKS landscape will require larger long-read cohorts. Third, our ecDNA analysis relied primarily on short-reads, which has reduced resolution in repetitive and telomere-proximal regions and may not fully capture ecDNA structures embedded within highly rearranged ALT+ genomes. Longitudinal sampling and expanded long-read sequencing will be needed to follow TMM dynamics over time and to clarify how telomere-mediated healing contributes to tumor evolution and therapeutic resistance in CKS.

Our findings suggest that ALT activity in CKS is associated not only with telomere maintenance, but with broader differences in genomic instability, transcriptional state, and structural complexity that have potential therapeutic implications. ALT status warrants consideration as a stratifying variable in future CKS studies and model systems. Integrating ALT status into standard CKS profiling, expanding availability of preclinical CKS models will allow functional validation of biomarkers and therapeutic strategies proposed in this work, accelerating path towards clinical evaluation and improved clinical care for sarcoma patients.

## Materials and Methods

### Patient samples

Patients provided written informed consent for multi-layered molecular profiling of tumor tissue and, for DNA sequencing, a matched blood sample, as well as for the longitudinal collection of clinical information as part of the DKFZ/NCT/DKTK MASTER registry trial^20^ (ClinicalTrials.gov ID: NCT05852522). The study was approved by the Ethics Committee of Heidelberg University (protocol no. S-206/2011) and conducted in accordance with the Declaration of Helsinki^101^. Biological curation and clinical annotation of molecular profiles were performed as previously described, and a multi-institutional MTB involving treating physicians provided recommendations for clinical management^102^. Participants were not financially compensated for taking part in the study. The final cohort included 776 patients diagnosed with bone or soft-tissue sarcomas, including both treatment-naive and previously treated cases. The complex karyotype sarcoma (CKS) cohort used for comparative analyses of genomic and transcriptomic features between tumors comprised dedifferentiated liposarcomas (DDLS), leiomyosarcomas (LMS), uterine leiomyosarcomas (ULMS), osteosarcomas (OS), and undifferentiated soft-tissue sarcomas (USTS; **Supplementary Table 1**)

### Sample processing and sequencing

Tumor tissue was obtained via surgical resection or biopsy and immediately snap-frozen. Board-certified pathologists reviewed cryosections to assess tumor cell content and the extent of necrosis. All specimen processing was performed under accredited conditions at the NCT/DKFZ Heidelberg Sample Processing Laboratory. DNA and RNA were extracted from fresh-frozen tumor tissue using the AllPrep DNA/RNA/miRNA Universal Kit (Qiagen) and the QIAamp DNA Mini Kit (Qiagen). Quality and concentration of nucleic acids were assessed using a Qubit Fluorometer (Life Technologies) and a 4200 or 2200 TapeStation System (Agilent). WES, WGS, and RNASeq data were generated under accredited conditions at the DKFZ Genomics and Proteomics Core Facility. For WES and WGS, a minimum coverage of 60X was ensured for both tumor and matched control samples. For RNASeq, a minimum of 70 million reads was generated per sample. Additional technical details have been described previously^103,104^. Raw sequencing data underwent preliminary processing and quality control using automated DKFZ bioinformatics workflows^105^.

### C-circle assay

C-circle (CC) assays were performed on tumor-derived genomic DNA (gDNA) following an adapted version of the original protocol^9^. Before amplification, 30 ng of gDNA was digested with *HinfI* and *RsaI* restriction enzymes (1 U/µL each; New England Biolabs) in CutSmart buffer supplemented with RNase A (10 µg/mL; Roche) at 37°C for 1 h to remove RNA, linearize chromosomal DNA, and facilitate selective amplification of circular telomeric DNA. Each sample was set up in triplicate, with and without phi29 DNA polymerase (New England Biolabs). The amplification master mix contained 0.2 mg/mL BSA (New England Biolabs), 0.1% Tween-20 (Sigma-Aldrich), 1 mM each of dATP, dGTP, and dTTP, and 7.5 U phi29 DNA polymerase in 2X phi29 reaction buffer. Reactions were incubated at 30°C for 8 h, followed by enzyme inactivation at 65°C for 20 min. Amplified products were mixed with 2X SSC buffer (0.3 M sodium chloride, 0.03 M sodium citrate, pH 7) and transferred onto a positively charged Roti-Nylon plus 0.45 µm membrane (Carl Roth) using a Bio-Dot microfiltration apparatus (Bio-Rad, Cat. no. 170-6545). Membranes were left to dry for 10 min at room temperature, baked at 120°C for 20 min to crosslink DNA, and rehydrated in 2X SSC. CC amplification products were detected using the TeloTAGG Telomere Length Assay Kit (Roche). Membranes were pre-hybridized for 1 h at 42°C in DIG Easy Hyb solution, followed by overnight incubation at 42°C with a digoxigenin-labelled telomere probe (30 nM) on a rotating platform. After hybridization, membranes were washed twice for 5 min at room temperature in 2X SSC containing 0.1% SDS, followed by two additional washes in 0.2X SSC supplemented with 0.1% SDS for 20 min at 50°C. Membranes were subsequently rinsed for 5 min in washing buffer and blocked for 30 min before incubation with Anti-DIG-AP Fab fragments (75 mU/mL; Roche) for 60 min. Excess antibody was removed by two 15-min washes in washing buffer. Membranes were then equilibrated in detection buffer for 5 min and incubated with CDP-Star chemiluminescent substrate solution (Roche) for 10 min before signal acquisition on a ChemiDoc MP imaging system (Bio-Rad). Quantification of CC signals was performed in *ImageLab* (v6.0.34). Dot intensities were determined from uniformly sized circular regions, after subtraction of local background and the corresponding no-phi29 control signal for each sample. Raw CC levels were calculated by averaging technical replicates. Each membrane included a U2OS gDNA dilution series (4, 8, 16, 32, and 64 ng input gDNA per reaction) with matched no-amplification controls to assess assay linearity and generate a standard curve. CC levels were then normalized to the signal corresponding to 20 ng U2OS gDNA, calculated from the standard curve to account for inter-membrane variation. Each membrane additionally included U2OS as ALT+ control and HeLa as ALT-control. Samples showing CC levels ≥ 0.1 arbitrary units (AU) relative to U2OS were classified as CC-positive (ALT+), whereas samples below this cutoff were considered CC-negative (ALT-).

### Whole-exome sequencing and variant calling

A detailed list of pseudonymized CKS patients with WES data (*n* = 190) is provided in **Supplementary Table 1**. Sequencing reads were mapped to the 1000 Genomes Phase 2 assembly of the Genome Reference Consortium human genome (build 37, version hs37d5 or 1KGRef_PhiX) using *BWA-mem* (v0.7.15). Duplicate reads were marked and removed using Picard’s *MarkDuplicates* tool (v1.125; https://github.com/broadinstitute/picard). The GATK *Base Quality Score Recalibration (BQSR)* algorithm (v4.2.5.0; https://github.com/broadinstitute/gatk) was applied to enhance base quality scores assigned during sequencing. Germline SNVs and indels were identified using the GATK *HaplotypeCaller* tool (v4.2.0) following the recommended filtering criteria. Germline variants were then annotated for their coding effects and functional impact with *PAVE* (v1.6; https://github.com/hartwigmedical/hmftools/tree/master/pave), integrating annotations from the *ClinVar* database (May 2023 release). Somatic SNVs and indels were identified through the DKFZ *One Touch Pipeline (OTP)* as previously described (). Somatic SNVs were detected using *SAMtools/bcftools* (v0.1.19, https://github.com/DKFZ-ODCF/SNVCallingWorkflow). Each variant was assigned an initial confidence score of 10, which was subsequently adjusted based on overlaps with repetitive regions, DAC-blacklisted regions, DUKE-excluded regions, self-chain sequences, and segmental duplications, as defined by the *ENCODE* project^106^. Confidence scores were also reduced for variants exhibiting PCR or sequenci^106^ng strand biases. Somatic indels were called using *Platypus*^107^ (v0.8.4), with confidence scores calculated through internal filtering criteria. Both SNVs and indels with a confidence score of 8 or above were annotated with *ANNOVAR*^108^ (October 2019 release) and *GENCODE* (v19) gene models. The resulting variants were cross-referenced against the *dbSNP* (build 141) and *1000 Genomes Project* repositories. Variants predicted to alter protein-coding sequences were classified as synonymous or non-synonymous substitutions, stop-gain or stop-loss events, splice-site alterations, and frameshift or in-frame indels.

### Whole genome sequencing and variant calling

**Supplementary Table 1** includes all pseudonymized CKS patients with WGS data and the corresponding sequencing quality metrics (*n* = 148). Sequencing reads were aligned to the 1000 Genomes Phase 2 assembly of the Genome Reference Consortium human genome (build 37, version hs37d5 or 1KGRef_PhiX) using *BWA-mem*^109^ (v0.7.15). BAM files were sorted with *SAMtools*^110^ (v0.1.19), followed by duplicate read removal and base quality score recalibration as previously described for WES data. Somatic and germline SNVs and indels were identified and filtered using *SAGE* (v3.3; https://github.com/hartwigmedical/hmftools/tree/master/sage) with default parameters. The coding effects and functional impact of these variants were annotated with *PAVE* (v1.6, https://github.com/hartwigmedical/hmftools/tree/master/pave). Somatic structural variants (SVs) were detected with *GRIDSS*^111^ (v2.13.2) with default settings, and high-confidence calls were refined using *GRIPSS* (v2.3; https://github.com/hartwigmedical/hmftools/tree/master/gripss).

### Copy number alteration analysis

Copy number (CN) profiles of CKS samples, incorporating tumor purity and ploidy estimates, were determined using *ASCAT*^112^ (v3.1.2) and *Sequenza*^113^(v3.0.0) for WES data with standard parameters, and *PURPLE* (v3.8.4; https://github.com/hartwigmedical/hmftools/tree/master/purple) for WGS data. Purity and ploidy estimates from *ASCAT* and *Sequenza* were concordant in the majority of samples. For samples with multiple solutions, different combinations of purity and ploidy values were systematically evaluated, and the solution best supported by the data was selected manually. The final purity and ploidy estimates were used to calculate total and allele-specific CN for each genomic segment. CN segments were classified into three heterozygosity states: heterozygous segments (A > 0, B > 0, where A and B represent the major and minor alleles, respectively), LOH segments (A > 0, B = 0, indicating loss of one parental allele), and homozygous deletions (A = 0, B = 0, reflecting complete loss of both parental alleles). Segments were further categorized using a threshold of 0.7 to identify duplications, amplifications, and deletions based on deviation from the expected ploidy, while regions within ±0.3 of the ploidy value were classified as CN neutral^114^. For male samples, gains and losses on chromosomes X and Y were evaluated relative to half the ploidy.

CN segments derived from *PURPLE*-based CNA calling in CKS tumors with WGS data were adjusted to ensure compatibility with 1-based genomic coordinates from the UCSC Genome Browser human reference genome (GRCh37/hg19). Log2-transformed relative CN values were converted to absolute CN using the formula *absolute CN = 2^segmean+1^*. Segments with absolute CN exceeding 500 were excluded as likely reflecting segmentation errors, low-coverage regions, and/or inaccurate purity/ploidy estimates. CN frequencies across chromosomes were calculated separately for ALT- and ALT+ CKS tumors and visualized using the *GenVisR* R package^115^ (v1.36.0), applying thresholds of ≥ 4.25 for gains and ≤ 1.75 for losses to account for the pronounced genomic instability characteristic of CKS tumors.

For segment-level analyses, CN segments were classified by size into focal (0.02-10 Mb) and broad (> 10 Mb) categories based on the empirical distribution of segment sizes observed across the WGS cohort (**Supplementary Figure 2a-b**). Segments overlapping centromeres were assigned to the p- or q-arm based on the fraction of their length overlapping each arm, with ≥ 50% overlap required for arm assignment. Segments smaller than 0.02 Mb or with more than 50% of their length falling within centromere boundaries were excluded from all downstream analyses.

For arm-level summarization of broad and focal CNAs based on ALT status across CKS entities, purity-corrected CN segments were independently processed using *CNApp*^51^(v1.1.0). *CNApp* re-segments input data to correct for amplitude divergence due to technical variability and generates arm-level log2 CN ratio matrices. Within this framework, broad CNAs were defined as alterations spanning ≥ 50% of a chromosomal arm or ≥ 90% of an entire chromosome, whereas focal CNAs were defined as alterations affecting < 50% of a chromosomal arm. Sex chromosomes were excluded from all arm-level analyses to avoid gender-specific CN biases unrelated to tumor biology. Mean log2 CN ratios per chromosomal arm were calculated separately for ALT- and ALT+ samples within each CKS subtype. To quantify differences in arm-level CN signal between ALT-and ALT+ samples within each entity, pseudocount-adjusted ratios were computed as *(CNALT++1)/(CNALT-+1)*. Arms with ratios ≥ 1.3 or ≤ 0.7 (corresponding to ≥ 30% relative difference) were annotated with single arrows, and arms with ratios ≥ 1.5 or ≤ 0.5 (≥ 50% relative difference) with double arrows, with upward and downward arrows indicating higher and lower CN signal in ALT+ samples, respectively. CN patterns were visualized using the *pheatmap* R package (v1.0.12; https://CRAN.R-project.org/package=pheatmap) without row scaling to preserve the actual magnitude of CN ratios across subtypes and ALT status groups.

### *ATRX*, *DAXX*, and *TERT* alteration analysis

Somatic and germline mutations, broad and focal CNAs, and SVs in *ATRX*, *DAXX*, and *TERT* were extracted from WGS data available for the CKS cohort as described above. Only variants passing all quality filters and not assigned to a low-confidence tier were retained. Variants were left-aligned and multiallelic sites were split using *SAMtools/bcftools* ^110^(v1.16) against the GRCh37/hs37d5 reference genome. Functional annotation was performed using Ensembl^116^ *VEP* (v110.1), with *CADD* scores computed using v1.7 for SNVs and v1.6 for indels^117^, and pathogenicity scores for missense variants were obtained using *REVEL*^118^(v1.3). Alterations were manually verified using the Integrative Genomics Viewer^119^ (IGV) and integrated with matched RNA-seq-based gene expression data quantified as log2(TPM+1). Expression levels were evaluated relative to gene-specific wild-type reference thresholds, defined as the 5^th^ percentile for reduced expression and the 75^th^ percentile for increased expression. Given the low and heterogeneous expression of *TERT* in cancer, together with the limited sensitivity of bulk RNA-seq for *TERT* transcript detection, expression-based thresholds were not applied for *TERT* classification. Each sample was assigned to one of the following functionally informed alteration categories: loss of heterozygosity (LOH), deleterious (encompassing loss-of-function variants, missense variants with REVEL score ≥ 0.5, variants with CADD PHRED score ≥ 30, homozygous deletions, and/or other loss events supported by reduced expression), low-level gain (duplications without increased expression), high-level gain (duplication and/or amplification events, with or without increased expression), *TERT*-activating (characterized by known activating promoter mutations and/or enhancer hijacking events), non-deleterious, and wild-type. To screen for *TERT* promoter mutations, all CKS tumors were interrogated for the canonical C228T and C250T activating substitutions^120^, and variant allele frequencies were compared between matched tumor and control samples to confirm somatic origin. To identify potential *TERT* enhancer hijacking events, SVs with at least one breakpoint located within 1 Mb upstream of the *TERT* transcription start site (TSS) were extracted from WGS data, as described previously^13^. The juxtaposed genomic coordinates of each SV were intersected with 65,950 predicted super-enhancers from the *dbSUPER* database^29^. Overlaps were classified as direct when the juxtaposed position fell within a super-enhancer interval, or indirect when a super-enhancer was located within 1 Mb of the juxtaposed position. Tumors were considered to harbor a potential enhancer hijacking event when a SV breakpoint and its rearranged enhancer site were both located within 20 Kb of the *TERT* TSS.

### Chromothripsis analysis

Chromothripsis events across the CKS cohort were identified from WGS-derived CN and SV data using *ShatterSeek* ^35^(v1.1). For each sample, SV calls included breakpoint coordinates, SV type (deletions, duplications, head-to-head inversions, and tail-to-tail inversions), and strand orientations, while CN data included chromosomal segment boundaries and total CN. Chromothripsis calls were classified as high-confidence when a chromosomal region harbored at least six interleaved intrachromosomal SVs, seven or more contiguous CN segments oscillating between two states, a significant fragment joins test, and either a significant chromosomal breakpoint enrichment or exponential distribution of breakpoints test (*P* < 0.05). Low-confidence calls required the same SV and fragment join criteria but with three to six oscillating CN segments, or fulfilled the CN criterion without meeting both statistical tests. Events involving chromosome Y were removed before analysis. To visualize the genomic architecture of chromothripsis-affected chromosomes, SV and CN profiles of representative samples were displayed using *ReConPlot* ^39^(v0.2), which integrates three panels per case: a chromosome ideogram, total and minor CN profiles, and SV rearrangements, where arcs connect breakpoints within the displayed chromosomal region and vertical lines with overhangs indicate SVs with only one breakpoint within the displayed region.

### Mutational signature analysis

Somatic single-base substitution (SBS) mutational signatures were extracted using feature matrices derived from WGS data generated for CKS tumors, stratified by ALT status and analyzed independently. Feature matrices were generated using *SigProfilerMatrixGenerator* ^121^(v1.2.20;) before signature decomposition. Signature decomposition was performed using *SigProfilerExtractor*^122^(v1.1.23) with the human reference genome GRCh37/hg19, testing between 1 and 10 signatures and running 100 non-negative matrix factorization replicates to ensure robustness of inference. Extracted signatures were matched against the COSMIC SBS reference catalogue^123^ (v3.4) following established procedures^124^. Prior to analysis, outlier samples were excluded if their total mutation burden exceeded five times the cohort median or if they were flagged for sequencing artifacts. Only SBS signatures detected in at least six samples were retained for downstream interpretation and visualization. Proposed etiologies and abbreviations for each retained signature are indicated in **Figure 2g**.

### HRD quantification

The *scarHRD* ^44^ score. The score is calculated as the unweighted sum of three genomic instability metrics: loss of heterozygosity (HRD-LOH; defined as LOH regions exceeding 15 Mb that do not span entire chromosomes), telomeric allelic imbalance (TAI; allelic imbalances extending to the telomeric end of a chromosome), and large-scale state transitions (LST; chromosomal breaks between adjacent segments of at least 10 Mb separated by no more than 3 Mb). The *scarHRD* score was computed using the *scarHRD* R package^42^ (v0.1.1) on combined WES and WGS datasets (*n* = 324), using *ASCAT*- and *PURPLE*-derived segmentation data as input, respectively. ALT- and ALT+ CKS samples with a *scarHRD* score ≥ 42 were classified as homologous recombination deficient (HRD+). To orthogonally validate *scarHRD* results, HRD probability was independently quantified using *HRProfiler* (v0.0.1). *HRProfiler* is a machine learning approach based on a linear kernel support vector machine trained on six mutational features derived from SBS, indel, and CN mutational contexts. Specifically, the tool integrates features enriched in HRD tumors, including LOH events spanning 1-40 Mb, deletions ≥ 5 bp at microhomology sequences, and heterozygous CN amplifications of 10-40 Mb, alongside features enriched in homologous recombination-proficient (HRP) tumors, including C>T substitutions at NpCpG contexts and large heterozygous segments exceeding 40 Mb. Somatic SNV/indel VCF files and *ASCAT*-derived CN segmentation files were used as input for both WES- and WGS-based analyses. Sequencing-specific pre-trained breast cancer models were then applied to the corresponding data modalities across the combined CKS cohort (*n* = 289), with all features normalized using *StandardScaler* before classification. Samples with an HRD probability ≥ 0.42 were classified as HRD+, consistent with the *scarHRD* score threshold of 42.

### RNA sequencing and gene expression quantification

RNASeq reads were aligned to the human reference genome (GRCh37/1KGRef_PhiX) using *STAR* ^125^(v2.7.10a) in two-pass mode, with *GENCODE* (v19) gene models. Duplicate reads were marked using *Sambamba* ^126^(v0.6.5). Gene-level read counts and TPM values were quantified using *FeatureCounts* ^127^(*Subread* v1.6.5), with genes on chromosomes X, Y, and MT and rRNA/tRNA genes excluded from library size estimation. All workflow parameters followed the DKFZ/ODCF RNASeq workflow (Release 4.0.0; https://github.com/DKFZ-ODCF/RNAseqWorkflow). TPM values were log2-transformed after addition of a pseudocount of 1 for downstream analyses.

### Differential gene expression and pathway analysis

To assess whether ALT status drives transcriptomic variation independently of sarcoma subtype, partial least squares discriminant analysis (PLS-DA) was performed separately within each CKS entity to identify latent components maximally separating ALT- and ALT+ samples, using gene expression profiles normalized with *DESeq2* ^128^(v1.44.0). Batch-corrected data were not used as input, as preliminary testing indicated overfitting of the resulting models. Genes with zero variance across all samples were excluded before analysis. An initial PLS-DA model retaining up to 10 components was fitted using the *plsda* function from the *mixOmics* R package^129^ (v6.28.0), and the optimal number of components was determined by 3-fold cross-validation repeated 50 times using the *perf* function, minimizing the balanced error rate. Final models were fitted with the optimized number of components determined for each entity (DDLS: 2, LMS: 3, ULMS: 3, OS: 2, and USTS: 2). Results were visualized as scatter plots of individual samples projected onto the first two PLS-DA components, with 95% confidence ellipses drawn per ALT status group.

Differential gene expression (DGE) analysis was performed between ALT- (*n* = 142) and ALT+ (*n* = 118) CKS transcriptomes using the *limma-voom* workflow^57,130^. Raw gene counts were first organized into a DGEList object using the *edgeR* R package^131^ (v4.2.2). Low-count genes were removed using the *filterByExpr* function, requiring a minimum count of 50 in at least 6 samples per group. Library sizes were normalized using the trimmed mean of M-values (TMM) method via *calcNormFactors*. *Voom* transformation was applied to estimate the mean-variance relationship of log2-transformed counts per million and assign precision weights to each observation. A design matrix was constructed with ALT status as the main factor and sequencing processing date as a covariate to adjust for batch effects across sequencing runs. Linear models were fitted to each gene using *limma* (v3.60.6) and pairwise contrasts between ALT- and ALT+ groups were tested using empirical Bayes moderation of standard errors. In total, 16,743 protein-coding genes were analyzed, of which 1,677 were differentially expressed between ALT- and ALT+ tumors (|log2fold-change| ≥ 0.6 and adjusted *P* ≤ 0.01; Benjamini-Hochberg correction). Among these, 888 and 789 genes were significantly up- and down-regulated in ALT+ CKS tumors, respectively, and are noted in **Supplementary Table 1**.

Gene set enrichment analysis (GSEA) was performed on DEGs ranked by log2(fold-change) using the *genekitr* R package ^132^(v1.2.8), querying gene sets from the Hallmark, Reactome, WikiPathways, KEGG, Gene Ontology Biological Process (GOBP), and Gene Ontology Molecular Function (GOMF) collections from the *MSigDB* database^133^. Multiple testing correction was applied using the Benjamini-Hochberg method. Functional relationships among enriched gene sets were examined using semantic similarity-based network analysis implemented within *genekitr*, using the Fruchterman-Reingold layout algorithm and Wang similarity method. For cytoband-level analysis, normalized enrichment score (NES) values from the *MSigDB* C1 positional gene set collection were plotted against -log10(FDR), with dot size proportional to the number of DEGs per cytoband and dot color reflecting NES magnitude. Pathway activity was inferred using *decoupleR* ^63^(v2.10.0) by testing gene-level t-statistics from the *limma-voom* model against the *PROGENy* regulatory network^64^ (top 500 target genes per pathway). Activity scores were estimated using a multivariate linear model, with a minimum gene set size of 5.

### Fusion gene analysis

Somatic gene fusions were detected from RNASeq data using *Arriba* ^134^(v1.2.0), which employs a single-pass alignment strategy based on *STAR* to extract chimeric reads, followed by a multi-step filtering framework combining blacklist-based filters to remove alignment artifacts and whitelist-based filters to retain true fusion events. Blacklist-based filters discard candidates supported by reads with homopolymers, tandem repeats, or excessive mismatches, as well as candidates between homologous genes or with insufficient supporting reads relative to background noise levels. Whitelist-based filters retain candidates with breakpoints at annotated splice sites or supported by SVs from matched WGS data. Each predicted fusion was assigned one of three confidence classes (low, medium, or high) based on the number and balance of supporting split reads and discordant mates, the event type, breakpoint distance, presence of corroborating events, and whether breakpoints were intragenic. Only high-confidence fusion calls were retained for downstream analyses.

### Analysis of telomere content, telomere variant repeats, and telomeric insertions

Telomere content from WES and WGS datasets was determined using *TelomereHunter* ^76^(v1.1.0). Telomeric reads were extracted by looking for sequences containing at least six non-consecutive occurrences of four of the most common telomeric repeat variants: TTAGGG, TCAGGG, TGAGGG, and TTGGGG. For downstream analysis, only unmapped reads or those with low mapping confidence (mapping quality < 8) were included, as they are more likely to originate from telomeric regions. Telomere content was quantified by normalizing the number of telomeric reads to the total number of reads in the sample with a GC content between 48% and 52%.

To identify telomere variant repeats (TVRs), hexamer motifs following the pattern NNNGGG were searched within the extracted telomeric reads. Only hexamers in which all bases had a minimum Phred quality score of 20 were considered. The 18 base pairs immediately upstream and downstream of each detected TVR were retrieved to define the surrounding sequence context. TVRs flanked by canonical telomeric repeats ((TTAGGG)3-NNNGGG-(TTAGGG)3), referred to as “singletons”, were selected for further evaluation. TVR frequencies were calculated by normalizing absolute counts to the total read count per sample. The expected singleton counts in an arbitrary context were calculated as log2 ratio of telomere content between tumor and control samples.

To identify telomeric repeat loci, defined as insertions of telomeric sequences into non-telomeric regions of the genome, we searched for tumor-specific discordant paired-end reads. These were characterized by one read containing telomeric repeats and the other uniquely aligning to the genome outside of telomeres with high confidence (mapping quality > 30). Candidate regions were identified in 1 kb windows containing at least four discordant read pairs in the tumor sample and none in the matched control. To pinpoint the exact insertion (junction) sites, we required a minimum of three supporting split reads spanning the insertion breakpoint. These split reads had to include at least one TTAGGG repeat to confirm the presence of the telomeric sequence. To avoid false positives due to common artifacts or structural variations in the population, regions showing discordant pairs in 15 or more control samples were excluded. All candidate loci were manually inspected using IGV to validate telomeric insertions and exclude residual artifacts. Across the sarcoma cohort, telomeric insertions were identified in 96 samples, with 2,026 of 2,248 candidate events retained following manual IGV curation. Insertions were further classified based on orientation: an insertion was considered two-sided if a second telomeric repeat locus with opposite orientation was found within 10 kb downstream on the same chromosome, sharing the same repeat motif on the forward strand. All other events were classified as one-sided. The full detection pipeline is available at https://github.com/linasieverling/TelomereRepeatLoci.

### Telomere fusion analysis

Telomere fusions (TFs) were detected using *TelFusDetector* ^14^(v1) applied to WGS data from the CKS cohort using default parameters. Briefly, the algorithm identifies reads containing at least two consecutive TTAGGG and two consecutive CCCTAA repeats, allowing up to two mismatches to capture TVRs, and classifies candidate events as inward, outward, or in-out based on the relative orientation of the telomeric repeat blocks. TFs mapping to the chromosome 9 endogenous fusion locus, poorly characterized genomic loci, and mitochondrial sequences were excluded. The TF rate for each sample was defined as the number of TFs normalized to 1X genome coverage and corrected for sequencing depth, read length, and tumor purity (derived using *ASCAT*). In total, 4,545 TFs were detected across 123 of 148 CKS samples, after excluding three cases with exceptionally high TF counts (> 400). To enable high-confidence assignment of TF breakpoints to specific chromosome ends, TF events were re-annotated using the anchored-mate framework in *TelFusDetector*, whereby a mate was considered anchored if it mapped to a subtelomeric region with mapping quality (MQ) ≥ 10. Events in which both supporting reads were anchored to subtelomeric regions were used to assign both breakpoint positions to specific chromosome ends, with each fusion represented as a direct connection between two subtelomeric loci. For events supported by one subtelomeric anchor and one pure telomeric mate, the subtelomeric breakpoint was connected to a central telomeric hub, reflecting fusions in which one end could not be resolved to a specific chromosome terminus. All unanchored events were excluded from downstream breakpoint-location comparisons between ALT- and ALT+ CKS tumors.

### Extrachromosomal DNA analysis

Extrachromosomal DNA (ecDNA) was identified from Nanopore sequencing data (*n* = 8) using *CoRAL* ^90^ with default settings. From the resulting BED output files, ecDNA regions were extracted for each tumor sample, and the CN of every unique ecDNA reference segment was calculated. The cumulative ecDNA CN per sample was then computed by summing the CN across all ecDNA-associated genomic segments. To assess ecDNA prevalence across CKS subtypes in the broader cohort, ecDNA was additionally identified from short-read WGS data (*n* = 137) using *AmpliconArchitect* ^135^(v1.5.r7) with default parameters. For eleven cases, amplicon reconstruction could not be resolved due to suboptimal seed regions and CN segmentation, excessively long seed intervals, and/or insufficient breakpoint support. Samples harboring at least one reconstructed ecDNA amplicon were classified as ecDNA+. To determine whether ecDNA structures preferentially arise from specific chromosomes and to characterize the gene content of these amplicons, the chromosomal distribution of protein-coding genes carried on ecDNA was mapped and stratified by ALT status and CKS subtype (**Supplementary Table 1**).

### Logistic regression analysis of genomic predictors of ALT positivity

To identify genomic features strongly associated with ALT positivity across the CKS cohort, logistic regression models were fitted with ALT status as the binary outcome and outward TFs, HRD positivity (defined by *scarHRD*), deleterious *ATRX*/*DAXX* alterations, WGD, high-confidence chromothripsis, and ecDNA status as predictor variables. Each feature was assessed individually, followed by subtype-adjusted models including CKS entity as a covariate to account for differences in subtype composition. Odds ratios and corresponding 95% Wald confidence intervals were calculated from model coefficients, and *P* values were adjusted for multiple testing across predictor terms within each model using the Benjamini-Hochberg method.

### Extraction of high-molecular-weight genomic DNA from tissue

Fresh-frozen tumor tissue samples (20-50 mg per sample) were obtained from the NCT/DKFZ Heidelberg Sample Processing Laboratory. High-molecular-weight (HMW) gDNA was extracted using the Puregene Tissue Kit (Qiagen), following the manufacturer’s protocol with modifications for long-read Nanopore sequencing. Briefly, tissue samples were lysed and processed according to the kit instructions. To ensure complete dissolution of DNA, lysates were incubated at 65°C for no more than 1 hour. Subsequently, samples were incubated overnight at room temperature with gentle agitation (200 rpm). Following extraction, genomic DNA was aliquoted and stored at - 20°C until use. Throughout the extraction and handling process, wide-bore pipette tips were used to minimize mechanical shearing and preserve DNA integrity. Prior to library preparation, the quality and quantity of isolated HMW gDNA were assessed using a Qubit Fluorometer (Invitrogen), and fragment size distribution was evaluated using the 4200 or 2200 TapeStation System (Agilent) and the 2100 Bioanalyzer (Agilent).

### Long-read library preparation

HMW gDNA was sheared for long-read sequencing (10-30 Kb) and the resulting fragments were analyzed with the Femto Pulse system (Agilent). The concentration of sheared DNA was measured with the Qubit dsDNA High Sensitivity Assay Kit (Invitrogen). Libraries were prepared using the Oxford Nanopore Ligation Sequencing Kit V14 (SQK-LSK114) according to the manufacturer’s protocol. The resulting libraries were sequenced on PromethION R10.4.1 flow cells (FLO-PRO114M) using one tumor sample per flow cell. Sequencing was performed at EMBL GeneCore on the P2 Solo device and at the DKFZ Sequencing Open Lab on the PromethION (P24) device. Flow cell priming and loading were carried out according to the instructions from Oxford Nanopore Technologies (ONT). We applied washing and reloading of flow cells to increase data output at a maximum sequencing time of 96 hours.

### Long-read basecalling and alignment

*Dorado* (v0.6.0; https://github.com/nanoporetech/dorado) was used to basecall and align the sequencing data to the human reference genome (GRCh38/hg38) with minimap2^136^. We used the most accurate PromethION basecalling model (sup) with modified base detection of 5-methylcytosine (5mC) and 5-hydroxymethylcytosine (5hmC) in CG contexts (sup, 5mCG_5hmCG model). *NanoPack*^137^ and *Alfred*^138^ were used for alignment quality control and estimating alignment-based sequencing parameters. The average whole-genome coverage was 24.74X with a range of 14.70X to 38.61X for all tumor samples. The average N50 read length was 15,625 bp with a range of 13,283 bp to 21,683 bp for all tumor samples. Using the aligned sequencing data, we estimated a sequencing error rate of 1.47% with a range of 1.10% to 1.90% for all cases. Using verifyBamID^139^, we did not detect any sample cross-contamination.

### Germline variant discovery and phasing

SNVs and indels were called using *Clair3*^140^, and long-read-based phasing of heterozygous markers was performed using *WhatsHap*^141^. We used the default options of *Clair3* and generated haplo-tagged BAM files for allele-specific methylation analyses with *modkit* (v0.6.2; https://github.com/nanoporetech/modkit).

### Somatic variant and copy number variant calling

We used *ClairS* (v0.4.4; https://github.com/HKU-BAL/ClairS) to call somatic SNVs and indels. Somatic SVs were identified using *Delly* ^142^ and *Severus* ^143^ with a panel-of-normal (PoN) from Schloissnig et al.^144^ to distinguish germline and somatic SVs. We used *sansa* (v0.2.5; https://github.com/dellytools/sansa) to merge the SV VCF files from *Delly* and *Severus* and extracted a consensus somatic SV set from both tools to maximize specificity. Somatic CN variants and tumor read-depth profiles were computed using *CNVkit*^145^ and *Delly*^142^ (*cnv* subcommand). We used the ‘*wgs*’ method in *CNVkit* with a 5 kb resolution and applied the default GRCh38/hg38 mappability map with a 25 kb window size for *Delly*.

### Telomeric repeat identification

We run *Lorax* ^81^(PMID: 37082141) with the GRCh38/hg38 reference sequence to search for canonical and non-canonical telomeric repeats, namely TTAGGG, TCAGGG, TGAGGG, TTGGGG, ATAGGG, CTAGGG, GTAGGG, TAAGGG, TTCGGG, and TTTGGG. Lorax was run with default parameters, except for *m* = 25 and *s* = 251, to maximize sensitivity of telomeric repeat identification. We filtered the read-level telomeric breakpoints found by *Lorax* for a mapping quality above 20 and for congruent orientation between soft clips and telomeric repeats (e.g., a match between “clipped_left” with “left_telomeric” and “clipped_right” with “right_telomeric”, respectively). Next, we excluded breakpoints at less than 1 million bp away from centromeres or telomeres to maximize specificity for neo-telomeres and intra-telomeric repeats at SV breakpoints. We obtained information about the repeat classification (intra-telomeric or neo-telomeric) from the *Lorax* output file and clustered the reads for each sample using a region of +/-1000 bp around each SV breakpoint with *BEDtools*^146^. To enhance specificity and exclude potential ONT chimeras, we selected only clusters with at least 2 supporting reads for each telomeric repeat class. After merging regions for each sample, we ensured that no region appeared in multiple samples, as these overlaps likely represent germline variations in variable number tandem repeats (VNTRs) and therefore should be excluded. Because of variable read length and sampling location, intra- and neo-telomeric repeat classification may vary between reads of the same telomeric repeat cluster. We therefore operationally defined a cluster to be intra-telomeric if at least one read spanned the entire telomeric repeat. Likewise, a cluster was classified as neo-telomeric if all supporting reads were assigned as neo-telomeric by *Lorax*. Since most intra-telomeric repeats were shorter than 3000 bp, we further re-classified neo-telomeric repeat clusters with a length <3000 bp as potentially neo-telomeric, as we cannot exclude for these clusters that the coverage and read length were insufficient to span the telomeric repeat.

### Consensus computation of reads spanning intra- and neo-telomeric repeats

For each breakpoint region previously classified as a somatic telomeric repeat, we collected all the reads from the corresponding sample spanning the +/- 1,000bp region and ran *Lorax* on them with the same parameters as before. From all telomeric repeat-containing reads at a given locus, we computed a consensus using *Alfred* with *d* = 0.2. After the consensus computation, we aligned all telomeric reads identified by *Lorax* for that region to the consensus to later call modified bases across the entire telomeric repeat region. To estimate the length and class (intra- or neo-telomeric) of each consensus sequence, we used *Lorax* and re-aligned the consensus sequence to the reference GRCh38/hg38 for SV breakpoint identification. For each consensus sequence, a match plot (including alignment matches of one or more telomeric repeat-containing long reads against GRCh38/hg38) and a telomeric signal plot (used to visualize the intensity of the repeat signal across the length of the consensus sequence) were created using *wally*^81^. A masked GRCh38/hg38 reference sequence with no beginning and end of chromosomes was used for the match plot to avoid the alignment of telomeric repeats to the actual telomeres. For all consensus sequences with a single intra-telomeric repeat, the prefix of the consensus sequence was identified as the region from the beginning of the consensus sequence to the beginning of the repeat. Analogously, the suffix was the region of the consensus after the repeat to the end of the consensus. Prefix and suffix FASTA files were then aligned on the reference with *minimap2* (*map-ont* option) and the coordinates of the alignment were obtained with *BEDtools bamtobed* for each consensus sequence. For each sample, the mapping locations of the prefix and suffix of each consensus were linked via a unique identifying index and plotted onto the read-depth profile of the tumor sample. For complex rearrangements with multiple SV breakpoints in close vicinity, the prefix and suffix may have multiple mapping locations. The prefix and suffix are interchangeable and indicate only where the adjacent genomic segments of a somatic telomeric repeat are anchored in the reference genome. For neo-telomeric repeats, only the prefix or the suffix is present.

### Motif enrichment analysis

We used 1000 bp from the prefix and suffix of each consensus sequence adjacent to the repeat for motif analysis using *XSTREME* ^147^(https://doi.org/10.1101/2021.09.02.458722), with default parameters. We applied the motif search separately for intra-telomeric (those with prefix and suffix) and neo-telomeric consensus sequences (those with only prefix or suffix). All identified motifs and statistical metrics are provided in **Supplementary Table 1**.

### DNA methylation analysis

We applied *modkit* to the alignment of telomeric repeat-containing reads for each consensus sequence and computed methylation frequencies across the entire consensus. For each possible methylation position (CpG context), the mean methylation call probability was calculated from the read-level data. The positions were then ranked with increasing and decreasing numbers for the suffix and prefix of intra-telomeric repeats, respectively, and with increasing numbers for neo-telomeric repeats. A rank of 0 indicates the SV breakpoint with the telomeric repeat and increasing/decreasing ranks to the right and left are progressively more distal CpG sites neighboring the telomeric repeat. Only consensus sequences with more than 150 methylation positions were considered, and the interval between -1000 and +1000 positions was visualized.

### Statistical analysis and plotting

Statistical analyses were performed in R (v4.4.1) using *ggplot2* (v3.5.1; https://cran.r-project.org/web/packages/ggplot2/index.html), *rstatix*(v0.7.2; https://github.com/kassambara/rstatix), limma^57^ (v3.60.6), edgeR^131^ (v4.2.2), *ggpubr* (v0.6.0; https://rpkgs.datanovia.com/ggpubr/) and *ggsignif* (v0.6.4; https://const-ae.github.io/ggsignif/). Comparisons between two groups were primarily conducted using two-sided Wilcoxon rank-sum tests, whereas categorical variables and event frequencies were compared using two-sided Fisher’s exact tests. Wilcoxon tests and *P* value adjustments were implemented using *rstatix*. Correlation analyses were performed using Pearson’s correlation coefficients. Logistic regression models with a binomial family were fitted to assess genomic predictors of ALT positivity. Odds ratios and 95% confidence intervals were derived from Wald-based estimates and the resulting *P* values were adjusted using the Benjamini-Hochberg method. Unless otherwise specified, multiple testing correction was performed using the Benjamini-Hochberg method, with Bonferroni correction applied for selected multiple pairwise subgroup comparisons. Adjusted *P* values < 0.05 were considered statistically significant. The majority of the plots were generated using *ggplot2*, with statistical annotations added using *ggpubr* and *ggsignif*. Significance levels are denoted as follows: *ns*, not significant; * *P* < 0.05, ** *P* < 0.01, *** *P* < 0.001, and **** *P* < 0.0001.

### Use of Large Language Model Tools

During the preparation of this work, the authors used ChatGPT (chatgpt.com) to improve readability and style. After using this tool, the authors reviewed and edited the content as needed and take full responsibility for the content of the final publication.

## Supporting information

Supplementary Figures_Legends_Table 1 description

## Data Availability

The WGS/WES, RNA-seq and long-read sequencing data used in this study are available via controlled access in the German Human Genome-Phenome Archive (GHGA, <u>data.ghga.de</u>) under the GHGA Accession [PLACEHOLDER, to be updated]. Further details, including the data access policy for the study, can be found there.

## Authors Contributions

N.B., T.R., L.F., and P.C. conceptualized the study. N.B., M.L.R., T.R., L.Si., K.R., I.C.-C., L.F., and P.C. designed the experimental and analytical methodology. T.R., M.L.R., L.Si., I.C.-C., and L.F. developed the computational tools and pipelines. K.P., C.G., coordinated sample collection and processing. N.B., A.H., C.K., M.G.W., and S.Sta. performed C-circle assay with input and infrastructure support from C.Sc., F.W., K.R.,V.B., and P.C. N.B., R.P., M.-V.T., S.K., P.Hor., D.H. supported short-read molecular profiling (WGS, WES, RNAseq) and clinical data collection and curation. N.B., T.R., S.St., L.V., V.B. and P.C. generated and curated long-read sequencing data. N.B., M.L.R., R.P., T.R., L.Si., M.G.W. performed formal data analysis with validation contributions from C.E., and U.T. B.Ka., P.Hoh., K.S.-O., U.K., D.R.L., A.L., N.P., T.K., C.H.B., M.B., P.M., F.K., S.B., and C.He. recruited patients and contributed patient samples to the MASTER program, which is supervised by H.G. and S.F. N.B., M.L.R, T.R., L.F., and P.C. performed data interpretation and visualization. N.B. and P.C. wrote the manuscript, with contributions from M.L.R, T.R. I.C.-C and L.F. and critical review from all authors. J.O.K., B.B., L.F., and P.C. supervised the work. P.C. acquired funding and coordinated overall project with additional funding support from J.O.K, M.B.,P.M.

## Acknowledgements

This work was supported by Emmy Noether Individual Programme Grant of the German Research Foundation (DFG; CH 2302/1-1), Sarcoma Foundation of America Grant No. 2022 SFA 01-22, and National Leiomyosarcoma Foundation Grant Annual RFP Call 2023 to P.C. The MASTER program is supported by the NCT Overarching Clinical Translational Trial Program, the NCT Heidelberg Molecular Precision Oncology Program, and DKTK. Additional funding support was received from the German Federal Ministry of Research, Technology and Space (BMBFTR) within the German Cancer Consortium and for PM4Onco – FKZ01ZZ2322A to M.B.,P.M. and the European Research Council (ERC Advanced grant (SEE-MAGIC) grant no.101098056 to J.O.K.

