## Supplementary Figures_Legends_Table 1 description for "Mapping the genomic and transcriptomic features associated with telomere maintenance mechanisms in complex karyotype sarcomas"

**Biondi et al.**

**Supplementary Figure legends**

**Supplementary Figure 1. Screening of human bone and soft-tissue sarcoma samples for ALT activity using the C-circle assay.** C-circle assay dot blots of pan-sarcoma samples ($n$ = 776). Following PCR-based rolling circle amplification of C-circles using Φ29 DNA polymerase, membrane blotting, chemiluminescence-based detection, and subtraction of background signal, C-circle levels were calculated by normalizing to 20 ng of U-2OS standard genomic DNA. Cases with C-circle levels ≥ 0.1 arbitrary units were considered ALT+. The top and bottom rows of the C-circle assay dot blots indicate test (with Φ29 DNA polymerase) and control (without Φ29 DNA polymerase) samples, respectively.

**Supplementary Figure 2. Distribution and predicted functional impact of *ATRX*, *DAXX*, and *TERT* somatic alterations across ALT- and ALT+ complex karyotype sarcomas.** (**a**) Arm-level classification of broad and focal CNAs in CKS tumors based on WGS data ($n$ = 146). CN segments ranging from 0.02 Mb to 10 Mb were classified as focal, while those larger than 10 Mb were designated as broad. Segments are sorted by size, with the red line representing the smoothed density estimate. The zoom-in panel highlights the 0.02 Mb cutoff, below which fragments were discarded. (**b**) Classification of centromere-overlapping broad and focal CN segments into the p- or q-arm, with representative examples mapped to chromosomes 12 (upper panel) and 17 (lower panel). Segments were assigned to the p-arm (green) or q-arm (purple) if 50% or more of their length overlapped with the respective arm. Segments spanning more than 50% of the centromere (red/gray) were classified as centromeric and excluded from downstream analyses. (**c**) Comparison of *ATRX* and *DAXX* TPM levels across CKS samples ($n$ = 134) stratified by alteration category (two-sided Wilcoxon rank-sum tests with Benjamini-Hochberg correction; *** $p < 0.001$, **** $p < 0.0001$). The horizontal line within each boxplot represents the median TPM value. (**d**) Absolute counts of somatic mutations, broad and focal CNAs, and SVs in *ATRX*, *DAXX*, and *TERT* between ALT- and ALT+ CKS samples ($n$ = 146, WGS only), further categorized by WGD status (upper annotation bar). Dot size and color represent the total alteration count for each subtype, stratified by ALT status (right annotation bar). Inactivating (-) and activating (+) labels were assigned to each alteration to denote predicted functional impact using gene-specific rules that integrated transcript levels when possible. All deleterious somatic mutations (loss-of-function variants, variants with CADD PHRED ≥ 30, missense variants with REVEL ≥ 0.5, homozygous deletions, and/or other loss events supported by reduced expression) were categorized as inactivating. For somatic broad and focal CNAs in *ATRX* and *DAXX*, homozygous deletions and other deletion events associated with reduced expression (≤5$^{th}$ percentile relative to the gene-specific wild-type expression distribution) were considered inactivating. *ATRX* and *DAXX* duplications and/or amplifications resulting in increased expression (≥50$^{th}$ percentile of the corresponding wild-type distribution) were considered activating. For *TERT* CNAs, because *TERT* expression was not consistently measurable across all CKS samples, a permissive cutoff for detectable expression was set to $\log_2$(TPM+1) ≥ 0.05. Copy number (CN) gains with detectable expression and/or at least one *TERT* promoter mutation or enhancer hijacking event were classified as activating, whereas CN losses annotated as deleterious (Figure 1b) were classified as inactivating. Alterations without an impact label either fell outside the expression thresholds used for classification or lacked RNA-seq data. No protein expression data were available to independently validate these events. (**e**) Count of matched control and tumor samples across CKS patients classified by *TERT* promoter mutation status, mutation position, and ALT status. (**f**) Representative example of an enhancer hijacking event in DDLS60, involving a cis-type translocation relocating a distal enhancer near the *TERT* transcription start site (TSS). The blue bars mark enhancer regions, while yellow arrows indicate genes, with arrow direction representing the transcription orientation. The *TERT* locus is highlighted in red, with distances between the TSS, SV breakpoint, and enhancer indicated. The sample size (*n*) for each group is indicated. CNA: copy number alteration, SV: structural variant, WGD: whole-genome doubling, LOH: loss of heterozygosity, Mb: megabase, TPM: transcripts per million, U.d.: undetermined, E.H.: enhancer hijacking, DDLS: dedifferentiated liposarcoma, LMS: leiomyosarcoma, ULMS: uterine leiomyosarcoma, OS: osteosarcoma, USTS: undifferentiated soft-tissue sarcoma.

**Supplementary Figure 3. WES-based genomic analyses, WGS-based characterization of SVs and chromosome-resolved map of chromothripsis in ALT- and ALT+ complex karyotype sarcomas.** (**a**) Analysis of ploidy levels between ALT- and ALT+ CKS samples using WES data (*n* = 176). (**b-d**) Load of somatic broad CNAs (b), focal CNAs (c), and mutations (d) in protein-coding genes in ALT- and ALT+ CKS tumors based on WES data. The width of the violin plots reflects the density of the data distribution. (**e**) Absolute counts of SVs per CKS sample (*n* = 145, WGS only) stratified by ALT status, sarcoma subtype, and SV type (Del: deletion, Dup: duplication, Ins: insertion, H2H Inv: head-to-head inversion, T2T Inv: tail-to-tail inversion, Tra: translocation). All pairwise comparisons were conducted using two-sided Wilcoxon rank-sum tests with the Benjamini-Hochberg correction (ns: not significant, * $p < 0.05$, ** $p < 0.01$, *** $p < 0.001$, **** $p < 0.0001$). The center horizontal line of boxplots indicates the median value. (**f**) Distribution of low-confidence and high-confidence chromothripsis events across individual chromosomes in ALT- (upper) and ALT+ (lower) CKS tumors. The upper annotation bar highlights the percentage of ALT-specific chromothripsis calls per chromosome. The sample size (*n*) for each group is indicated. CNA: copy number alteration, SV: structural variant, DDLS: dedifferentiated liposarcoma, LMS: leiomyosarcoma, ULMS: uterine leiomyosarcoma, OS: osteosarcoma, USTS: undifferentiated soft-tissue sarcoma.

**Supplementary Figure 4. Chromothriptic architechture and homologous recombination deficiency in complex karyotype sarcomas.** (**a**) Representative examples of canonical chromothripsis events involving different chromosomes in selected ALT- and ALT+ DDLS, LMS, ULMS, OS, and USTS samples. Each case is shown as a ReConPlot, formed by a chromosome ideogram (bottom panel, with red cytobands marking centromeres), CN profiles (middle panel), and SV rearrangements (top panel). SVs connecting breakpoints within the displayed region are shown as arcs, whereas SVs with only one breakpoint in the displayed region are shown as vertical lines with overhangs. Black CN segments indicate total copy number, whereas grey CN segments indicate minor copy number. (**b**) *scarHRD* score analysis between ALT- and ALT+ CKS samples based on ploidy status (*n* = 324, combined WES and WGS datasets). A cut-off value of 42 (dotted line) marks the threshold above which samples were considered HRD+. (**c**) Quantification of HRD probability using *HRProfiler* in the CKS cohort ($n$ = 289, combined WES and WGS datasets). CKS subtypes are ranked based on increasing median HRD probability. (**d**) Comparison of HRD probability values between merged ALT- and ALT+ CKS samples (left) and further stratification by CKS subtype (right). (**e**) HRD probability analysis between ALT- and ALT+ CKS samples based on ploidy status. The dotted line represents the HRD cut-off value set at 0.42 above which samples were classified as HRD+. The center horizontal line of boxplots indicates the median value. Pairwise comparisons were conducted using two-sided Wilcoxon rank-sum tests with the Bonferroni correction (ns: not significant, * $p < 0.05$, ** $p < 0.01$, *** $p < 0.001$, **** $p < 0.0001$). The sample size ($n$) for each group is indicated. CN: copy number, SV: structural variant, Mb: megabase, HRD: homologous recombination deficiency, DDLS: dedifferentiated liposarcoma, LMS: leiomyosarcoma, ULMS: uterine leiomyosarcoma, OS: osteosarcoma, USTS: undifferentiated soft-tissue sarcoma.

**Supplementary Figure 5. Subtype-resolved comparative analysis of copy number profiles between ALT- and ALT+ complex karyotype sarcomas.** (**a**) Proportion of CN gains (red) and losses (purple) across chromosomes for ALT- (top panel) and ALT+ (bottom panel) CKS tumors ($n$ = 146, WGS only), stratified by subtype. Profiles from entities with fewer than 10 WGS samples are included for completeness but should be viewed as exploratory. (**b**) Representative examples of GenomeRangerPlots illustrating unique and shared focal duplications involving *TOP3A*, *CENPV*, *TRIM16*, and *NCOR1* at chromosome 17 in merged CKS samples (WGS only). CN segments were ranked based on size and gene specificity. *MAP2K4* was excluded because only a single shared event was detected with *TOP3A*. The sample size ($n$) for each group is indicated. CN: copy number, DDLS: differentiated liposarcoma, LMS: leiomyosarcoma, ULMS: uterine leiomyosarcoma, OS: osteosarcoma, USTS: undifferentiated soft-tissue sarcoma.

**Supplementary Figure 6. Transcriptional programs and pathway activity associated with TMM status in CKS.** (**a-b**) Pie charts summarizing the distribution of tumor samples with RNA-sequencing data ($n$ = 260) based on CKS subtype (a) and ALT status (b). (**c**) GSEA-based comparison of selected biological processes enriched in ALT- and ALT+ CKS tumors. DEGs were ranked by $\log_2$(fold-change) values. The most recurrent DEGs contributing to the gene sets are shown. (**d**) Activity inference of curated PROGENy pathways across CKS subtypes using *decoupleR*. The pathway activity score is color-coded to show whether it is higher or lower in ALT+ tumors. (**e**) Scatterplot illustrating transcriptional modulation of p53 pathway components in ALT+ CKS tumors. Genes are compared by their inferred influence on the pathway (PROGENy weight) and differential expression (t-value). Key genes contributing to pathway activation or repression are highlighted. The sample size ($n$) for each group is indicated. GSEA: gene set enrichment analysis, DEG: differentially expressed gene, NES: normalized enrichment score, DDLS: dedifferentiated liposarcoma, LMS: leiomyosarcoma, ULMS: uterine leiomyosarcoma, OS: osteosarcoma, USTS: undifferentiated soft-tissue sarcoma.

**Supplementary Figure 7. Pan-sarcoma analysis of telomere content and its correlation with telomere fusion rate in complex karyotype sarcomas.** (**a**) Telomere content analysis across the pan-sarcoma cohort ($n$ = 340, combined WES and WGS datasets). A comprehensive list of sarcoma subtype abbreviations is provided in **Supplementary Table 1**. (**b**) Pearson correlation analysis between telomere content and TF rate in ALT- and ALT+ CKS tumors (*n* = 80). Each dot represents a tumor sample. Pearson's correlation coefficient (*r*), 95% confidence interval, and *P* value are shown. The center horizontal line of boxplots indicates the median value. All pairwise comparisons were conducted using two-sided Wilcoxon rank-sum tests with Benjamini-Hochberg correction (ns: not significant, * $p < 0.05$, ** $p < 0.01$, *** $p < 0.001$, **** $p < 0.0001$). The sample size (*n*) for each group is indicated. T/C: tumor/control, TPM: transcripts per million, DDLS: dedifferentiated liposarcoma, LMS: leiomyosarcoma, ULMS: uterine leiomyosarcoma, OS: osteosarcoma, USTS: undifferentiated soft-tissue sarcoma.

**Supplementary Figure 8. Characterization of telomeric repeat clusters and their sequence context in CKS using Nanopore sequencing.** (**a**) Comparison of somatic SV burden per chromosome with or without canonical TRCs, identified by Nanopore sequencing across the eight CKS samples and stratified by ALT status. Pairwise comparisons were performed using two-sided Wilcoxon rank-sum tests with Benjamini-Hochberg correction (ns: not significant, *** $p < 0.001$). (**b**) Genome-wide distribution of TRCs between ALT- and ALT+ CKS samples. Each mark represents one TRC, with color denoting the CKS entity and symbols representing TRC classes (intra-/inter-chromosomal or neo-telomeric). (**c**) Correlation between C-circle levels and the number of TRCs across the 8 CKS patients, colored by ALT status. Pearson's correlation coefficient (*r*), 95% confidence interval, and *P* value are shown. (**d**) Length distribution of TRCs across CKS subtypes stratified by ALT status (left) and TRC class (right). The dashed line marks the cutoff used to classify neo-telomeric events shorter than 3 kb as potentially neo-telomeric. (**e**) Total count of TRCs per sample, stratified by event type and ALT status. (**f**) DNA methylation probabilities at CpG sites across consensus sequences assembled from long reads spanning ±1 Kb around TRC-associated SV breakpoints, shown for intra-chromosomal insertions and neo-telomeres. (**g**) *XSTREME* motif analysis of ±1 Kb regions surrounding intra-/inter-chromosomal insertions and neo-telomere. Upper panels show enriched motifs identified by XSTREME, whereas lower panels show the corresponding consensus motifs derived from TRC-supporting long reads. (**h**) Representative examples of complex telomeric repeat-containing long reads with multiple genomic mapping locations, illustrating inter-chromosomal rearrangements associated with telomere-mediated healing. Read orientation and mapped genomic intervals are indicated. SV: structural variant, TRC: telomeric repeat cluster, Kb: kilobase, DDLS: dedifferentiated liposarcoma, ULMS: uterine leiomyosarcoma, OS: osteosarcoma, USTS: undifferentiated soft-tissue sarcoma.

**Supplementary Figure 9. Telomere repeat cluster profiling in ALT- and ALT+ CKS tumors using Nanopore sequencing.** Circos plots showing the genome-wide distribution of TRCs detected by Nanopore sequencing in the eight CKS samples, arranged by subtype and ALT status. From outermost to innermost, tracks indicate cytobands, CN profile, SV breakpoint count, and *CoRAL*-predicted amplicon count. Internal links mark intra- or inter-chromosomal TRCs, neo-telomeres, and putative neo-telomeric events. TRC: telomeric repeat cluster, CN: copy number profile, DDLS: dedifferentiated liposarcoma, ULMS: uterine leiomyosarcoma, OS: osteosarcoma, USTS: undifferentiated soft-tissue sarcoma.

**Supplementary Figure 10. Chromosomal distribution of *CoRAL*-resolved linear amplicon burden across CKS subtypes using Nanopore sequencing.** Focal linear amplifications were reconstructed using *CoRAL* across eight ALT- and ALT+ CKS samples. The total number of *CoRAL*-resolved linear amplicons is shown per CKS entity and stratified by ALT status (lower left panel). Additionally, *CoRAL*-predicted linear amplicons were aggregated by chromosome within each subtype to calculate their cumulative amplicon length (right panel), color-coded based on ALT status. The cumulative length of *CoRAL*-resolved linear amplicons from chromosome 12 in ALT- and ALT+ DDLS samples is shown separately (upper left panel). Mb: megabase, Kb: kilobase, DDLS: dedifferentiated liposarcoma, ULMS: uterine leiomyosarcoma, OS: osteosarcoma, USTS: undifferentiated soft-tissue sarcoma.

**Supplementary Figure 11. Genomic architecture and gene content of ecDNA CKS.** (**a**) Analysis of the focal amplicon architecture using *AmpliconArchitect* across ALT- and ALT+ CKS tumors (*n* = 137, WGS only). Detected amplicons were assigned to four decomposition classes, including linear, extrachromosomal DNA (ecDNA), complex non-cyclic, or breakage-fusion-bridge (BFB). Bars show the number of *AmpliconArchitect*-predicted amplicons in each class, stratified by CKS subtype and ALT status. (**b**) Chromosomal distribution of protein-coding genes identified on ecDNA amplicons in ALT- and ALT+ CKS samples. For each chromosome, stacked bars show the total number of ecDNA-associated genes. Representative recurrent or subtype-enriched genes are annotated at their chromosomal positions and colored by entity, while genes shared between ALT- and ALT+ CKS tumors are included in the middle panel, with bold labels indicating canonical oncogenes (**Supplementary Table 1**). TF: telomere fusion, WGD: whole-genome doubling, HRD: homologous recombination deficiency, DDLS: dedifferentiated liposarcoma, ULMS: uterine leiomyosarcoma, OS: osteosarcoma, USTS: undifferentiated soft-tissue sarcoma.

**Supplementary Table 1.** This table contains following information in separate sheets. 1. Sarcoma abbreviations used in this study. 2. Results of the C-circle assay evaluation, 3. A list of *ATRX*, *DAXX* and *TERT* mutations detected in CKS cohort. 4. A list of *BRCA1* & *BRCA2* alterations detected in the CKS cohort. 5. Differentially expressed genes between ALT+ and ALT- CKS as detected by *LimmaVoom*. 6. Results of the Xstreme Motif analysis of the long-read sequencing data.

### Supplementary Figure 1

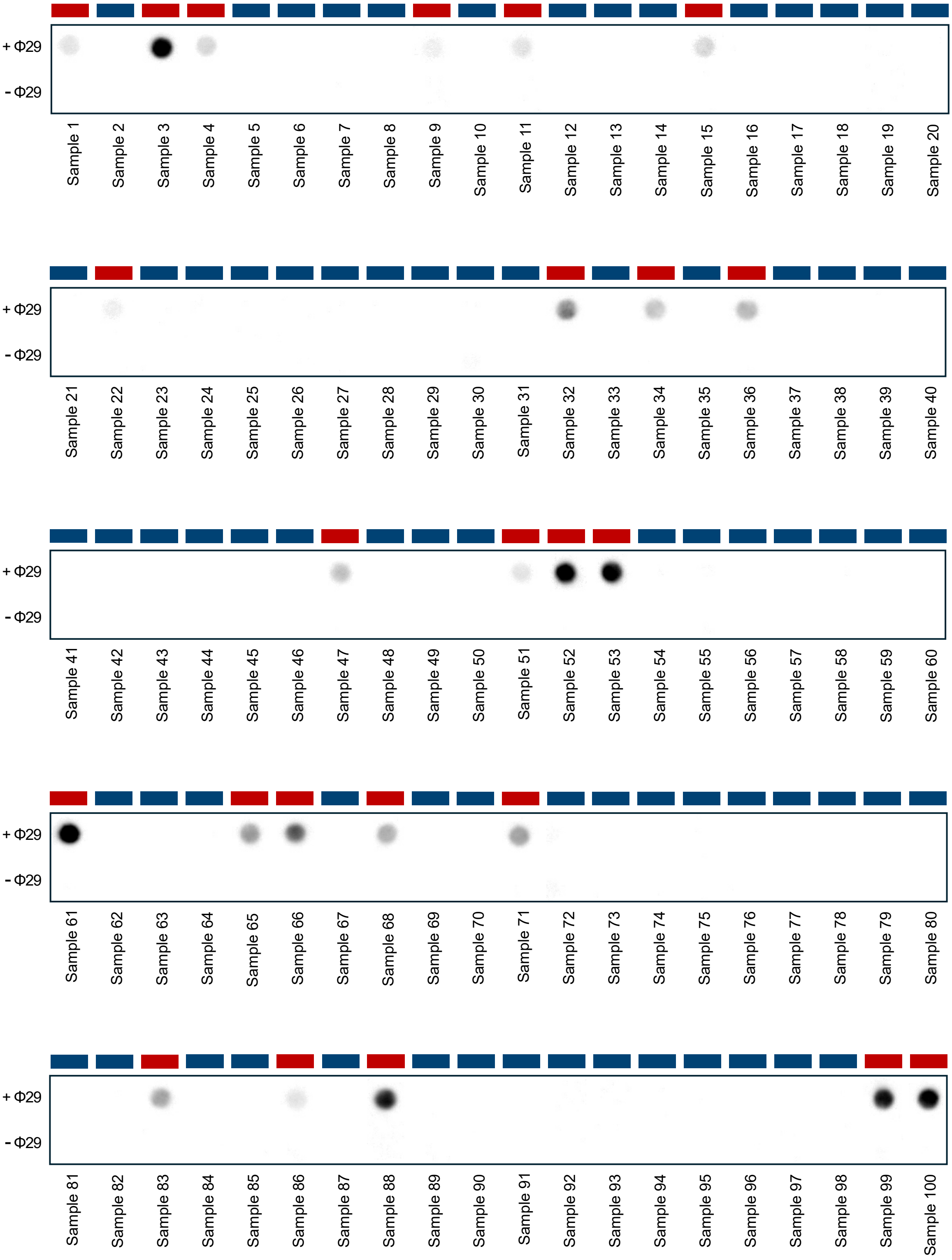


**ALT status:** ALT- ALT+ *(continued)*

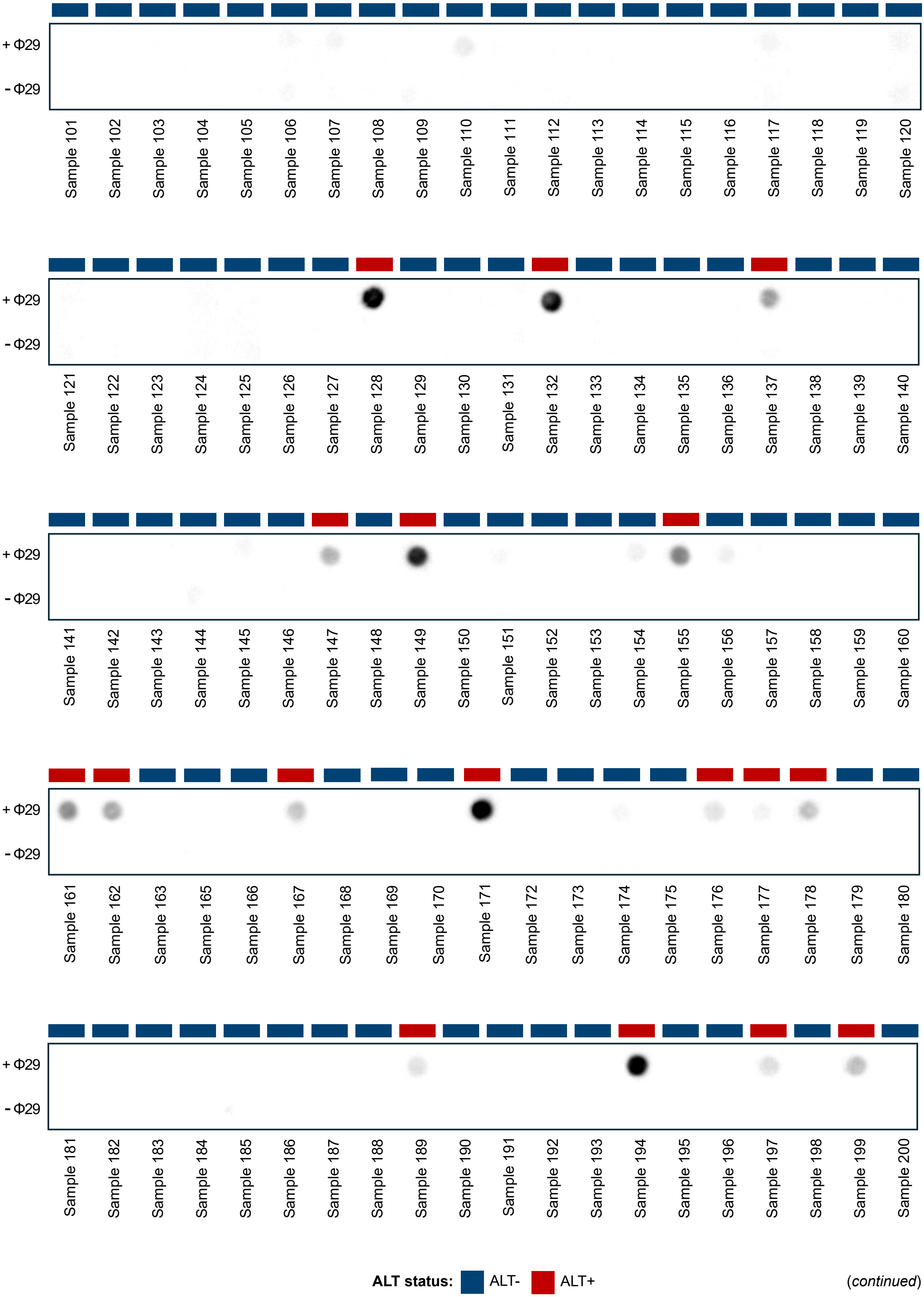

+ Φ29
- Φ29
Sample 101
Sample 102
Sample 103
Sample 104
Sample 105
Sample 106
Sample 107
Sample 108
Sample 109
Sample 110
Sample 111
Sample 112
Sample 113
Sample 114
Sample 115
Sample 116
Sample 117
Sample 118
Sample 119
Sample 120
+ Φ29
- Φ29
Sample 121
Sample 122
Sample 123
Sample 124
Sample 125
Sample 126
Sample 127
Sample 128
Sample 129
Sample 130
Sample 131
Sample 132
Sample 133
Sample 134
Sample 135
Sample 136
Sample 137
Sample 138
Sample 139
Sample 140
+ Φ29
- Φ29
Sample 141
Sample 142
Sample 143
Sample 144
Sample 145
Sample 146
Sample 147
Sample 148
Sample 149
Sample 150
Sample 151
Sample 152
Sample 153
Sample 154
Sample 155
Sample 156
Sample 157
Sample 158
Sample 159
Sample 160
+ Φ29
- Φ29
Sample 161
Sample 162
Sample 163
Sample 165
Sample 166
Sample 167
Sample 168
Sample 169
Sample 170
Sample 171
Sample 172
Sample 173
Sample 174
Sample 175
Sample 176
Sample 177
Sample 178
Sample 179
Sample 180
+ Φ29
- Φ29
Sample 181
Sample 182
Sample 183
Sample 184
Sample 185
Sample 186
Sample 187
Sample 188
Sample 189
Sample 190
Sample 191
Sample 192
Sample 193
Sample 194
Sample 195
Sample 196
Sample 197
Sample 198
Sample 199
Sample 200
ALT status: ALT- ALT+


*(continued)*

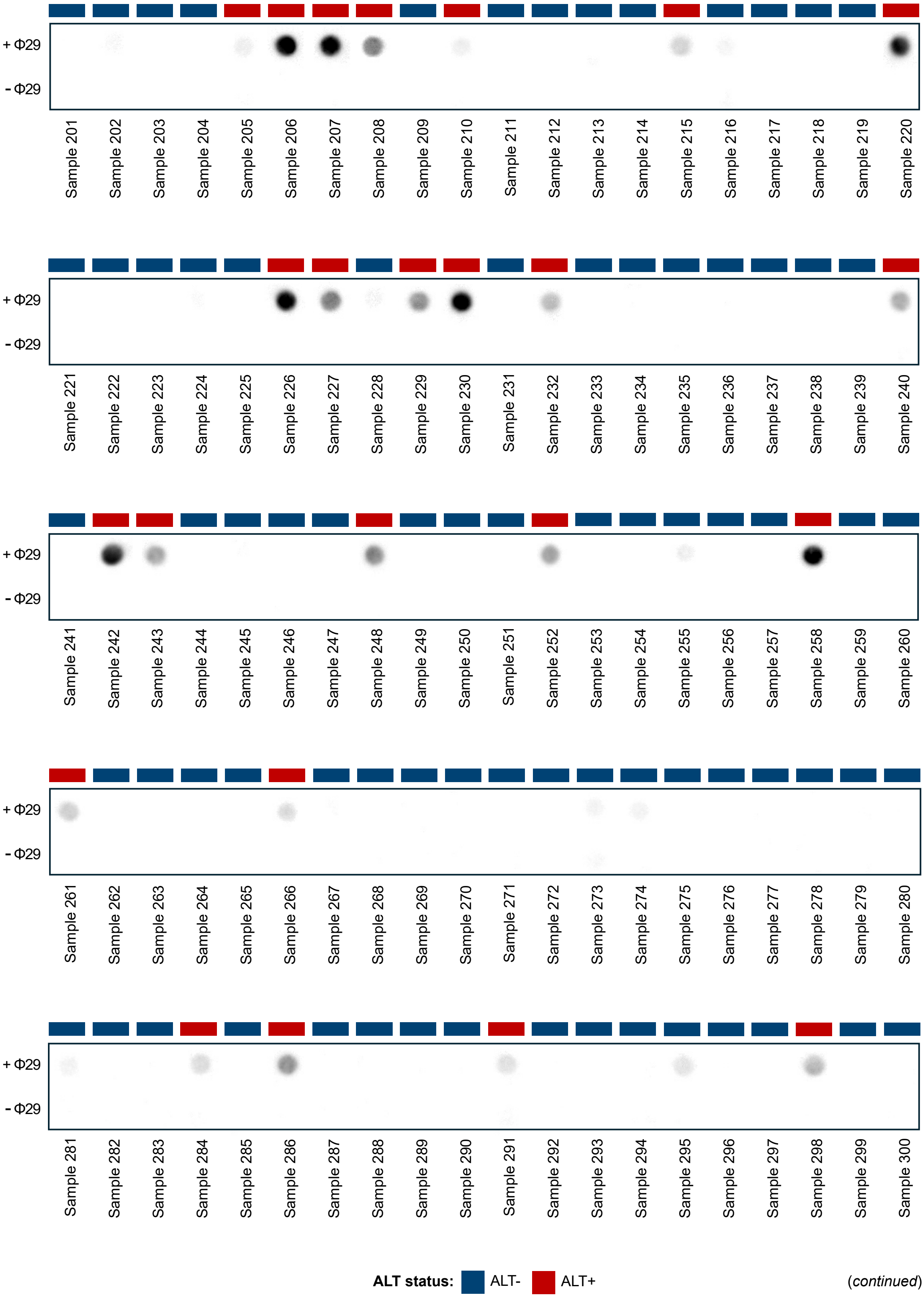
+ Φ29
- Φ29
Sample 201
Sample 202
Sample 203
Sample 204
Sample 205
Sample 206
Sample 207
Sample 208
Sample 209
Sample 210
Sample 211
Sample 212
Sample 213
Sample 214
Sample 215
Sample 216
Sample 217
Sample 218
Sample 219
Sample 220
+ Φ29
- Φ29
Sample 221
Sample 222
Sample 223
Sample 224
Sample 225
Sample 226
Sample 227
Sample 228
Sample 229
Sample 230
Sample 231
Sample 232
Sample 233
Sample 234
Sample 235
Sample 236
Sample 237
Sample 238
Sample 239
Sample 240
+ Φ29
- Φ29
Sample 241
Sample 242
Sample 243
Sample 244
Sample 245
Sample 246
Sample 247
Sample 248
Sample 249
Sample 250
Sample 251
Sample 252
Sample 253
Sample 254
Sample 255
Sample 256
Sample 257
Sample 258
Sample 259
Sample 260
+ Φ29
- Φ29
Sample 261
Sample 262
Sample 263
Sample 264
Sample 265
Sample 266
Sample 267
Sample 268
Sample 269
Sample 270
Sample 271
Sample 272
Sample 273
Sample 274
Sample 275
Sample 276
Sample 277
Sample 278
Sample 279
Sample 280
+ Φ29
- Φ29
Sample 281
Sample 282
Sample 283
Sample 284
Sample 285
Sample 286
Sample 287
Sample 288
Sample 289
Sample 290
Sample 291
Sample 292
Sample 293
Sample 294
Sample 295
Sample 296
Sample 297
Sample 298
Sample 299
Sample 300
ALT status:
ALT-
ALT+

*(continued)*

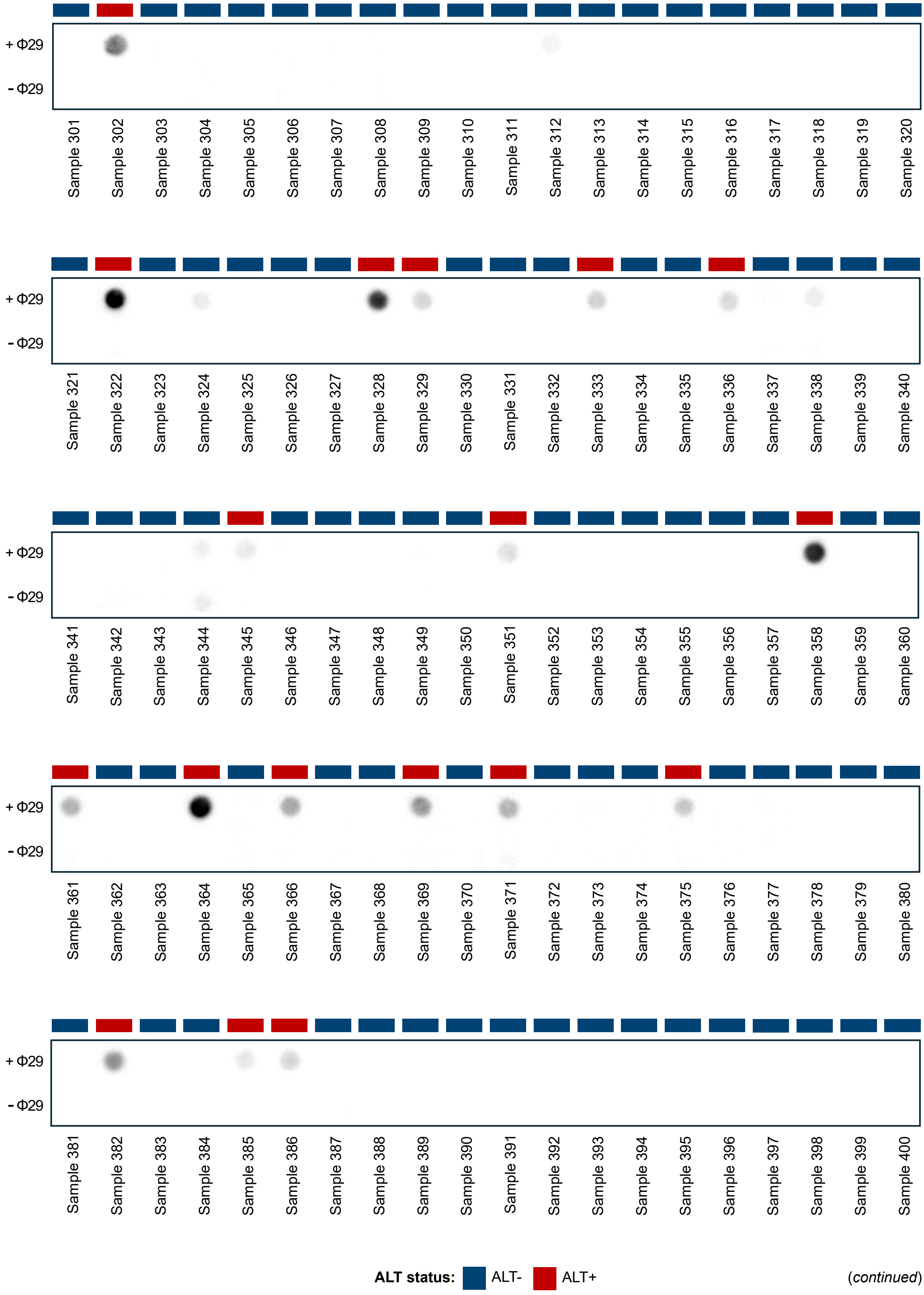
+ Φ29
- Φ29
Sample 301
Sample 302
Sample 303
Sample 304
Sample 305
Sample 306
Sample 307
Sample 308
Sample 309
Sample 310
Sample 311
Sample 312
Sample 313
Sample 314
Sample 315
Sample 316
Sample 317
Sample 318
Sample 319
Sample 320
Sample 321
Sample 322
Sample 323
Sample 324
Sample 325
Sample 326
Sample 327
Sample 328
Sample 329
Sample 330
Sample 331
Sample 332
Sample 333
Sample 334
Sample 335
Sample 336
Sample 337
Sample 338
Sample 339
Sample 340
Sample 341
Sample 342
Sample 343
Sample 344
Sample 345
Sample 346
Sample 347
Sample 348
Sample 349
Sample 350
Sample 351
Sample 352
Sample 353
Sample 354
Sample 355
Sample 356
Sample 357
Sample 358
Sample 359
Sample 360
Sample 361
Sample 362
Sample 363
Sample 364
Sample 365
Sample 366
Sample 367
Sample 368
Sample 369
Sample 370
Sample 371
Sample 372
Sample 373
Sample 374
Sample 375
Sample 376
Sample 377
Sample 378
Sample 379
Sample 380
Sample 381
Sample 382
Sample 383
Sample 384
Sample 385
Sample 386
Sample 387
Sample 388
Sample 389
Sample 390
Sample 391
Sample 392
Sample 393
Sample 394
Sample 395
Sample 396
Sample 397
Sample 398
Sample 399
Sample 400
ALT status: ALT- ALT+

*(continued)*

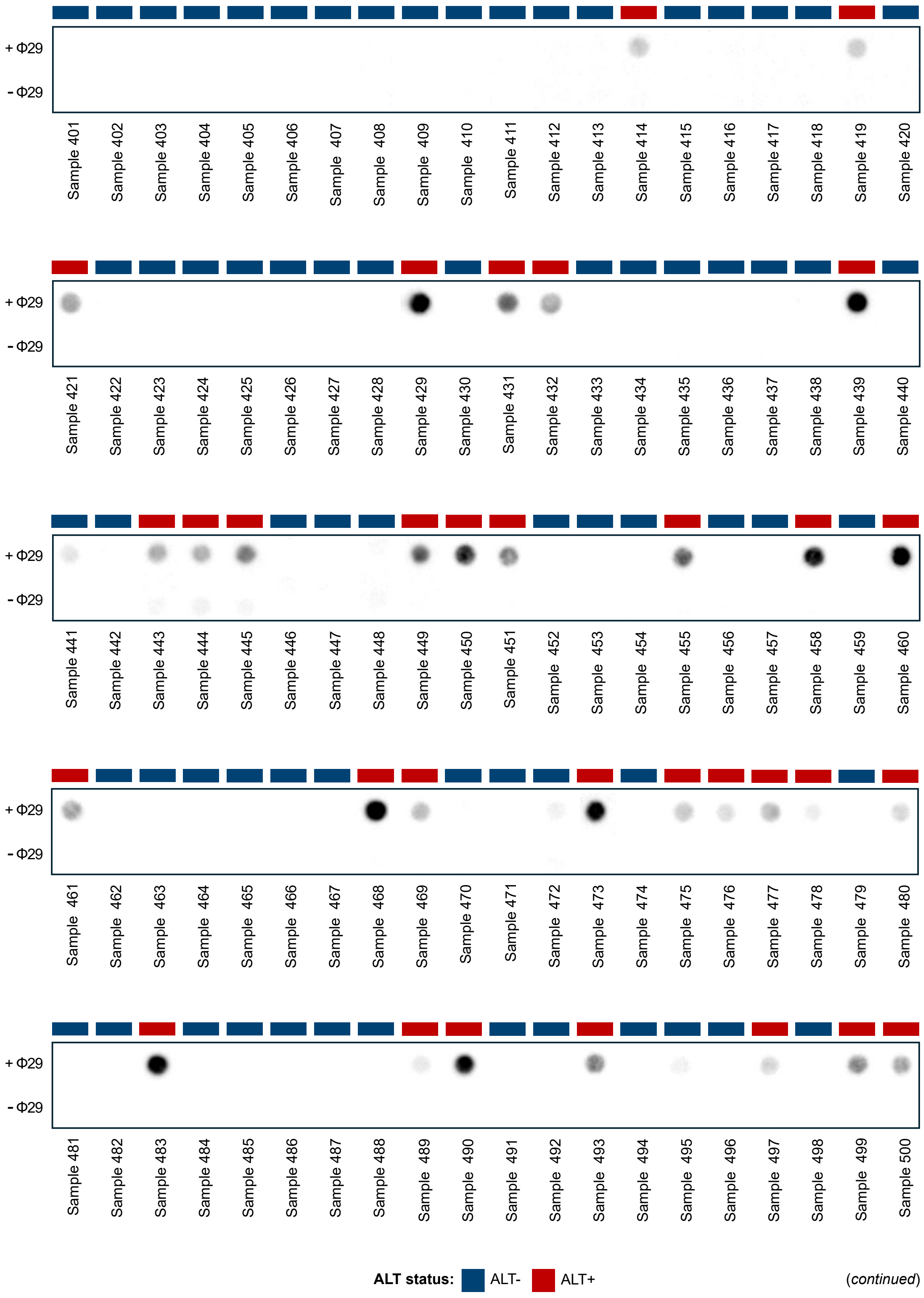
+ Φ29
- Φ29
Sample 401
Sample 402
Sample 403
Sample 404
Sample 405
Sample 406
Sample 407
Sample 408
Sample 409
Sample 410
Sample 411
Sample 412
Sample 413
Sample 414
Sample 415
Sample 416
Sample 417
Sample 418
Sample 419
Sample 420
Sample 421
Sample 422
Sample 423
Sample 424
Sample 425
Sample 426
Sample 427
Sample 428
Sample 429
Sample 430
Sample 431
Sample 432
Sample 433
Sample 434
Sample 435
Sample 436
Sample 437
Sample 438
Sample 439
Sample 440
Sample 441
Sample 442
Sample 443
Sample 444
Sample 445
Sample 446
Sample 447
Sample 448
Sample 449
Sample 450
Sample 451
Sample 452
Sample 453
Sample 454
Sample 455
Sample 456
Sample 457
Sample 458
Sample 459
Sample 460
Sample 461
Sample 462
Sample 463
Sample 464
Sample 465
Sample 466
Sample 467
Sample 468
Sample 469
Sample 470
Sample 471
Sample 472
Sample 473
Sample 474
Sample 475
Sample 476
Sample 477
Sample 478
Sample 479
Sample 480
Sample 481
Sample 482
Sample 483
Sample 484
Sample 485
Sample 486
Sample 487
Sample 488
Sample 489
Sample 490
Sample 491
Sample 492
Sample 493
Sample 494
Sample 495
Sample 496
Sample 497
Sample 498
Sample 499
Sample 500
ALT status: ALT- ALT+

(*continued*)

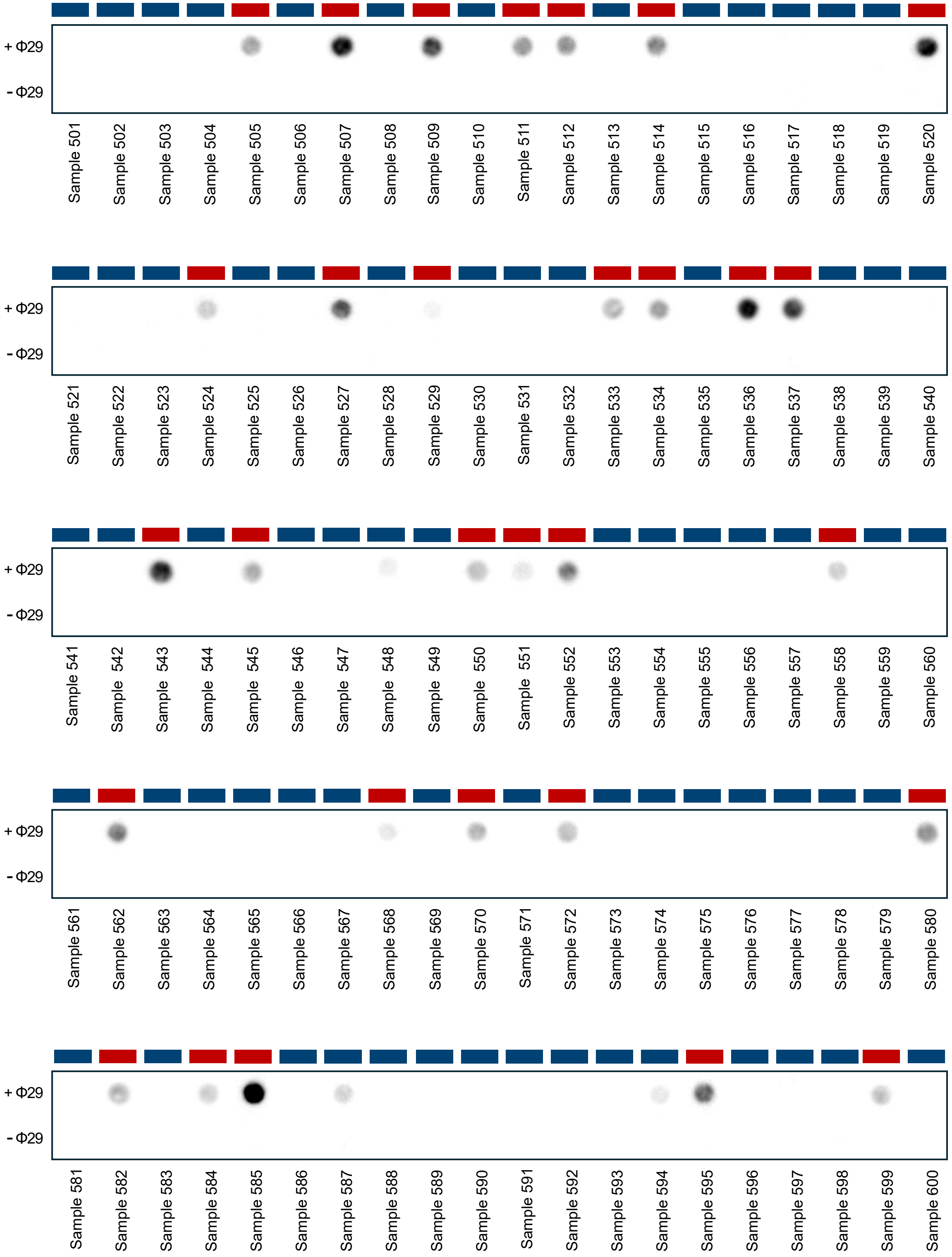


**ALT status:** ALT- ALT+

(*continued*)

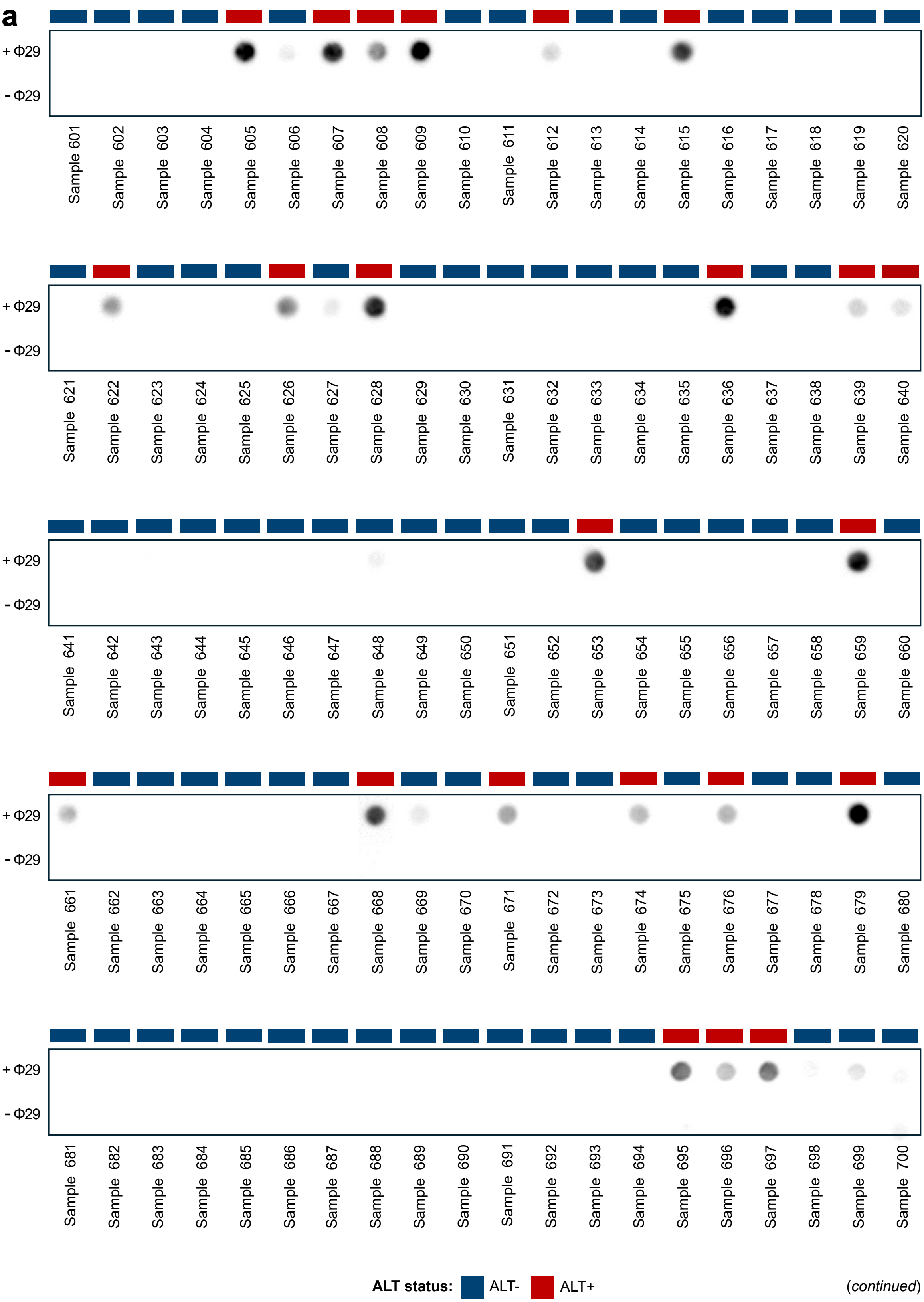


(*continued*)

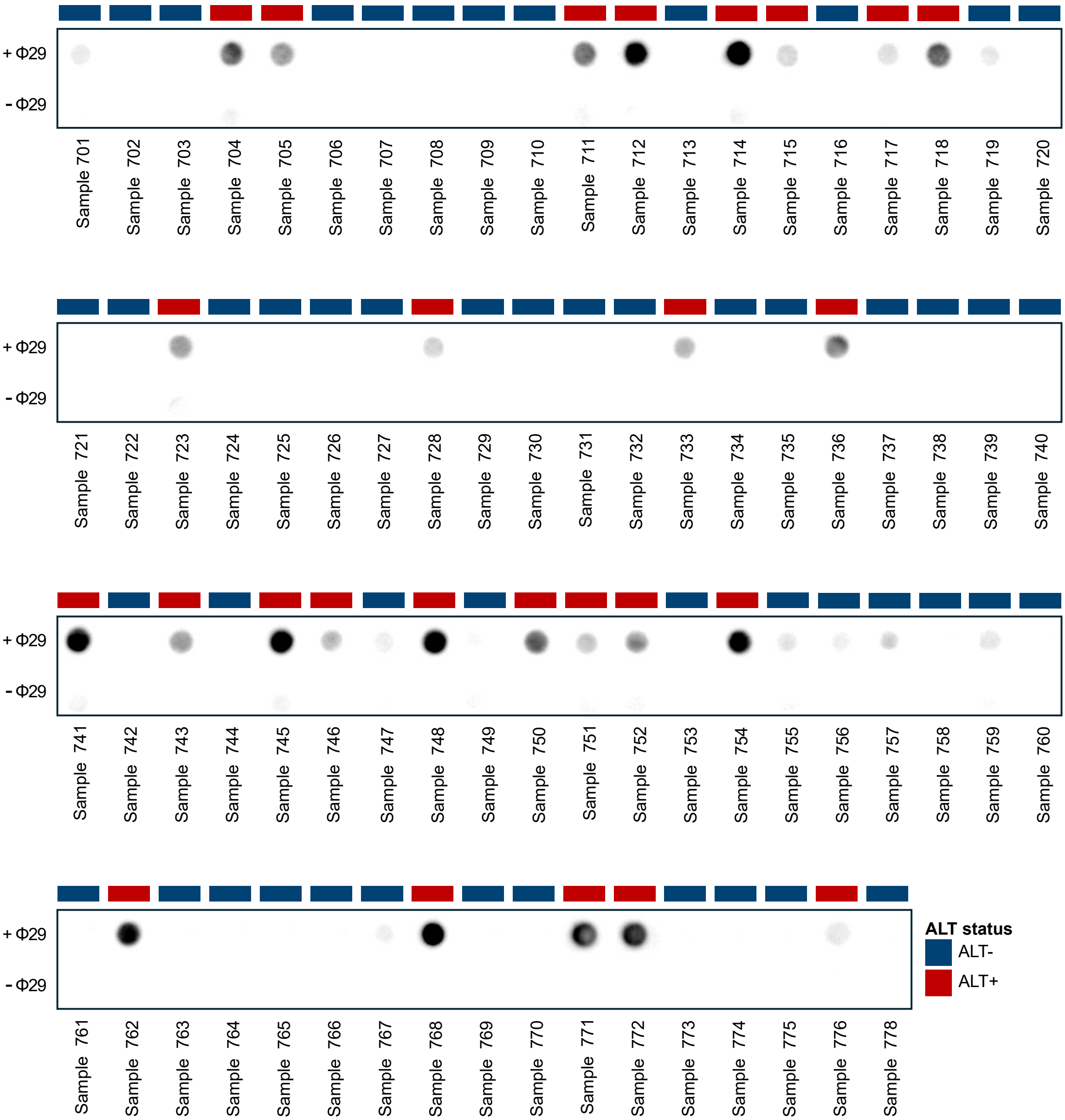
+ Φ29
- Φ29
Sample 701
Sample 702
Sample 703
Sample 704
Sample 705
Sample 706
Sample 707
Sample 708
Sample 709
Sample 710
Sample 711
Sample 712
Sample 713
Sample 714
Sample 715
Sample 716
Sample 717
Sample 718
Sample 719
Sample 720
+ Φ29
- Φ29
Sample 721
Sample 722
Sample 723
Sample 724
Sample 725
Sample 726
Sample 727
Sample 728
Sample 729
Sample 730
Sample 731
Sample 732
Sample 733
Sample 734
Sample 735
Sample 736
Sample 737
Sample 738
Sample 739
Sample 740
+ Φ29
- Φ29
Sample 741
Sample 742
Sample 743
Sample 744
Sample 745
Sample 746
Sample 747
Sample 748
Sample 749
Sample 750
Sample 751
Sample 752
Sample 753
Sample 754
Sample 755
Sample 756
Sample 757
Sample 758
Sample 759
Sample 760
+ Φ29
- Φ29
Sample 761
Sample 762
Sample 763
Sample 764
Sample 765
Sample 766
Sample 767
Sample 768
Sample 769
Sample 770
Sample 771
Sample 772
Sample 773
Sample 774
Sample 775
Sample 776
Sample 778
ALT status
ALT-
ALT+

### Supplementary Figure 2

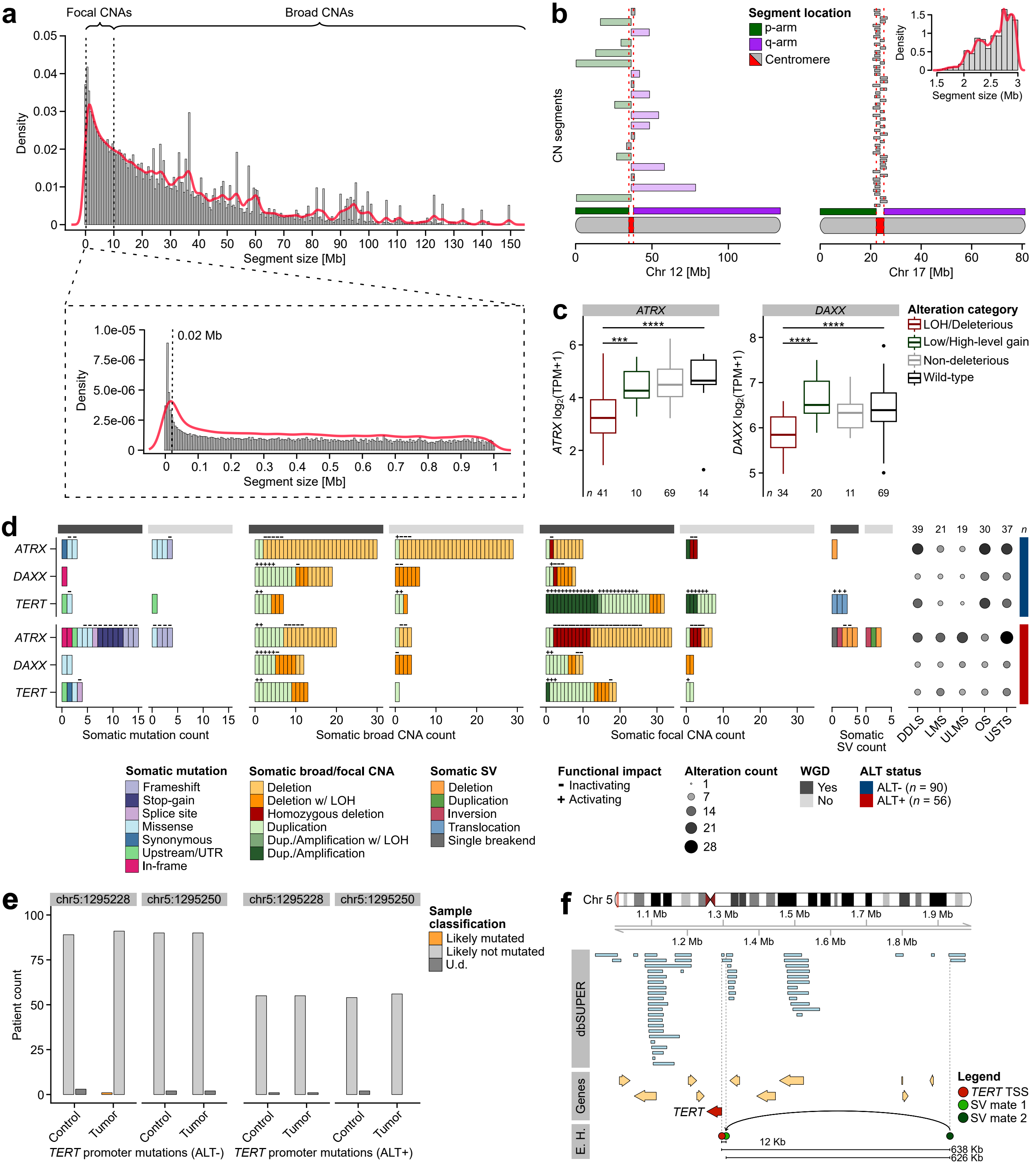

### Supplementary Figure 3

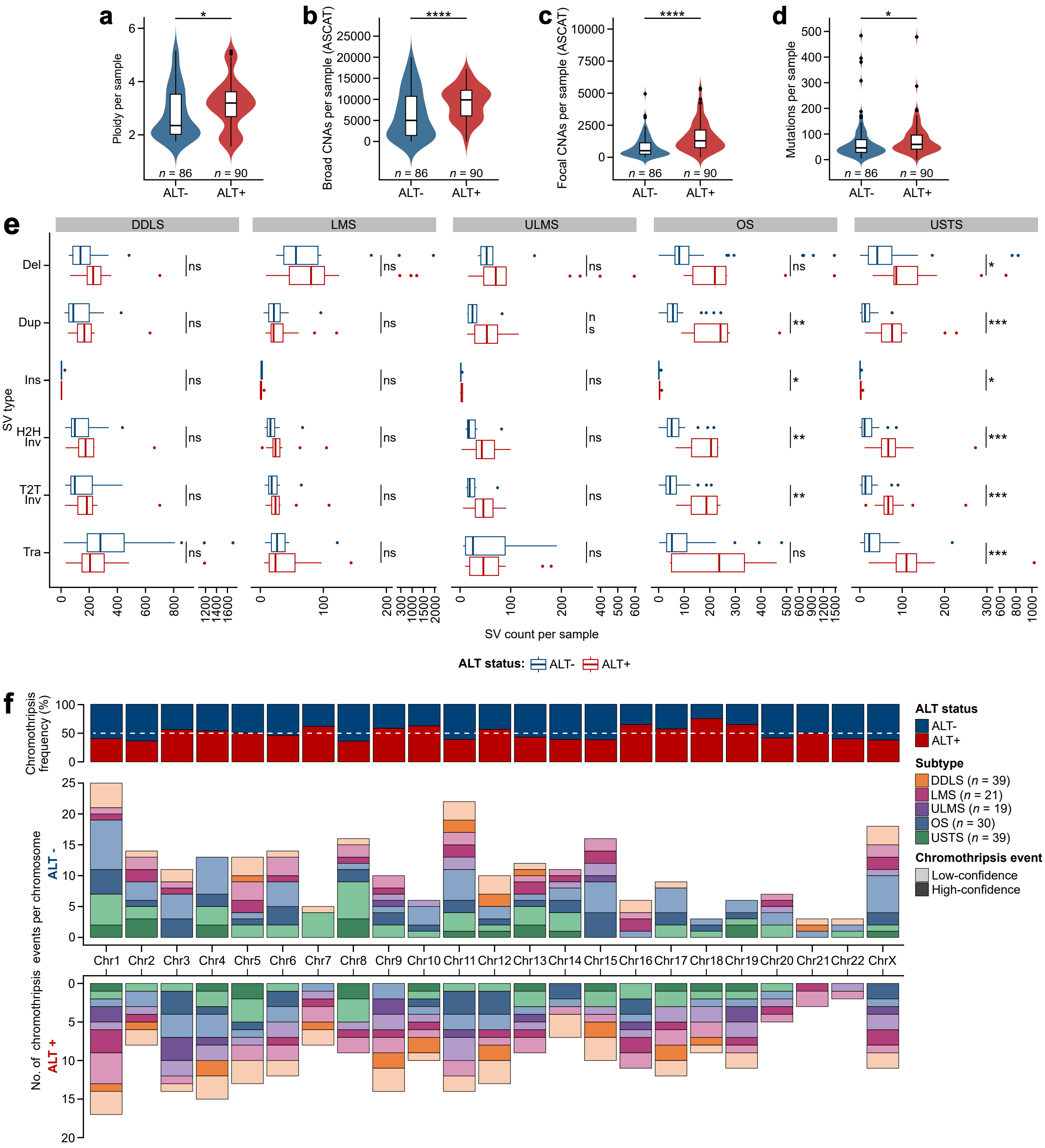

### Supplementary Figure 4

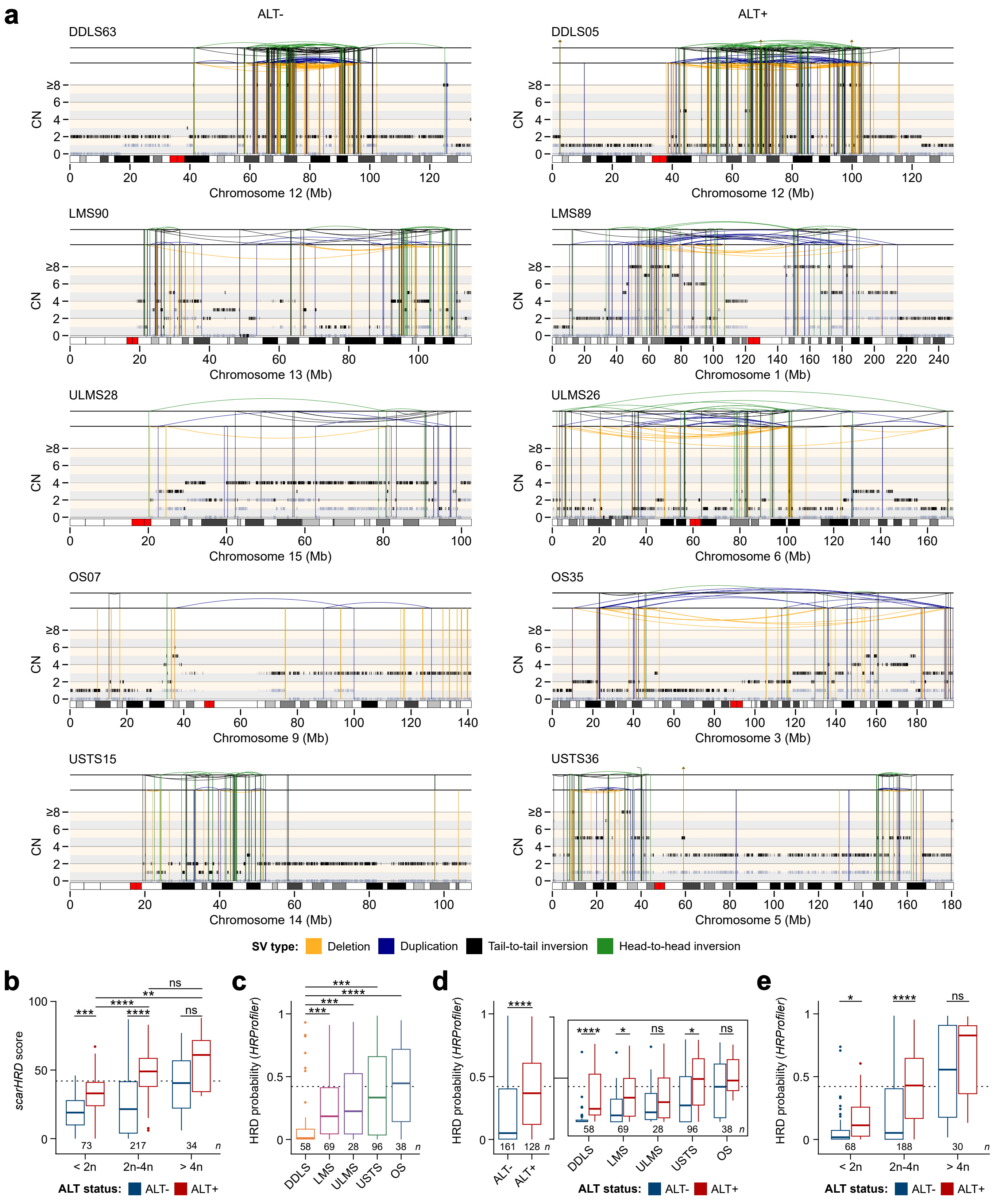

### Supplementary Figure 5

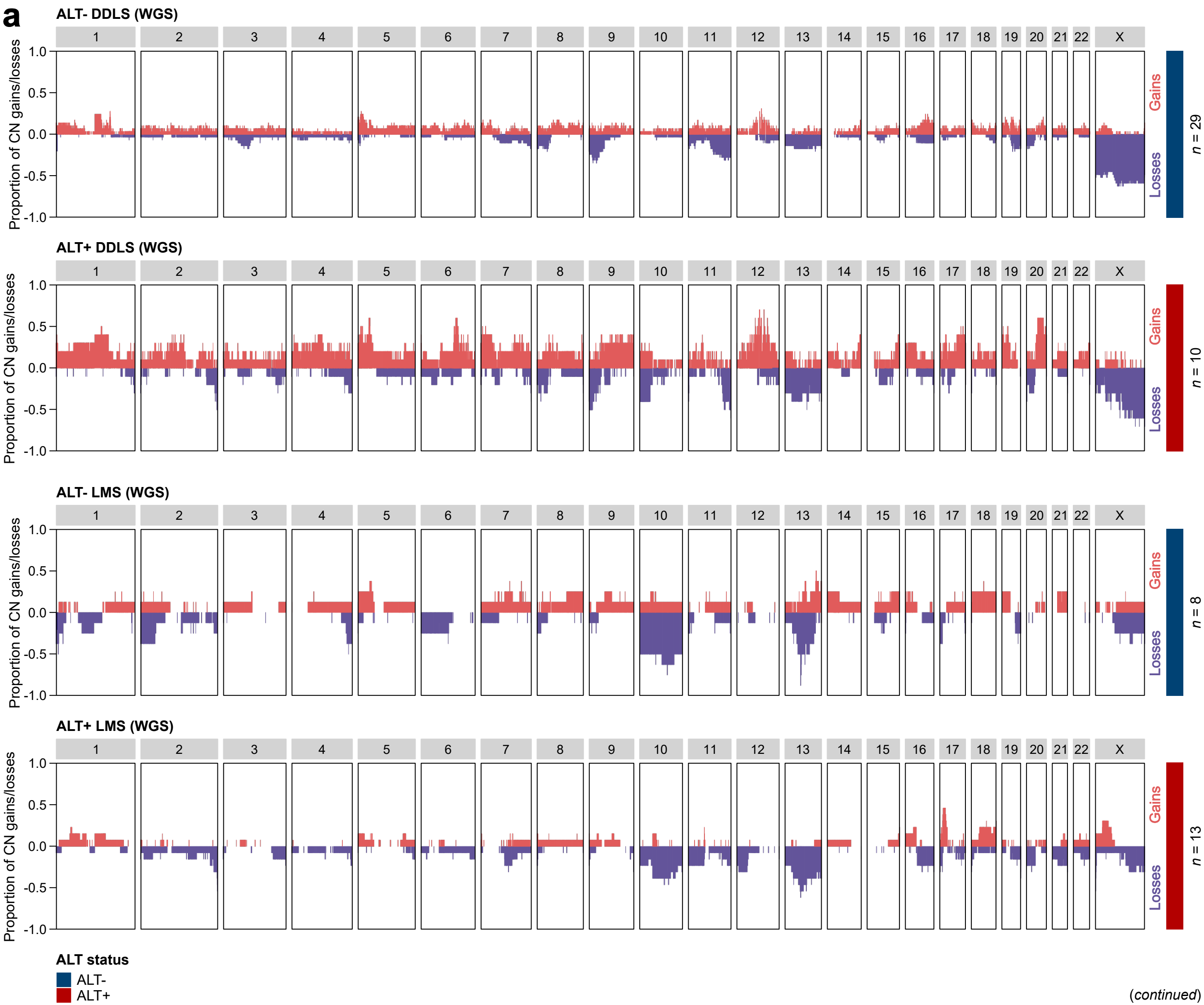


(*continued*)

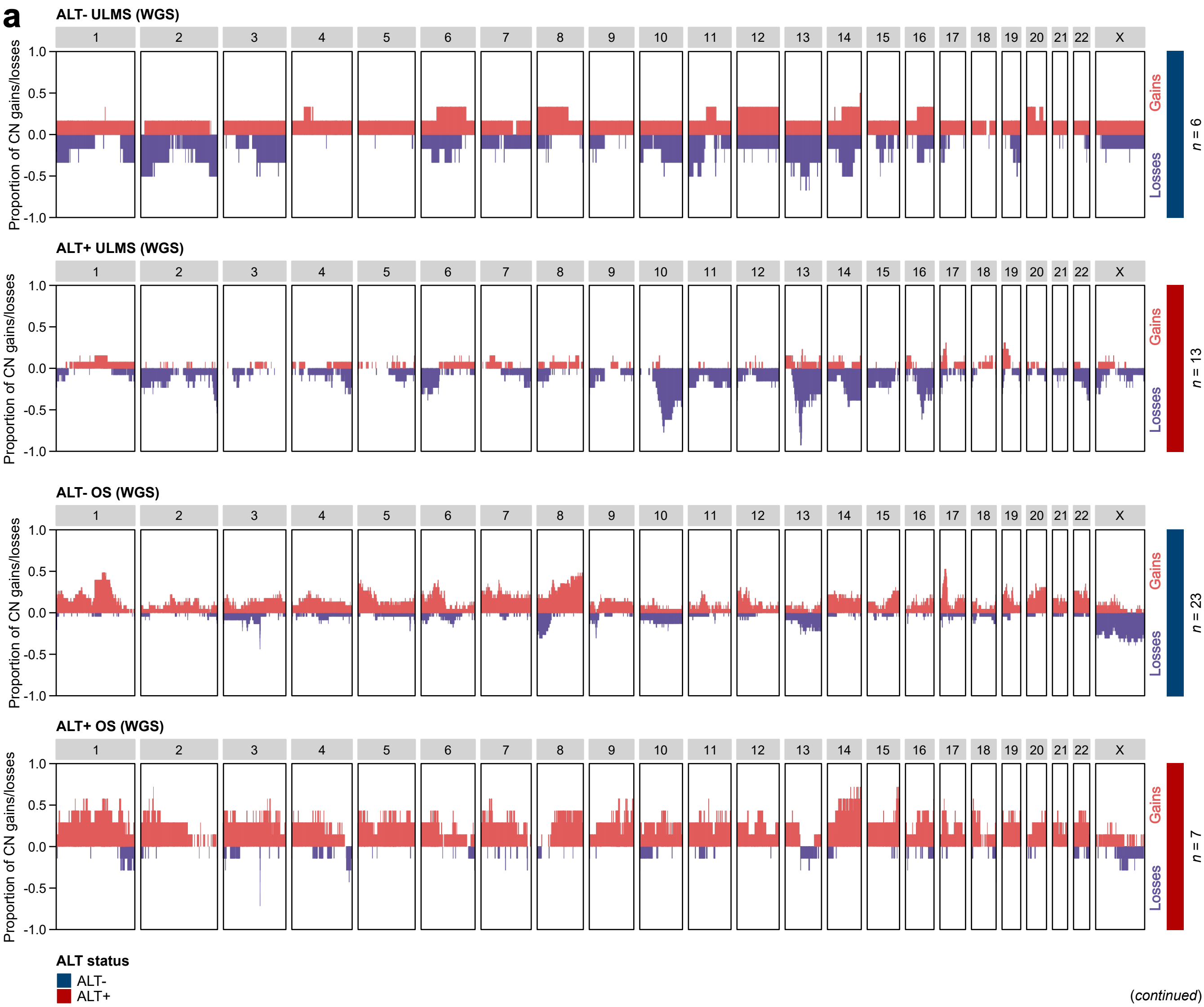
a
ALT- ULMS (WGS)
ALT+ ULMS (WGS)
ALT- OS (WGS)
ALT+ OS (WGS)
1 2 3 4 5 6 7 8 9 10 11 12 13 14 15 16 17 18 19 20 21 22 X
Proportion of CN gains/losses
1.0
0.5
0.0
-0.5
-1.0
Gains
Losses
n = 6
n = 13
n = 23
n = 7
ALT status
ALT-
ALT+

(*continued*)

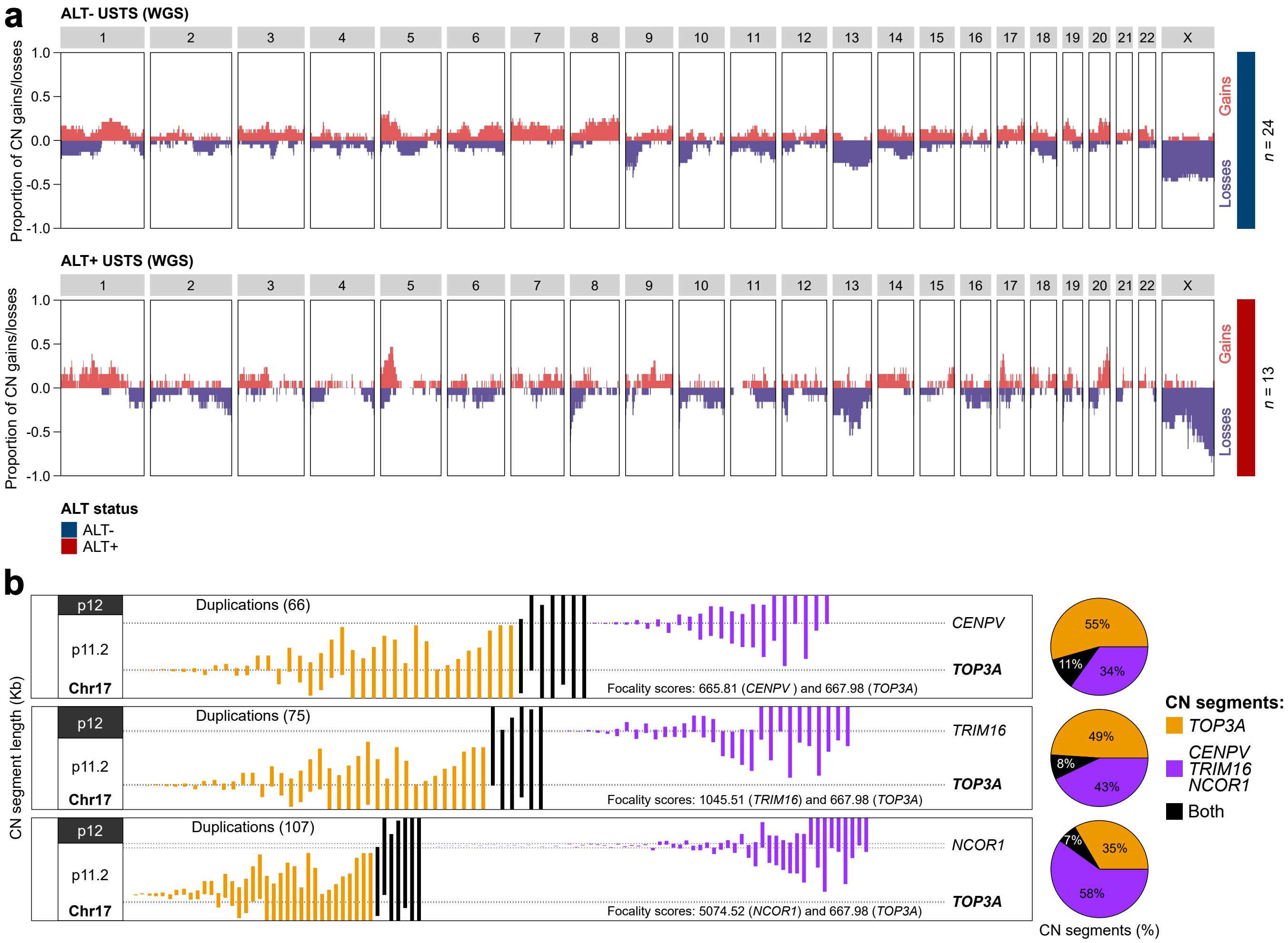
a
ALT- USTS (WGS)
ALT+ USTS (WGS)
1 2 3 4 5 6 7 8 9 10 11 12 13 14 15 16 17 18 19 20 21 22 X
Proportion of CN gains/losses
1.0
0.5
0.0
-0.5
-1.0
Gains
Losses
n = 24
n = 13
ALT status
ALT-
ALT+
b
CN segment length (Kb)
p12
p11.2
Chr17
Duplications (66)
Duplications (75)
Duplications (107)
CENPV
TRIM16
NCOR1
TOP3A
Focality scores: 665.81 (CENPV ) and 667.98 (TOP3A)
Focality scores: 1045.51 (TRIM16) and 667.98 (TOP3A)
Focality scores: 5074.52 (NCOR1) and 667.98 (TOP3A)
55%
11%
34%
49%
8%
43%
7%
35%
58%
CN segments (%)
CN segments:
TOP3A
CENPV
TRIM16
NCOR1
Both

### Supplementary Figure 6

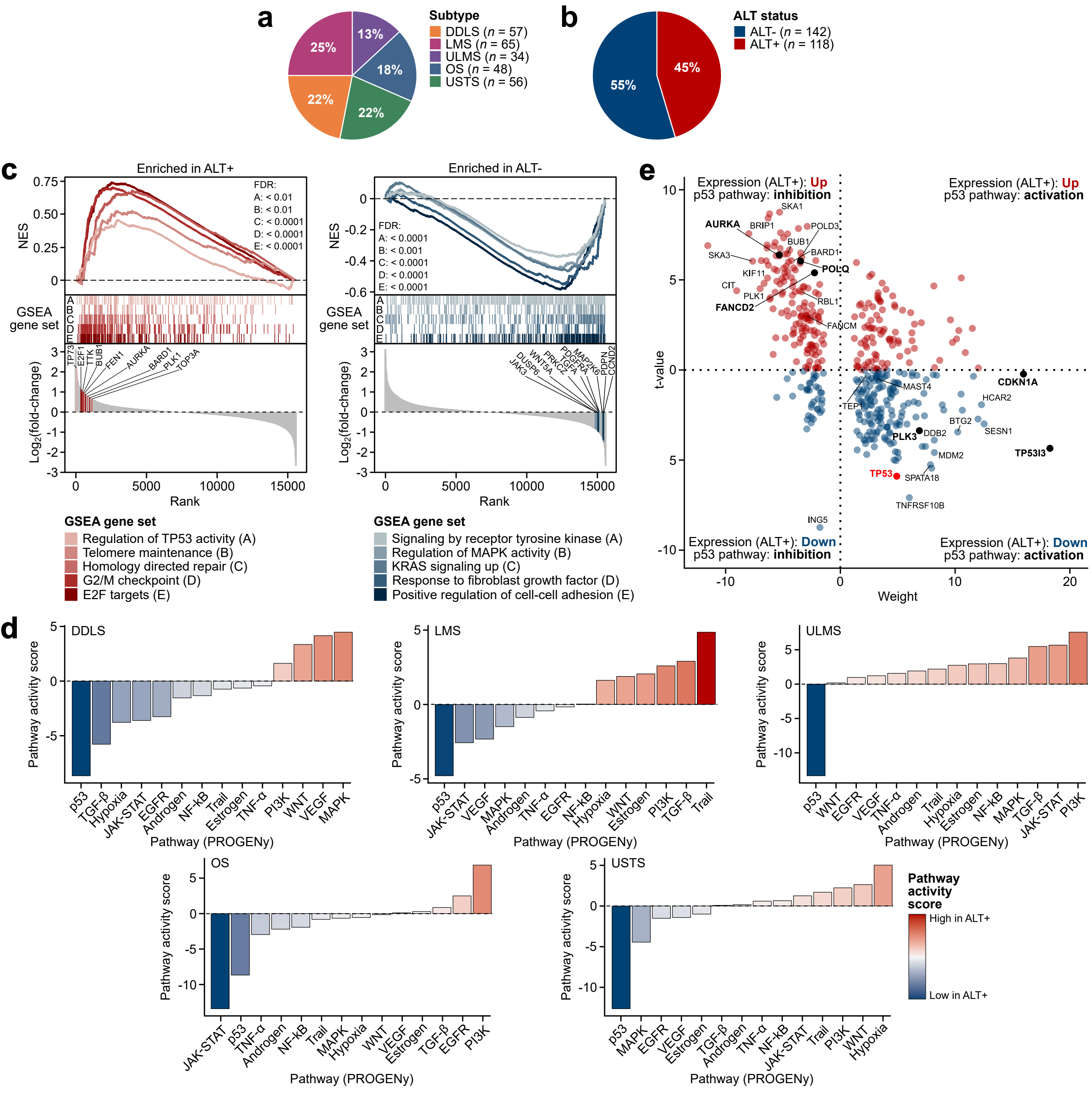

### Supplementary Figure 7

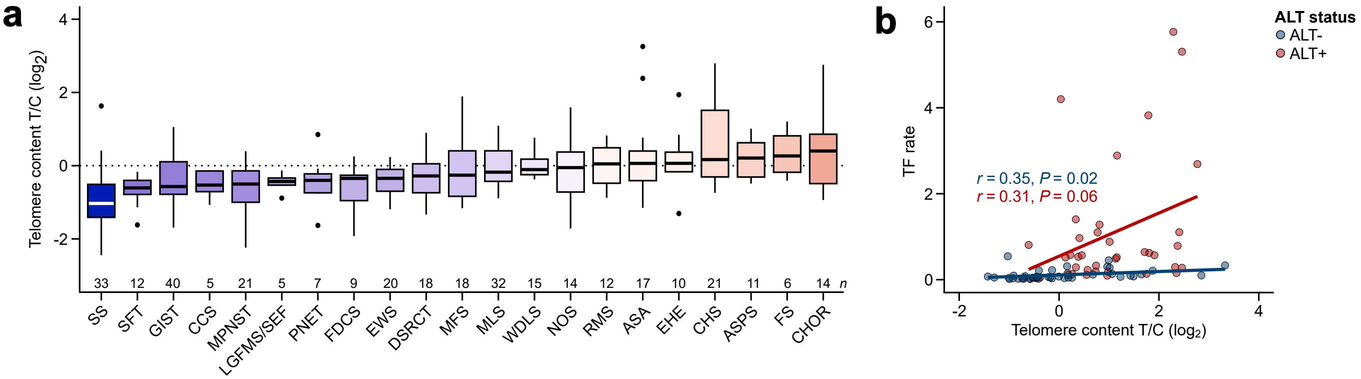

### Supplementary Figure 8

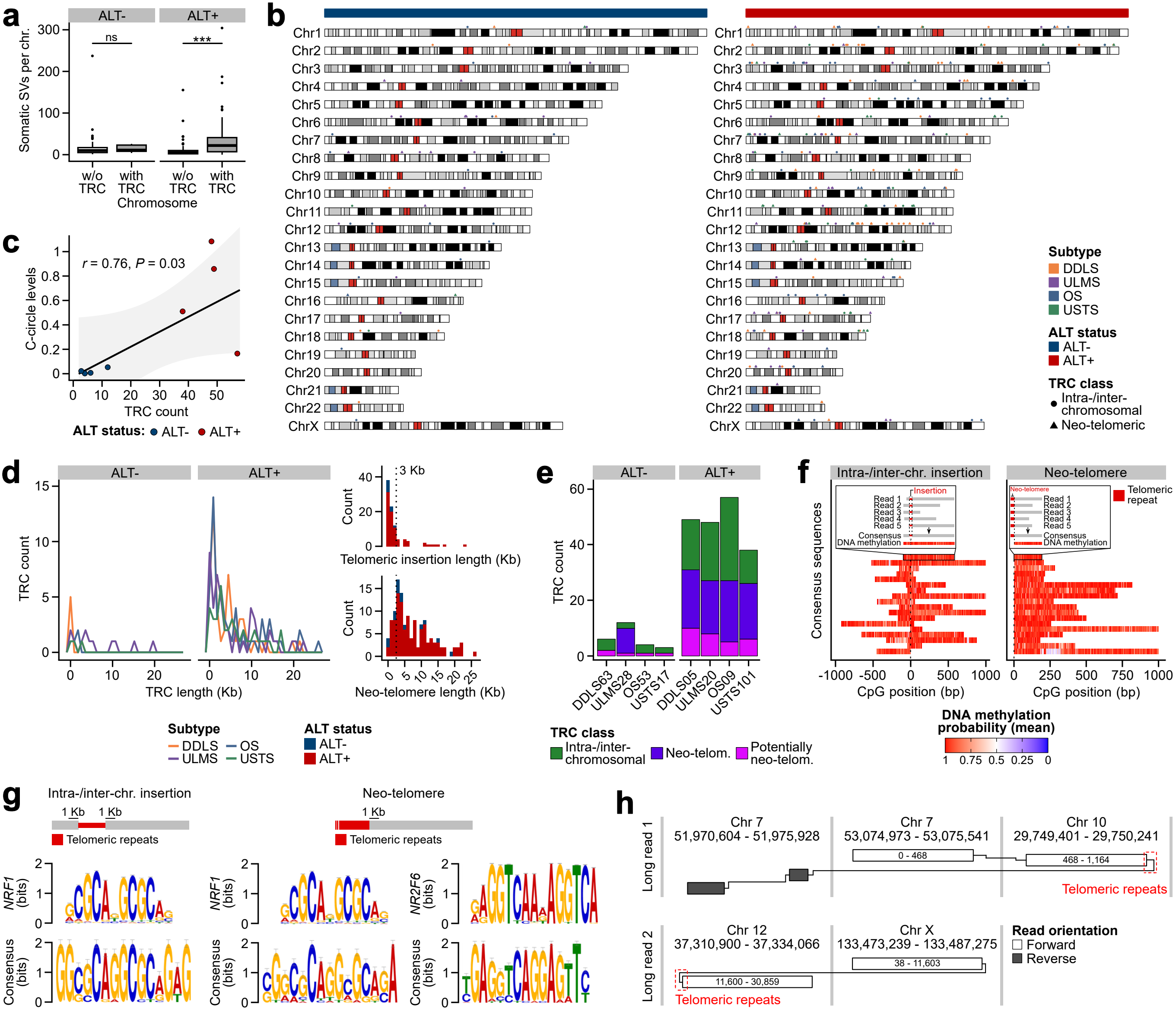

### Supplementary Figure 9

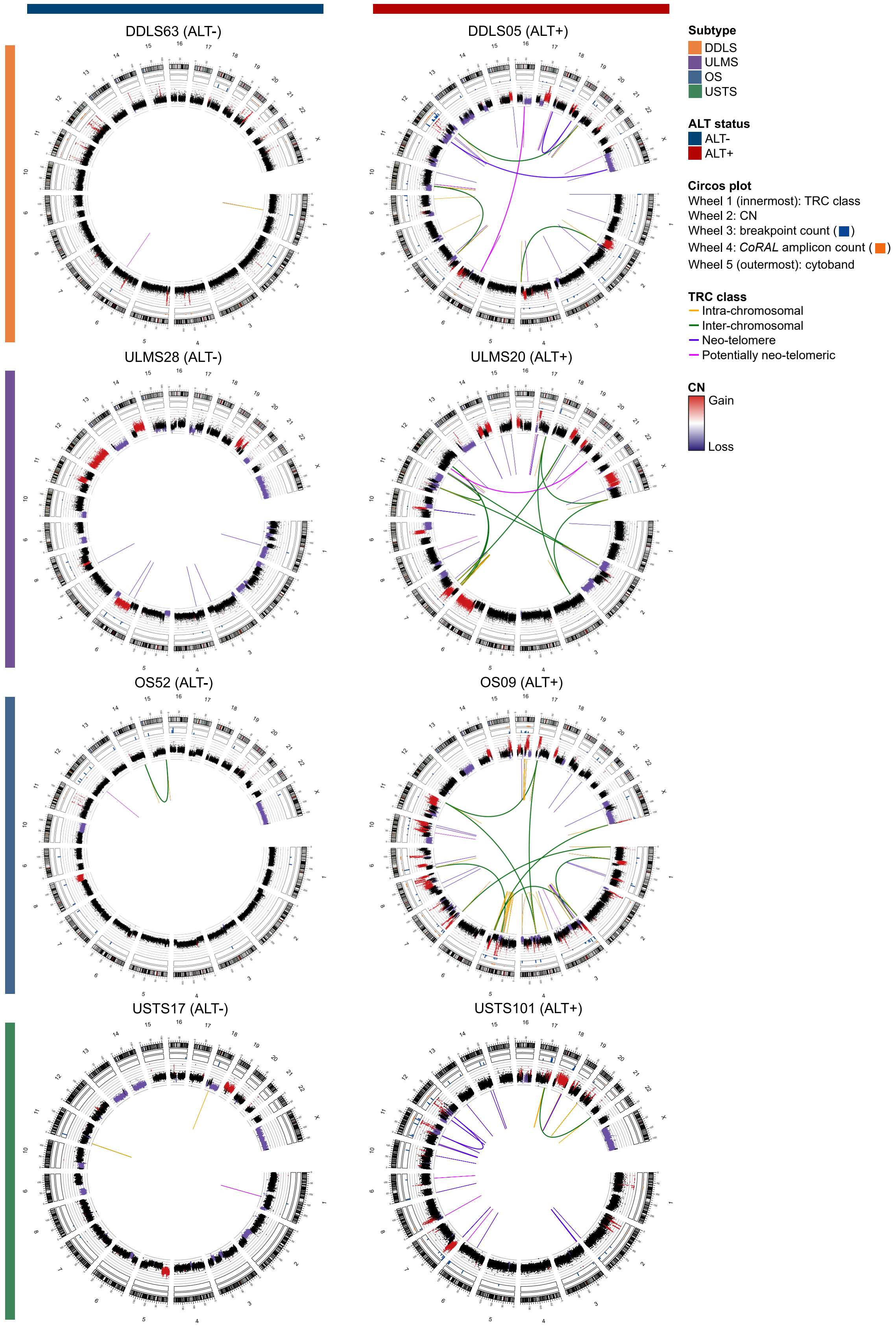

### Supplementary Figure 10

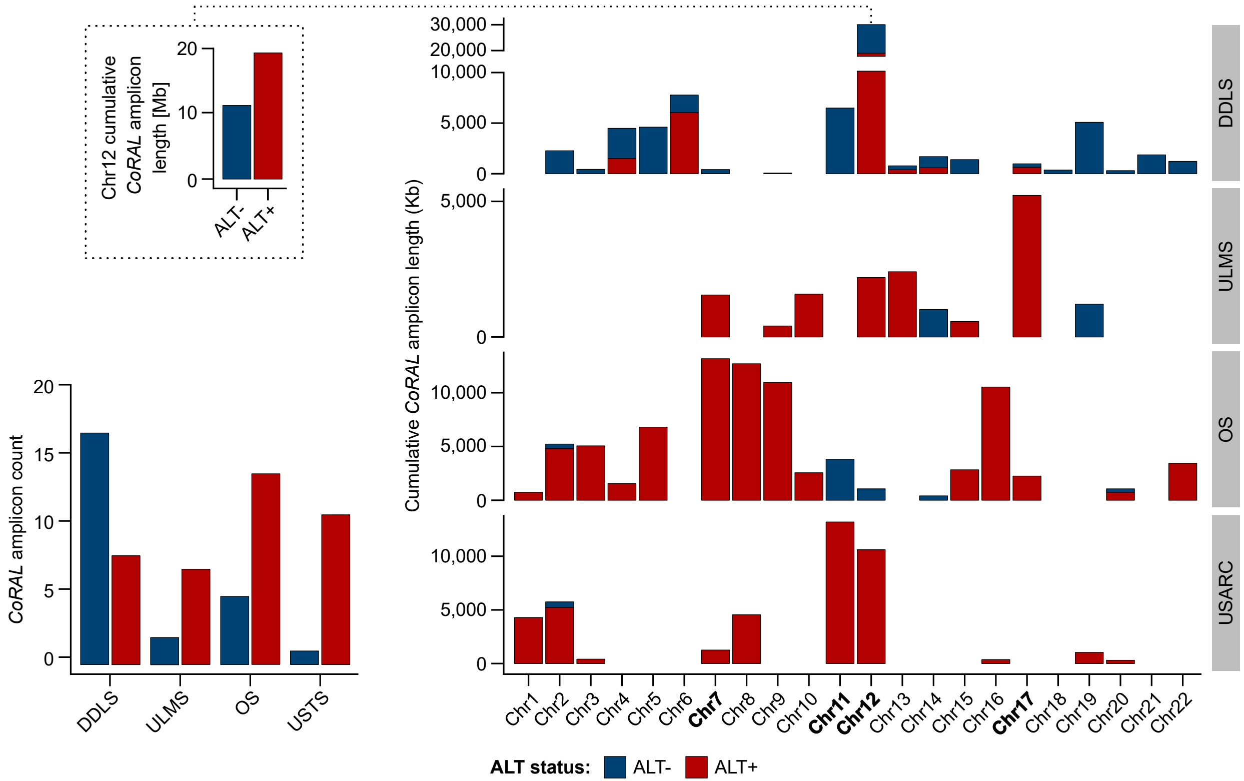

### Supplementary Figure 11

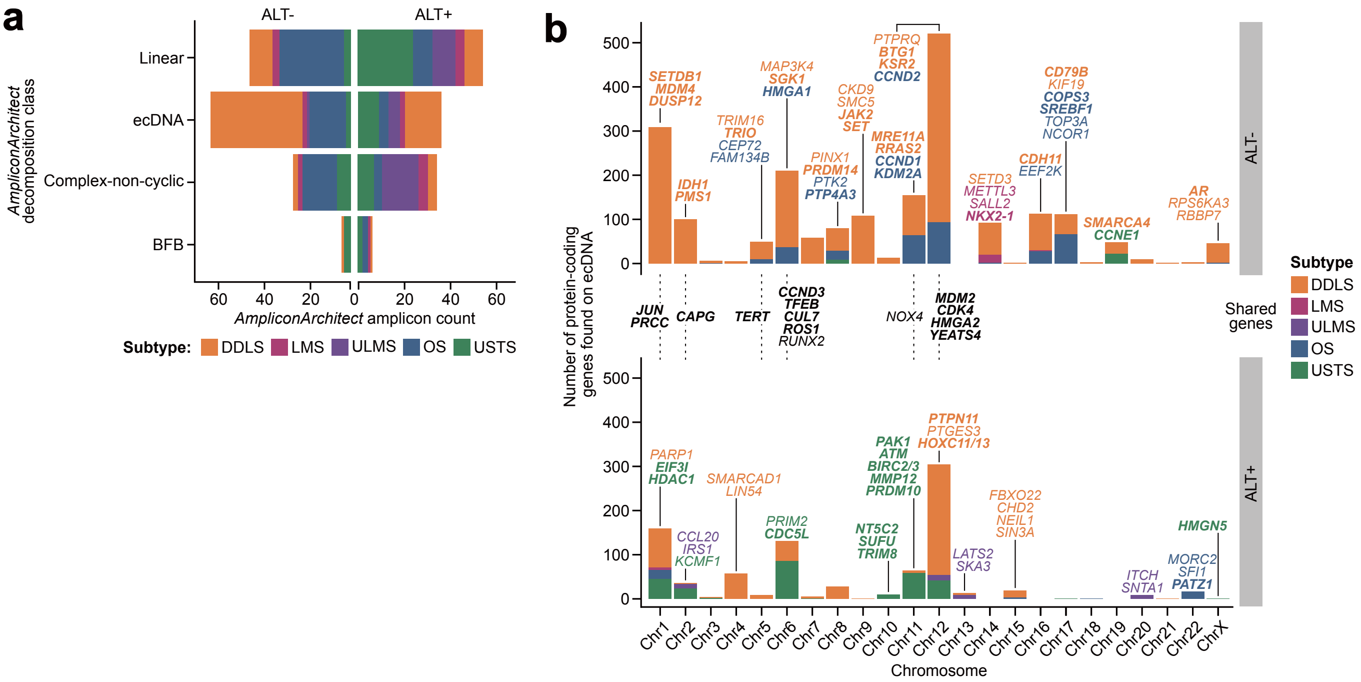
